# Biosynthesis of serotonergic Kanna alkaloids

**DOI:** 10.64898/2026.11.13.766392

**Authors:** Colin Y. Kim, Ryan S. Nett

**Affiliations:** Department of Molecular and Cellular Biology, Harvard University, Cambridge, MA, 02138, USA

## Abstract

Plants have repeatedly evolved to synthesize natural products that exploit serotonin signaling in animals. Although serotonin and most of its neuroactive analogs derive from tryptophan, the medicinal Kanna plant (*Mesembryanthemum tortuosum*) instead uses tyrosine and phenylalanine to construct a serotonin-like scaffold through unknown chemical transformations. Here, we elucidate the biosynthesis of Kanna alkaloids, including the serotonin reuptake inhibitor mesembrine. We reveal an unusual scaffold-forming strategy that is mediated by non-canonical transferase and oxidase enzymes, and we provide mechanistic insight on how an activated biosynthetic intermediate undergoes a multistep cyclization cascade to form the Kanna alkaloid scaffold. Collectively, our work uncovers how plants repurpose unconventional metabolic building blocks and enzymes to produce serotonergic molecules with promise as therapeutics for neurological disorders.

## Main Text

Serotonin is an ancient and widespread biological signaling molecule, used across kingdoms of life to connect an organism’s internal state to its environment (*1*). In humans, serotonin signaling plays a central role in mood regulation and is a principal target of modern antidepressants (*1–4*). Nature has repeatedly converged on the ability to produce serotonin mimics that can influence cognition (**Fig. 1A**), including psychedelics like *N*,*N*-dimethyltryptamine (DMT) from plants (*5*, *6*), psilocybin from fungi (*7*, *8*), and bufotenin from amphibians (*5*, *9*). More broadly, the potent neuroactive properties of serotonin mimics have led to their use as both medicines and critical research tools for understanding animal nervous systems (*10–13*).

**Fig. 1.**
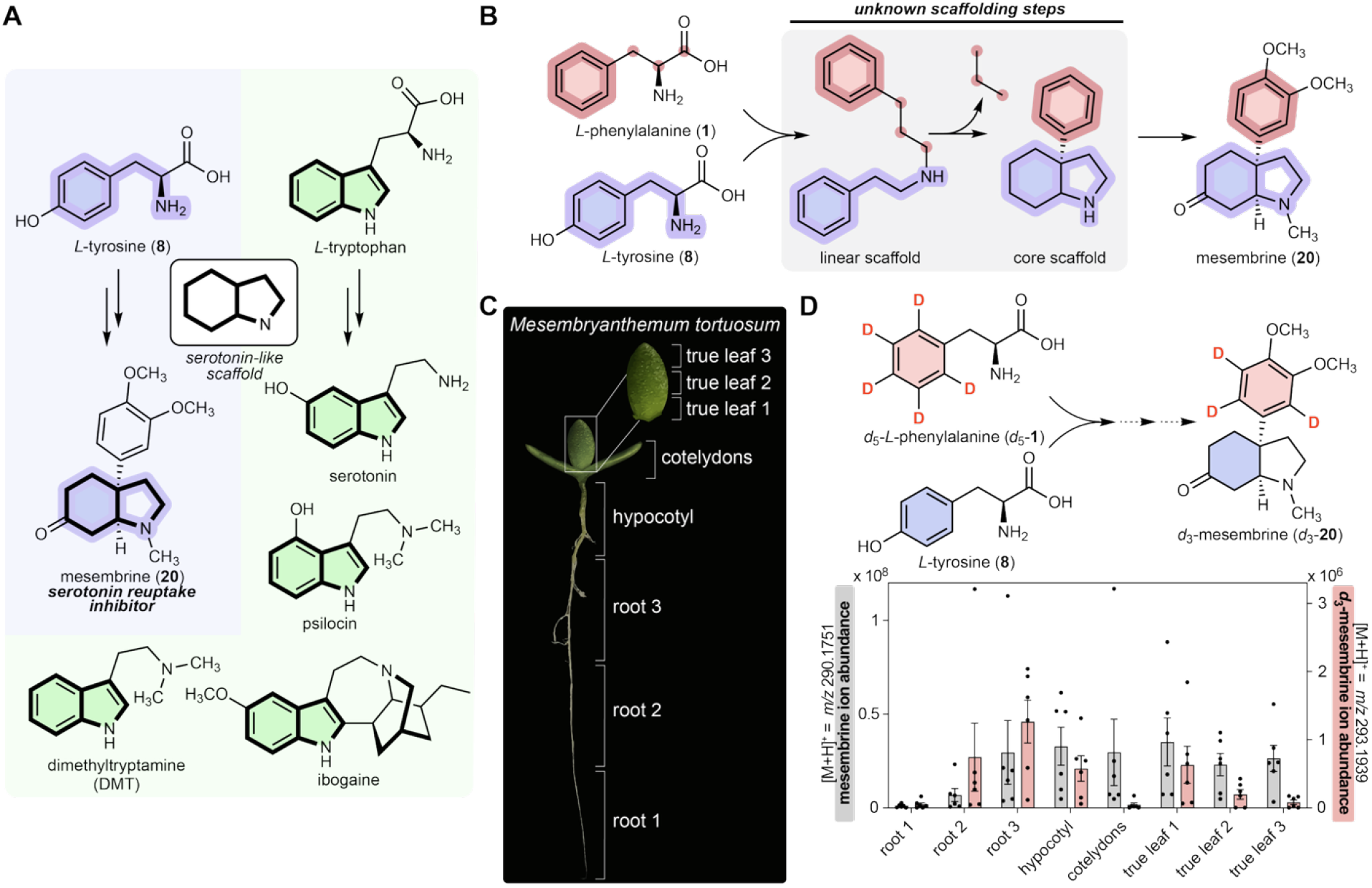
The chemical origins of serotonergic Kanna alkaloids. (**A**) Serotonin mimics found in nature. The serotonin-like scaffold of mesembrine (**20**) is derived from *L*-tyrosine, whereas most serotonergic natural products are derived from *L*-tryptophan. (**B**) Proposed biosynthetic pathway of mesembrine (**20**), including the unknown scaffolding steps highlighted as simplified carbon skeletons. (**C**) Image of 4-week-old *M. tortuosum* seedling with annotations of tissues harvested for metabolite profiling. (**D**) Isotope tracing experiment with *d_5_*-*L*-phenylalanine fed to *M. tortuosum* plant in (C). LC-MS profiling of unlabeled mesembrine (**20**) (gray, left axis) and *d_3_*-mesembrine (pink, right axis). Each bar graph represents the mean of six biological replicates across eight tissues with error bars representing SEM.

Nearly every known serotonergic natural product is built from *L*-tryptophan, including serotonin itself. A notable exception is the neuroactive alkaloid mesembrine (**20**), a serotonin reuptake inhibitor that instead derives from *L*-tyrosine and *L*-phenylalanine (*14*, *15*) (**Fig. 1A**). Mesembrine (**20**) contains an octahydroindole scaffold with the same ring connectivity as serotonin, and is one of many alkaloids produced by the South African succulent *Mesembryanthemum tortuosum*, commonly referred to as Kanna (*16*, *17*). *M. tortuosum* has historically been used as a mood-altering herbal remedy, and recent studies have evaluated the Kanna alkaloid extract as a potential treatment for depression (*18–21*). The serotonergic activity and unusual scaffold of the Kanna alkaloids led us to investigate the natural biosynthesis of these molecules, including how *L*-tyrosine and *L*-phenylalanine are modified and ultimately combined to form the serotonin-like scaffold. Previous studies proposed that this scaffold is generated from the cyclization of a linear precursor (*22–24*), with ring closure requiring an oxidative carbon-carbon (C-C) coupling between its two aromatic rings (**Fig. 1B**). Although this oxidative coupling reaction is analogous to other cytochrome P450 (CYP)-catalyzed reactions in plants (*25–27*), mesembrine (**20**) biosynthesis is atypical in that three carbons are lost in an extensive rearrangement cascade following oxidative phenol coupling (**Fig. 1B**).

Here, we leverage single-nuclei RNA-sequencing (snRNA-seq) and genome assembly in *M. tortuosum* to elucidate mesembrine (**20**) biosynthesis. We resolve how its scaffold is generated by identifying two key enzymes: an acyltransferase with dual *N*-acylation and non-canonical hydration activity, and a CYP that catalyzes an oxidative C-C coupling, which initiates a structural rearrangement to form the Kanna alkaloid scaffold. Together, our work establishes a biochemical and genetic basis for how *M. tortuosum* has evolved an unconventional biosynthetic strategy to serotonergic natural products.

### Kanna alkaloids are actively synthesized in hypocotyl and root cells

We first mapped active biosynthesis of mesembrine (**20**) by feeding *d_5_*-*L*-phenylalanine (1) to *M. tortuosum* seedlings and measuring label incorporation into this alkaloid across different tissues with liquid chromatography-mass spectrometry (LC-MS). (**Fig. 1C, D**). Although unlabeled mesembrine (**20**) was present at similar levels in most tissues, incorporation of the *d_3_*-label into mesembrine (**20**) varied across the plant. The highest *d_3_*-label enrichment was observed at the junction of the root and hypocotyl (**Fig. 1D**), indicating that these tissues are biosynthetically active and likely expressing the mesembrine (**20**) biosynthetic genes. We observed a similar pattern of label incorporation into putative alkaloid precursors to mesembrine (**20**) (**fig. S1**), further supporting both a biosynthetic hypothesis to mesembrine (**20**) and the root-hypocotyl junction as an active location of Kanna alkaloid biosynthesis (**figs. S2-S3**).

### Single-cell gene expression analysis reveals a core Kanna alkaloid biosynthetic pathway

We extracted nuclei from the hypocotyl and root tissues of *M. tortuosum* seedlings and used snRNA-seq to assess gene expression at cellular resolution (**Fig. 2A, B**). In parallel, we generated a *de novo,* chromosome-scale genome assembly of *M. tortuosum* and obtained bulk tissue RNA sequencing data to generate high-quality references for downstream gene expression profiling and comparative analyses (**fig. S4**). We hypothesized that Kanna alkaloid biosynthetic genes would have enriched expression in the nuclei isolated from the hypocotyl-root junction, and that co-expression analyses with our snRNA-seq data using predicted on-pathway genes as bait could identify modules of gene candidates involved in mesembrine (**20**) production.

**Fig. 2.**
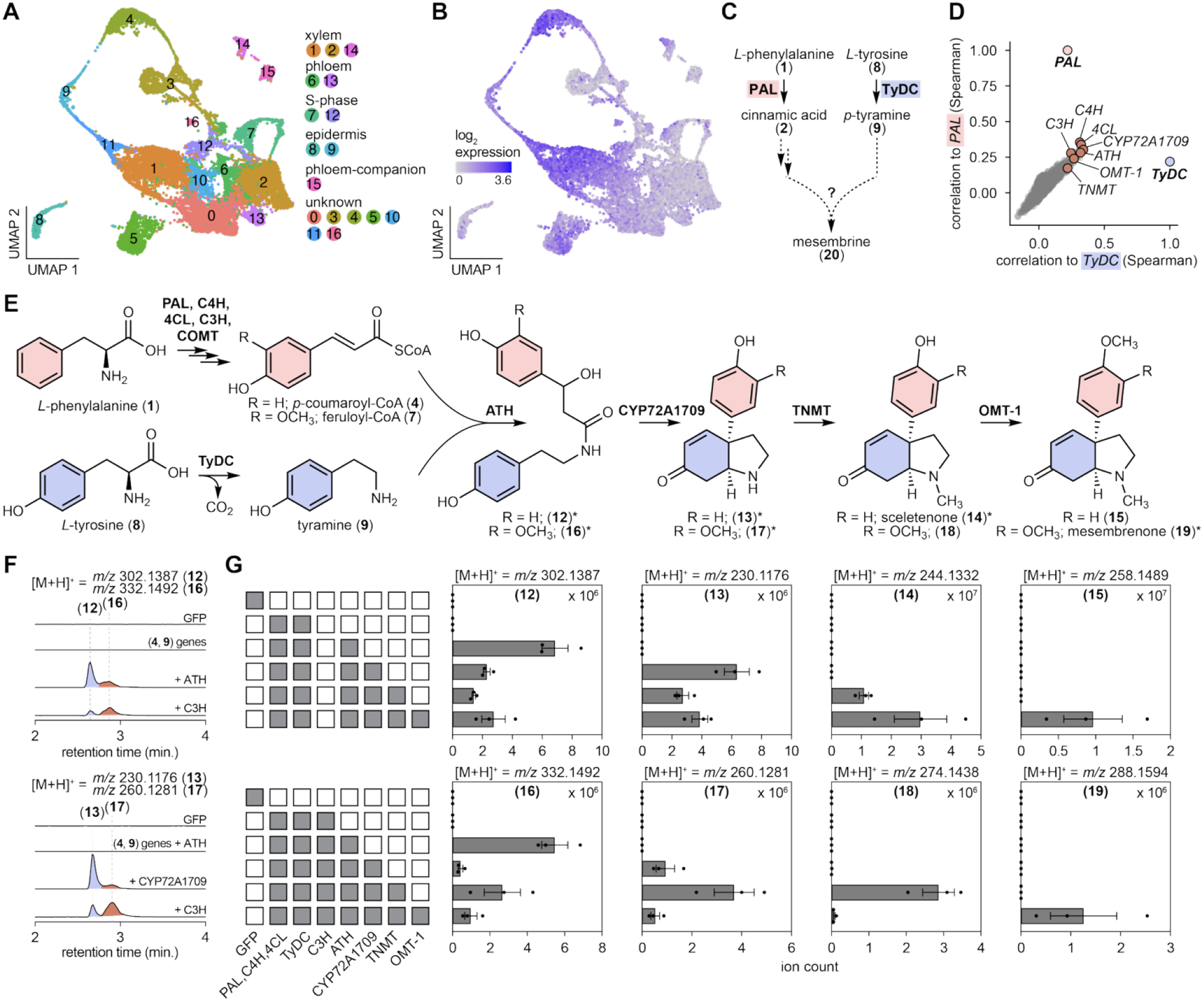
Discovery of a core Kanna alkaloid biosynthetic pathway. (**A**) UMAP clustering of single-nucleus RNA-seq (snRNAseq) profiles from biosynthetically active *M. tortuosum* tissue (15,629 nuclei, 17 clusters). (**B**) Expression of mesembrenone (**19**) biosynthetic genes (log_2_(count + 1)) in *M. tortuosum* nuclei (*PAL*, *C4H*, *4CL*, *TyDC*, *ATH*, *CYP72A1709*, *TNMT*, and *OMT-1*). (**C**) Reaction schematic of mesembrine (**20**) biosynthesis highlighting the proposed initial enzymes, phenylalanine ammonia lyase (PAL) and tyrosine decarboxylase (TyDC). (**D**) Plot showing the co-expression (via Spearman correlation coefficients) of *M. tortuosum* genes to *PAL* (pink) and *TyDC* (purple) using snRNAseq data. Orange dots indicate mesembrenone (**19**) biosynthetic genes. (**E**) Reconstituted biosynthesis of Kanna alkaloids using *N. benthamiana* transient expression. Asterisk (*) indicates structural validation via NMR or analytical standard (see figs. S19-S48). (**F**) Promiscuity of ATH and CYP72A1709 scaffolding enzymes, generating two distinct pathway outcomes to produce Kanna alkaloids. LC-MS extracted ion chromatograms (EICs) of **12** and **16** (top), **13** and **17** (bottom). GFP is shown as a negative control. (**4**, **9**) genes include *PAL*, *C4H*, *4CL* and *TyDC* with stepwise addition of ATH, CYP72A1709, and CYP98A286. We note that CYP98A286 exhibits pre-scaffolding hydroxylation activity (fig. S7). (**G**) Pathway buildup assays showing the mean of the peak area for EICs of each molecule.

Mesembrine (**20**) is derived from the amino acids *L*-phenylalanine (**1**) and *L*-tyrosine (**8**) (*14*, *23*, *24*). Biosynthesis could initiate through the activity of phenylalanine ammonia lyase (PAL) and tyrosine decarboxylase (TyDC), which would yield cinnamic acid (**2**) and *p*-tyramine (**9**), respectively. We anticipated that mesembrine (**20**) biosynthetic genes would co-express with *PAL* and *TyDC* genes, which are easily predicted by sequence homology (*28*, *29*). BLAST analysis of the *M. tortuosum* genome revealed single-copy *PAL* and *TyDC* orthologs that we then used as bait for gene co-expression analysis (**Fig. 2C**). Importantly, *PAL* and *TyDC* strongly co-expressed with each other, giving us confidence that their expected precursor-supplying activities are co-regulated in the cells that produce mesembrine (**20**) (**Fig. 2C-D, fig. S5**).

Guided by our biosynthetic proposal (**figs. S2-S3)**, we prioritized genes that both highly co-expressed with *PAL* and *TyDC* and had functional annotations (Pfam, BLASTp) that matched the predicted metabolic chemistry (**figs. S5**). To test biosynthetic gene candidates, we used *Agrobacterium*-mediated transient expression in *Nicotiana benthamiana* leaves, followed by LC-MS analysis. Using this approach, we found that expressing nine gene candidates together in *N. benthamiana* yielded mesembrenone (**19**), a direct precursor to mesembrine (**20**) (*22*, *23*) (**Fig. 2E-G**, **fig. S6**). In this batch of nine genes, five were essential for mesembrenone (**19**) production in *N. benthamiana*: *TyDC,* a BAHD acyltransferase (*ATH*), a CYP (*CYP72A1709*), an *O*-methyltransferase (*OMT-1*), and an *N*-methyltransferase (*TNMT*) (**fig. S6**). The other four genes were non-essential in our heterologous system but are homologous to phenylpropanoid metabolic genes that we expect to be on pathway: *PAL*, cinnamate-4-hydroxylase (*C4H*), 4-coumaroyl-CoA-ligase (*4CL*), and *p*-coumaroyl 3-hydroxylase (*C3H*). We suspect that endogenous phenylpropanoid metabolism in *N. benthamiana* made transient expression of these genes unnecessary, but we included them in subsequent experiments to increase precursor supply.

We then elucidated the function of each essential encoded enzyme to reveal a biosynthetic route for mesembrenone (**19**). First, we confirmed that TyDC produces *p*-tyramine (**9**) and we proposed that orthologous phenylpropanoid enzymes (PAL, C4H, 4CL) would generate *p*-coumaroyl-CoA (**4**) and feruloyl-CoA (**7**) in *N. benthamiana* (**fig. S6**). When ATH was added to this enzyme combination, we detected two major products by LC-MS (**12** and **16**). Purification and structural determination verified that **12** and **16** are linear amides generated from the *N*-acylation of **9** by either **4** or **7**, followed by subsequent alkene hydration (**figs. S19-S32**). The ATH products **12** and **16** are then consumed by CYP72A1709 to produce two new products (**Fig. 2F**) that we isolated and determined to be **13** and **17**, which each contain the characteristic Kanna alkaloid scaffold (**figs. S33-S46**). Thus, ATH and CYP72A1709 are responsible for catalyzing the key scaffolding reactions. Our data also suggest that these enzymes can accommodate different precursors to generate multiple scaffold-containing products in parallel, indicating more than one route to producing the Kanna alkaloids. In particular, the production of **12** and **16** by ATH indicates that either **4** or **7**–which only differ by the presence of a C3 methoxy–could act as co-substrates with **9** for scaffold formation. Indeed, adding the putative C3H from phenylpropanoid metabolism (CYP98A286), which would be on pathway to generate **7**, led to a 4.2-fold increase in **16** (**Fig. 2F, fig. S7**). Finally, combining TNMT and OMT-1 with the other identified enzymes led to the installation of *N*- and *O*-methyl groups, respectively, to generate mesembrenone (**19**) (**Fig. 2G, fig. S6**). Collectively, our data support a metabolic network towards Kanna alkaloid production that includes multiple entry points to the characteristic serotonin-like scaffold.

### Genomic analysis reveals a specialized NMT for Kanna alkaloids

Although we could reconstitute mesembrenone (**19**) biosynthesis in *N. benthamiana*, its yield was relatively low. In these experiments, we observed high levels of hordenine (**11**), a di-*N*-methylated derivative of *p*-tyramine (**9**) that we expected to be off-pathway (**fig. S8A-B**). Hordenine (**11**) was first identified in barley roots and is thought to inhibit growth of competing plants (*30–33*). Similarly, we found that hordenine (**11**) accumulates in the roots of *M. tortuosum* seedlings, unlike the Kanna alkaloids, which are enriched in the hypocotyl and leaves (**fig. S8C**). Consistent with hordenine (**11**) accumulation in the roots, *TNMT* expression is highest in this tissue (**Fig. 3C, fig. S8C**), suggesting that TNMT may be specialized for converting **9** into hordenine (**11**). Although TNMT can also *N*-methylate Kanna alkaloids, we hypothesized that another NMT gene may be involved in mesembrenone (**19**) biosynthesis (**fig. S8**).

**Fig. 3.**
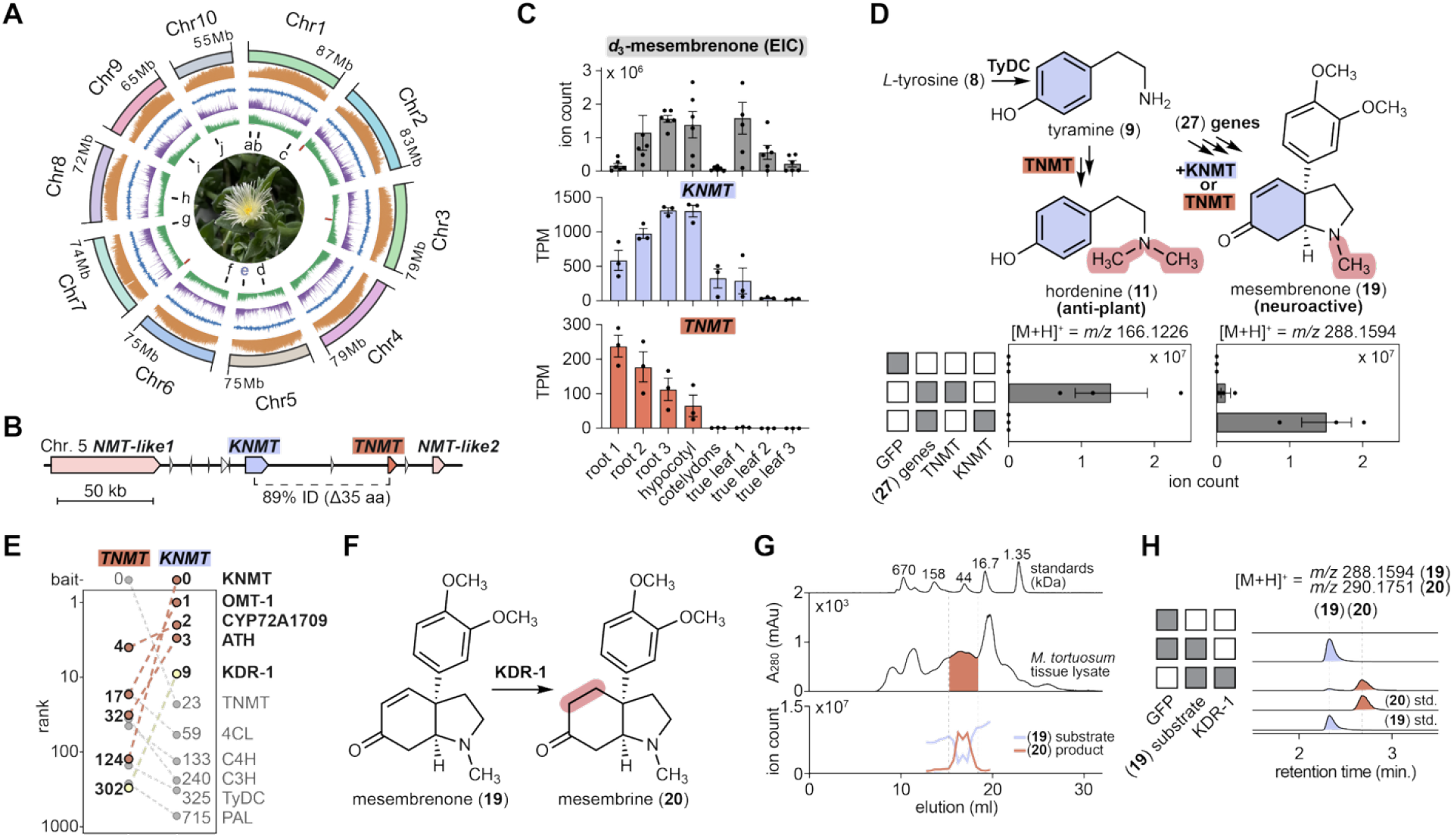
Genomic partitioning of *N-*methyltransferase genes guides discovery of mesembrine biosynthesis. (**A**) Circos plot summarizing genomic features for each pseudochromosome of *M. tortuosum*, including chromosome size, repeat element density (orange), GC content (blue), gene density (purple), transcript coverage (green), assembly gaps (red lines), and locations of mesembrine biosynthetic genes (from outer to inner tracks). a = *OMT-1* (*g272*); b = *KDR-1* (*g1529*); c = *ATH* (*g4212*); d = *CYP72A1709* (*g19370*); e = *KNMT* (*g20975*) and *TNMT* (*g20977*); f = *CYP98A286* (*g22226*); g = *C4H* (*g29247*); h = *PAL* (*g30732*); i = *TyDC* (*g35047*); j = *4CL* (*g37187*). (**B**) A locus of *N*-methyltransferase paralogs in the *M. tortuosum* genome. (**C**) LC-MS ion count from EIC of *d_3_-*mesembrenone from *d_5_*-*L*-phenylalanine feeding (top), and transcript expression of *KNMT* (middle) and *TNMT* (bottom) across *M. tortuosum* tissues shown in Fig. 1C. (**D**) Proposed pathway bifurcation of hordenine (**11**) and Kanna alkaloid biosynthesis. Pathway buildup assays in *N. benthamiana* showing the mean peak area for extracted ion chromatograms (EICs) of hordenine (**11**) and mesembrenone (**19**). (**E**) Comparison of snRNA-seq co-expression rank using *TNMT* (left) and *KNMT* (right) as bait genes. (**F**) Reaction schematic of KDR-1 activity. (**G**) NADPH-dependent activity-guided fractionation of the KDR-1 enzyme by size-exclusion chromatography of crude protein extract from *M. tortuosum*. A mixture of proteins with known size in kDa is injected as a standard. EICs of mesembrenone (**19**) and mesembrine (**20**) are shown. (**19** = *m/z* 288.1594; **20** = *m/z* 290.1751). (**H**) Discovery of KDR-1 converting **19** to **20**. Shown are LC-MS EICs for **19** and **20**.

By analyzing the *M. tortuosum* genome, we found that the *TNMT* gene locus on chromosome 5 had three paralogous NMT genes (**Fig. 3A-3B, fig. S8D-E**). While two paralogs were non-functional in our assays (**fig. S8F)**, we found that one paralog (Kanna alkaloid *N*-methyltransferase; KNMT) increased the production of mesembrenone (**19**) by 12.3-fold when used in place of TNMT (**fig. S8F-M**). Conversely, KNMT showed no ability to produce hordenine (**11**). Tissue-specific gene expression of *KNMT* matched the distribution of *d_3_-*mesembrenone in the isotope tracing experiment, further supporting that KNMT is specialized for Kanna alkaloid biosynthesis (**Fig. 3C**, **fig. S8G-M**).

### Discovery of a metabolic network to mesembrine

With the identification of KNMT, we could recapitulate mesembrenone (**19**) biosynthesis in *N. benthamiana* at 0.74 mg per gram dry leaf weight, which is within approximately 4-fold of the Kanna alkaloid levels in the native *M. tortuosum* plant (∼ 3 mg mesembrine (**20**)/g of leaf material) (*16*) (**fig. S8G**). To complete mesembrine (**20**) biosynthesis, we needed an enzyme capable of reducing the C-C double bond in the α,β-unsaturated ketone of mesembrenone (**19**). We initially prioritized oxidoreductase genes that strongly co-expressed with *PAL* and *TyDC* and shared less than 75% amino acid identity with their closest homologs from *M. crystallinum*, a close-relative to *M. tortuosum* that does not produce mesembrine (**20**). However, none of the candidates catalyzed the reduction of mesembrenone (**19**).

We then reassessed our gene co-expression analysis using *KNMT* as bait. In support of this approach, the top three correlated genes were *OMT-1*, *CYP72A1709*, and *ATH* (**Fig. 3E, fig. S9**). Additionally, the ninth ranked gene to *KNMT* was from the *NADH:flavin oxidoreductase* family (**Fig. 3E, fig. S9**), which would theoretically be capable of catalyzing the final reduction to produce mesembrine (**20**). In parallel to co-expression analysis, we used activity-guided fractionation followed by protein mass spectrometry (pMS) to isolate an enzyme fraction from *M. tortuosum* that catalyzes the NADPH-dependent reduction of mesembrenone (**19**) to mesembrine (**20**) (**Fig. 3F-G, fig. S10**). This approach led us to the same oxidoreductase enzyme candidate identified through co-expression to *KNMT*, supporting its potential role in Kanna alkaloid biosynthesis (**fig. S9-S10**). Indeed, this enzyme (Kanna alkaloid dehydrogenase/reductase-1; KDR-1) consumed mesembrenone (**19**) and produced mesembrine (**20**) during transient expression in *N. benthamiana* (**Fig. 3H**). Altogether, co-expression of KDR-1 with mesembrenone (**19**) pathway genes allowed us to reconstitute the total biosynthesis of mesembrine (**20**) in *N. benthamiana* (86 μg per gram dry weight) (**fig. S11**).

We initially overlooked KDR-1 due to its high amino-acid identity (87.3%) with an ortholog from *M. crystallinum* (**fig. S9**), as we had assumed that pathway-specific genes from *M. tortuosum* would have more divergent sequences. Unexpectedly, trace reductase activity was observed in KDR-1 orthologs from *M. crystallinum* and other plants that do not produce Kanna alkaloids (**fig. S12**). Thus, we postulate that KDR-1 activity was present in a promiscuous ancestral enzyme and that relatively few amino acid substitutions were sufficient to refine this enzyme to efficiently reduce mesembrenone (**19**) into mesembrine (**20**). This result highlights the potential pitfall of prioritizing biosynthetic candidates based on homology. Our data also reinforce how pathway assembly can incorporate multiple modes of enzyme evolution: modest substitutions that refine promiscuous enzymes such as in KDR-1 and KNMT, and neofunctionalization to yield specialized scaffolding enzymes like CYP72A1709 and ATH (*34*).

### ATH constructs a hydrated substrate primed for scaffold formation

With a biosynthetic pathway to mesembrine (**20**) elucidated, we next investigated how the serotonin-like scaffold is constructed by ATH and CYP72A1709. Similar to previous proposals (*35*), we hypothesized that scaffold formation required CYP72A1709-catalyzed phenol coupling of the linear amide precursors (**12** or **16**), followed by a retro-aldol rearrangement cascade that is facilitated by the β-hydroxy groups in these substrates. Considering this, we suspected that this β-hydroxy functional handle is essential for Kanna alkaloid biosynthesis.

Because we reconstructed mesembrine (**20**) biosynthesis in *N. benthamiana*, we initially hypothesized that ATH condensed *p*-coumaroyl-CoA (**4**) and *p*-tyramine (**9**) to produce *p*-coumaroyl-tyramine (**21**), which was then hydrated by an endogenous *N. benthamiana* enzyme to form **12**. However, we could not detect **21** in these experiments (**fig. S13A**). Alternatively, we considered that ATH may combine *p*-tyramine (**9**) with a hydrated CoA thioester (3-hydroxy-phloretoyl-CoA) that is generated from native *N. benthamiana* hydratase activity on *p*-coumaroyl-CoA (**4**). To resolve the uncertainty with this reaction, we assayed purified ATH with *p*-coumaric acid (**3**), *p*-tyramine (**9**), and purified 4-coumaroyl-CoA ligase (4CL) (**fig. S13B**). To our surprise, this assay directly produced **12** rather than **21**, indicating that ATH catalyzes both condensation of **4** and **9** and installation of the β-hydroxy group (**Figs. 4A, B, fig. S13C**).

**Fig. 4.**
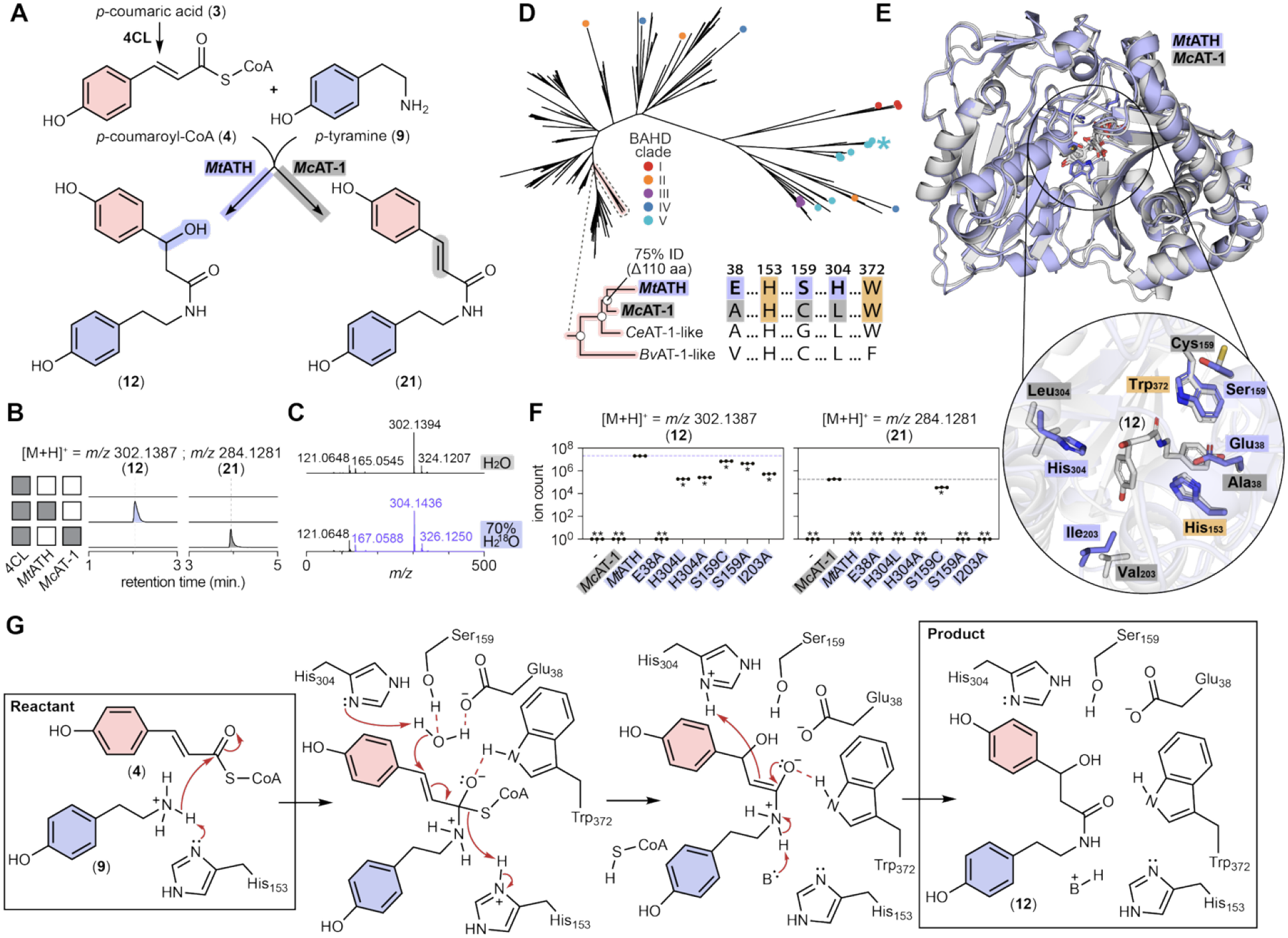
Dual *N*-acylation and hydration catalyzed by ATH. (**A**) Proposed reaction schemes for ATH and its canonical ortholog from *M. crystallinum*, *Mc*AT-1. (**B**) *In vitro* assays of ATH and *Mc*AT-1 with 4CL. LC-MS chromatogram traces showing **12** and **21**. (**C**) MS^1^ spectra of **12** for ATH *in vitro* assays conducted in H_2_O ([M+H]^+^ = *m/z* 302.1394) vs 70% H_2_^18^O ([M+H]^+^ = *m/z* 304.1436) showing that the ^18^O-label resides in the *L*-phenylalanine-derived fragment (*m/z* 167.0588). (**D**) Phylogenetic reconstruction from alignments of ATH and 182 orthologous proteins from the NCBI non-redundant protein database and genomes of species outlined in table S1. Previously characterized canonical BAHD enzymes belonging to clades I-V are shown. Cyan asterisk indicates the closest characterized homolog to ATH (*Os*THT) (*38*). White circles at the phylogenetic nodes indicate >90% bootstrap. Also shown is a protein sequence alignment of ATH and its closest orthologs highlighting key active-site residues. (**E**) Structural alignment of ATH and *Mc*AT-1 models with active-site residue differences around a **12**-docked pocket generated using Boltz-2. (**F**) Peak area of **12** (left) or **21** (right) detected from *in vitro* assays using ATH mutants compared to *Mc*AT-1. Negative control (-) is assay containing only 4CL. All assays were performed in three technical replicates, and the error bars represent SEM. Asterisk (*) indicates adjusted p < 0.0001 based on one-way ANOVA on log_10_-transformed peak areas. Double asterisks (**) indicates none detected. (**G**) Proposed mechanism of dual *N*-acylation and hydration catalyzed by ATH to conjugate and hydrate **4** and **9** to yield **12**, in which Glu38, Ser159 and His304 activate and position a catalytic water for addition across the α,β-unsaturated thioester intermediate.

This dual function has not been reported in the BAHD acyltransferase family, which is defined by a canonical acyl-transfer reaction mechanism (*36*, *37*). We found that ATH did not hydrate **21** when supplied directly *in vitro*, suggesting that hydration does not occur after release of the canonical *N*-acylation product (**fig. S13D**). Additionally, ATH did not hydrate **3** or **4**, indicating that hydration does not precede *N*-acyltransferase conjugation (**fig. S13E-G**). Moreover, *in vitro* ATH assays in 70% H_2_^18^O confirmed that water is the source of oxygen in the β-hydroxy group (**Fig. 4C, fig. S13H-K**). Together, these results suggest that *N*-acylation and hydration are coupled within a single ATH catalytic cycle.

We next investigated the evolutionary origin of the non-canonical, dual activity in ATH. Its closest characterized relative is a tyramine hydroxycinnamoyl transferase (*Os*THT; 22.2% amino acid identity) from rice (*Oryza sativa*) (*38*), but ATH belongs to a distinct clade and is most closely related to an ortholog from *M. crystallinum* (*Mc*AT-1) (**Fig. 4D**). Purified *Mc*AT-1 and 4CL demonstrated canonical activity, converting **3** and **9** to the canonical *N*-acylated product **21**, but not the hydrated scaffold **12** (**Fig. 4B, D**). Comparison of predicted structural models of ATH and *Mc*AT-1 identified several substitutions in ATH at active-site residues that are conserved in canonical acyltransferases (**Fig. 4D,E**). Mutating the ATH active-site residues to their counterparts in *Mc*AT-1 showed that E38A abolished activity, while H304L, H304A, S159C, S159A, and I203A significantly reduced production of **12** (**Fig. 4F**). Unexpectedly, S159C also restored formation of **21**, implicating S159 in positioning an active-site water to guide the reaction intermediate toward hydration (**Fig. 4F, fig. S14**). This single active-site residue switch suggests that the neofunctionalized activity in ATH arose through a relatively simple evolutionary change. Based on these results and the canonical hydratase enzyme mechanism (*39*, *40*), we propose that E38, S159, and H304 activate and coordinate catalytic water during the coupled *N*-acylation and hydration catalytic cycle, distinguishing ATH from its canonical acyltransferase relatives (**fig. S14**).

### CYP72A1709-initiated cascade yields the Kanna alkaloid scaffold

The distinctive scaffold-forming reaction in mesembrine (**20**) biosynthesis is initiated by CYP72A1709, which catalyzes formation of a C-C bond in **12** or **16** through oxidative phenol coupling **(Fig. 5A**). Critically, this generates an intermediate that is poised to spontaneously undergo a rearrangement cascade that involves breaking of a C-C bond through a retro-aldol fragmentation, hydrolysis of an amide bond to yield a three-carbon leaving group, and an aza-Michael addition to yield the bicyclic serotonin-like scaffold (*22*, *41*, *42*) **(Fig. 5A**). We confirmed CYP72A1709 activity in *N. benthamiana* with co-infiltrated substrates and through *in vitro* assays using CYP72A1709-enriched microsomes produced in yeast (*Saccharomyces cerevisiae*) (**Fig. 5B, fig. S15A-C**). Together, both assays support that CYP72A1709 is solely responsible for the conversion of the linear scaffolds **12** and **16** into the core scaffold-containing products **13** and **17**, respectively (**fig. S15C**).

**Fig. 5.**
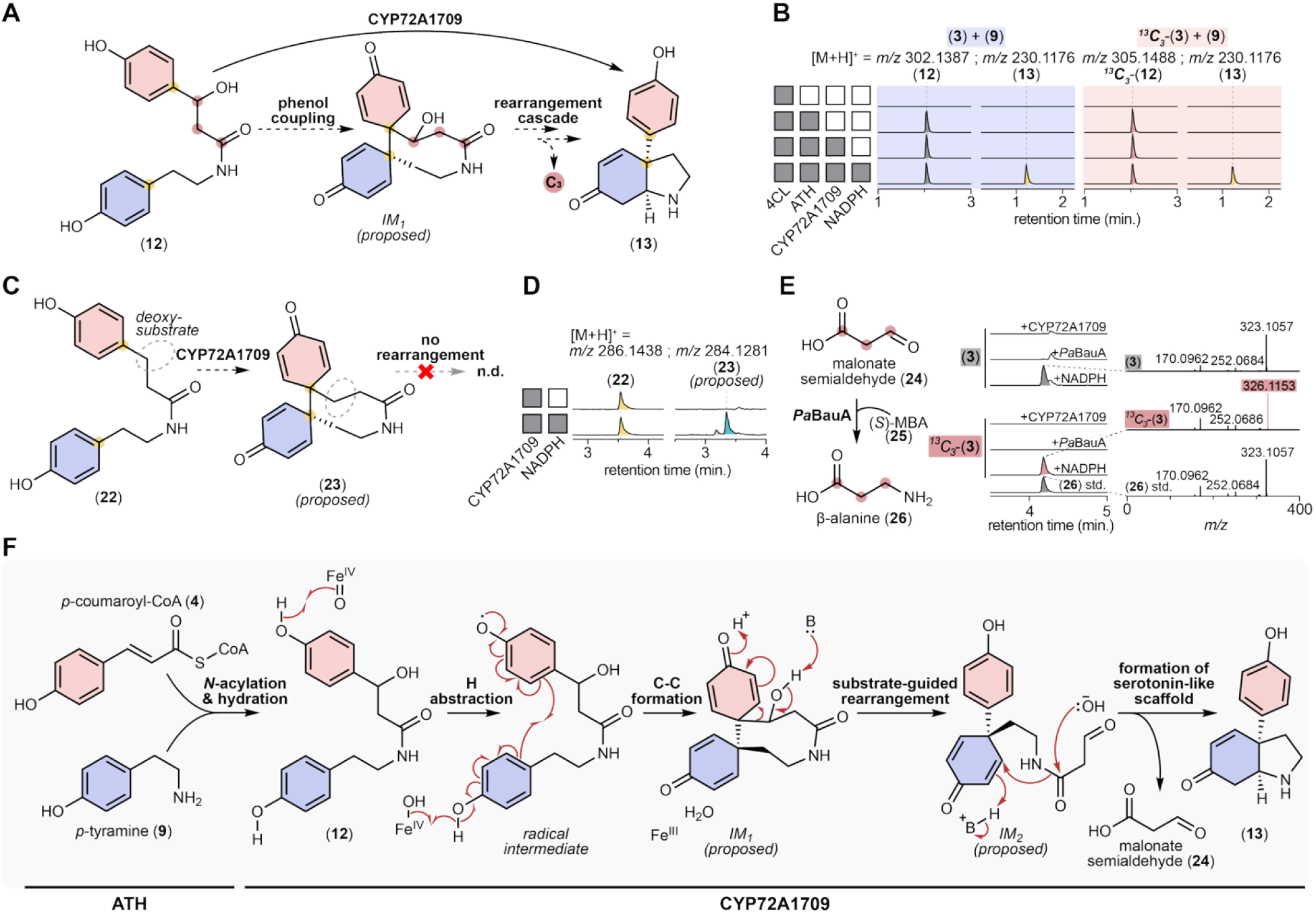
Mechanistic basis of serotonin-like scaffold formation in Kanna alkaloid biosynthesis. (**A**) Proposed reaction schematic of CYP72A1709 catalyzing C-C bond formation followed by a scaffolding cascade to convert **12** to **13**. (**B**) *In vitro* assays containing 4CL, ATH, CYP72A1709-enriched microsomes and NADPH with *p*-coumaric acid (**3**) and *p*-tyramine (**9**) (blue) or ^13^C_3_-*p*-coumaric acid (^13^C_3_-**3**) and **9** (pink). EICs of **12**, ^13^C_3_-**12**, and **13** are shown. **13** remains unlabeled, indicating that all three ^13^C atoms are lost with the byproduct. (**C**) Proposed reaction schematic of CYP72A1709 on the deoxy-substrate **22**, which lacks a β-hydroxy group. Phenol coupling yields an oxidized product ([M+H]^+^ = *m/z* 284.1282; proposed **23**) with no observed carbon loss. (**D**) LC-MS chromatograms of CYP72A1709-enriched microsome assay on **22**. EICs are shown for **22** ([M+H]^+^ = *m/z* 286.1438) and **23** ([M+H]^+^ = *m/z* 284.1281; proposed). (**E**) Malonate semialdehyde (**24**) released by CYP72A1709 is captured by *Pa*BauA using (*S*)-α-methylbenzylamine ((*S*)-MBA, **25**) as an amine donor to produce β-alanine (**26**). LC-MS EICs and MS/MS spectra from assays containing **3** (top) or ¹³C₃-**3** (bottom) with sequential addition of CYP72A1709, *Pa*BauA and NADPH to assays containing 4CL, ATH. **26** is detected via LC-MS after derivatization using dansyl chloride (dansylated **26** = *m/z* 323.1057 and dansylated ^13^C_3_-**26** = *m/z* 326.1153; both [M+H]^+^). Reactions are compared to a dansylated β-alanine (**26**) standard. (**F**) Proposed mechanism of serotonin-like scaffold generation catalyzed by ATH and CYP72A1709. ATH converts **4** and **9** via *N*-acylation and hydration mechanism as proposed in fig. S14. CYP72A1709 converts **12** to **13**, involving a Fe^IV^=O-mediated phenol radical coupling, formation of *IM_1_* and *IM_2_*, hydrolytic amide cleavage releasing malonate semialdehyde, and intramolecular aza-Michael cyclization.

Alkaloid ring systems are typically formed through condensation reactions that release simple byproducts like carbon dioxide, acetate, water, or ammonia (*41*, *43*). In contrast, the Kanna alkaloid scaffold-forming reaction is unusual, as it requires substantial rearrangement and carbon excision to release three carbons from the phenylalanine (**1**)-derived subunit (**Fig. 1B**, **Fig. 5A, fig. S15D**). We suspected that this rearrangement is mediated by the β-hydroxy group of the linear amide substrates (**12** or **16**) produced by ATH. Indeed, this β-hydroxy was essential for the rearrangement cascade, as CYP72A1709 assays with a linear substrate that lacks the β-hydroxy group (**22**) produced an oxidized product consistent with phenol coupling (*m/z* 284.1282; putative **23**), but without further carbon loss (**Fig. 5C, D**). Thus, we propose that CYP72A1709 catalyzes a conventional phenol coupling, and that the resulting intermediate is primed for a rearrangement cascade that requires the presence of the β-hydroxy group for the initial retro-aldol C-C bond cleavage to occur. Ultimately, this rearrangement leads to the excision of three carbons, thereby enabling an intramolecular aza-Michael addition to form the final serotonin-like scaffold (**Fig. 5A**).

We then assessed the three-carbon byproduct of the scaffold-forming reaction (**Fig. 5A**). An assay combining ^13^C_3_-*p*-coumaric acid (**^13^C_3_-3**) and **9** with recombinant 4CL, ATH, and CYP72A1709 corroborated that the three lost carbons derive from the aliphatic carbons of the phenylalanine-derived moiety in **12** (**Fig. 5B, fig. S15D, E**). Based upon our mechanistic proposal, we hypothesized that these three carbons were lost as malonate semialdehyde (**24**) (**Fig. 5E, F, fig. S16A**). Although we were unable to directly observe this byproduct, we verified the excised carbons to be malonate semialdehyde (**24**) by coupling the CYP72A1709 reaction with a bacterial β-alanine–pyruvate amino transferase (*Pa*BauA), which uses **24** as a co-substrate to produce β-alanine (**26**) (**Fig. 5E, fig. S16B**). Moreover, running the same assay in 70% H_2_^18^O produced a +2 *m/z* shift in **26**, indicating that the oxygen involved in hydrolytic cleavage comes from bulk water rather than molecular oxygen, the typical co-substrate of CYPs (**fig. S16C-G**). Altogether, our results reveal an unprecedented mechanism for the formation of serotonin-like scaffolds, wherein phenylalanine (**1**) and tyrosine (**8**) derivatives are coupled, rearranged, and cyclized through the tandem activity of two non-canonical enzymes (**Fig. 5F**).

## Discussion

Serotonergic natural products are typically derived from tryptophan, which already contains the indole serotonin scaffold. Here, we describe how a serotonin-like ring structure can be generated through extensive modification, coupling, and rearrangement of phenylalanine (**1**) and tyrosine (**8**). In addition to elucidating the biosynthesis of the serotonin reuptake inhibitor mesembrine (**20**), we reveal mechanisms by which two sequentially acting enzymes form its characteristic alkaloid scaffold and produce an unusual three-carbon byproduct. To our knowledge, malonate semialdehyde (**24**) has not been previously reported as a byproduct of scaffold formation in specialized metabolism. Instead, it is generally considered as a component of primary metabolism (*44*, *45*). When using *CYP72A1709* as a bait gene for co-expression analysis, we observe that the ninth highest ranked gene (*ALDH-like-1*) is annotated as *methylmalonate-semialdehyde dehydrogenase (acylating)*, a class of enzyme known to convert **24** into acetyl-CoA (*46–48*) (**fig. S17**). Although this remains speculative, the strong co-expression of these genes suggests that ALDH-like-1 may be coupled with CYP72A1709 to recycle the **24** byproduct back into primary metabolism (**fig. S17**), which would represent a unique case of carbon recovery in plant specialized metabolism.

Our work also reveals how tandem genomic duplication and neofunctionalization (*49*) of an *N*-methyltransferase gene can partition specialized metabolism to yield end-products with different biological activities. Although TNMT and KNMT share 89% amino acid identity, these enzymes have specialized catalytic roles for producing distinct, tyrosine-derived products: TNMT for synthesizing the anti-plant molecule hordenine (**11**), and KNMT for building the neuroactive Kanna alkaloids. In addition to their functional divergence, each of these genes are expressed differentially across *M. tortuosum* tissues. Specifically, *TNMT* expression is highest in roots where hordenine (**11**) accumulates, whereas *KNMT* expression is enriched in the hypocotyl zone where Kanna alkaloid biosynthesis occurs (**Fig. 3C, figs. S8, S9**). Together, the distinct enzyme functions and expression patterns of *TNMT* and *KNMT* provide a mechanism for controlling how amino acid precursors can be partitioned between the production of a molecule that inhibits the growth of neighboring plants or a complex alkaloid that can perturb animal serotonin signaling.

Single-cell co-expression analysis has emerged as a powerful strategy for discovering plant biosynthetic genes (*50–54*). In our work, this approach enabled the rapid identification of a Kanna alkaloid biosynthetic pathway, despite limited knowledge of the enzyme-mediated chemistry responsible for scaffold formation. The utility of snRNA-seq co-expression analysis is exemplified by comparing this approach to an equivalent analysis with bulk tissue RNA-seq data. Using *KNMT* as a bait, all pathway-specific genes ranked among the top 10 co-expressed in snRNA-seq-based analysis, whereas they ranked within the top 2,000 co-expressing to *KNMT* in bulk RNA-seq data (**fig. S11D**). By leveraging snRNA-seq, we reveal a minimal set of enzymes for reconstituting the production of mesembrine (**20**), including an elegant metabolic solution for building the serotonin-like scaffold from phenylalanine (**1**)-and tyrosine (**8**)-derived substrates. In addition to revealing an unconventional strategy for serotonergic scaffold formation, our work demonstrates how ATH and CYP72A1709 can accommodate different substrates to provide multiple entry points into the complex Kanna alkaloid biosynthetic network (**fig. S18**). Thus, beyond discovering and engineering a biosynthetic route to mesembrine (**20**), our results lay the groundwork for future efforts to engineer diverse serotonergic Kanna alkaloids that are relevant for both plant biology and human health.

## Acknowledgements

We thank Prof. Matias Kirst (University of Florida) and Dr. Noé Perron (Dana-Farber Cancer Institute) for sharing the *M. crystallinum* genome and snRNA-seq datasets. We also acknowledge Prof. David Nelson (University of Tennessee) for assisting with cytochrome P450 nomenclature. We thank Prof. Sang Gyu Kim (KAIST) and Conor McClune (Stanford University) for helpful discussions about snRNA-seq analysis. We are grateful to the Harvard Bauer Core Facility for guidance on single nuclei RNA-sequencing and the CCB Nuclear Magnetic Resonance Core Facility for NMR analysis. We acknowledge Prof. Jing-Ke Weng (Northeastern University) and Profs. Nicholas Bellono, Rachelle Gaudet, and Emily Balskus at Harvard University for providing critical feedback on this manuscript.

## Author contributions

Conceptualization: CYK, RSN

Methodology: CYK, RSN

Investigation: CYK, RSN

Visualization: CYK

Funding acquisition: CYK, RSN

Project administration: CYK, RSN

Supervision: RSN

Writing – original draft: CYK

Writing – review & editing: CYK, RSN

## Data, code, and materials availability

All data, code, and materials supporting the findings of this work will be made available upon request, following publication. Code used in this study is provided as a Supplementary Material. Raw sequencing data from *M. tortuosum* (short read, long read, snRNAseq, genome sequencing) will be deposited to the National Center for Biotechnology Information (NCBI) database and made available upon publication. Coding sequences for all genes functionally characterized in this study from *M. tortuosum* will be deposited to GenBank and made available upon publication.

## Supplementary Materials

## Materials and Methods

Figs. S1 to S48

Table S1

Supplementary Methods

Supplementary Code Description

References (*55-102*)

