## Supplementary Information for "Biosynthesis of serotonergic Kanna alkaloids"

**The PDF file includes:**

Materials and Methods
Figs. S1 to S48
Table S1
Supplementary Methods
Supplementary Code Description
References (*55*-*102*)

**Other Supplementary Materials for this manuscript include the following:**

Supplementary Code

### Materials and Methods

Chemical reagents

Chemicals used in this study were obtained from Sigma-Aldrich, unless noted otherwise. All isotopically labeled compounds including $^{13}C_9$-*L*-phenylalanine (CLM-2250-H-PK), $d_5$-*L*-phenylalanine (DLM-1258-1), $^{13}C_3$-*p*-coumaric acid (CLM-10642-PK), and $H_2{}^{18}O$ (OLM-240-97-1) were obtained from Cambridge Isotope Laboratories Inc. Chemical standard of (–)-mesembrine was obtained from Cayman Chemicals (Item No. 21100) and (±)-mesembrenone was obtained from MedChemExpress (Cat. No. HY-124434A).

Plant growth

*Mesembryanthemum tortuosum* (formerly *Sceletium tortuosum*) seeds were obtained from oursacredgarden.org. Seeds were surface sterilized by 70% ethanol 10 min, 5% bleach 10 min, five rinses in sterile water and sown on ½ Murashige & Skoog medium containing 1.5% (w/v) sucrose and 0.9% (w/v) phytoblend agar at pH 5.8. Plates were maintained in a growth chamber at 30 ºC with a 16-hour light and 8-hour dark cycle without humidity control. Upon germination and growth for 4 weeks, seedlings were either used for metabolomics, isotope tracing experiments, nuclei isolation, or transplanted onto soil (Miracle-Gro Succulent Potting Mix) with fertilizer (Southern Ag PowerPak 20-20-20 Water Soluble Fertilizer with micronutrients) and grown for bulk protein extraction and for genomic DNA isolation.

*Nicotiana benthamiana* seeds were germinated on soil (Pro-Mix HP AGTIV Reach) with fertilizer (Southern Ag PowerPak 20-20-20 Water Soluble Fertilizer with micronutrients). Plants were in a growth chamber at 30 ºC with a 16-h and 8-h dark cycle without humidity control, but with routine watering to keep the soil damp. In general, 4-5-week-old plants were used for *Agrobacterium*-mediated transient expression.

Microbial growth

*Escherichia coli* strain NEB 10-beta (New England Biolabs) was used for plasmid cloning and propagation and *E. coli* strain BL21 (DE3) (New England Biolabs) was used for recombinant protein production. *E. coli* strains were routinely grown in LB or TB media with kanamycin (50 µg/mL) at 37 ºC. *Agrobacterium tumefaciens* strain GV3101 was used for transient expression in *N. benthamiana* and grown in LB medium with kanamycin (50 µg/mL) and gentamicin (30 µg/mL) at 30 ºC. *Saccharomyces cerevisiae* strain INVSc1 (Invitrogen) was used for CYP72A1709 microsome production and grown in SC-Leu medium at 30 ºC.

Stable-isotope precursor feeding in *M. tortuosum* seedlings

To map the active tissues of Kanna alkaloid biosynthesis, $d_5$-*L*-phenylalanine was fed to intact, *M. tortuosum* seedlings (**fig. S1**). Four-week-old *M. tortuosum* seedlings grown on ½ MS were transferred to 6-well plates and submerged in sterile water containing 1 mM $d_5$-*L*-phenylalanine. After incubation for 1 day in the growth chamber, seedlings were removed and washed three times in water to remove residual $d_5$-*L*-phenylalanine from the plant surface. Each seedling was then dissected into eight tissue sections (root 1, root 2, root 3, hypocotyl, cotyledons, true leaf 1, true leaf 2, true leaf 3; Fig. 1C) and flash-frozen in liquid nitrogen before metabolite extraction.

#### Metabolite extraction

Frozen *M. tortuosum* and *N. benthamiana* tissue were lyophilized and homogenized to a fine powder with 5 mm steel beads using a ball mill (Retsch MM 400; 25 Hz, 2 min). Cold extraction solvent (water containing 0.1% formic acid unless otherwise noted) was added at 100 µL per mg dry tissue, vortexed, and incubated for ≥ 30 min. Samples were centrifuged (10,000 × *g*, 5 min, 4 °C) and supernatants were diluted into solvent appropriate for the LC method (acetonitrile + 0.1% formic acid for HILIC; water + 0.1% formic acid for C18) such that the quantified signals did not saturate the MS detector, occasionally requiring more than one dilution from the same sample. The diluted samples were then filtered using Multiscreen® 96 well Plate, 0.45 µm PTFE membrane plates (Sigma-Aldrich) prior to LC–MS analysis.

#### LC-MS and MS/MS analysis

The specific parameters for LC-MS are listed in the **Supplementary Methods**. All analyses were performed on an Agilent 1290 Infinity II UHPLC coupled to an Agilent 6546 quadrupole time-of-flight (Q-TOF) mass spectrometer. Electrospray ionization (ESI) was performed in positive or negative ion modes. Metabolites were analyzed via HILIC or C18 chromatography using either an InfinityLab Poroshell 120 HILIC (2.1 × 150 mm, 1.9 µm) or a ZORBAX RRHD Eclipse Plus C18 (2.1 × 50 mm, 1.8 µm) column. For targeted MS/MS analyses, collision energies were optimized for each target molecule and typically ranged from 10-30 V.

To analyze and process the LC-MS data, Agilent MassHunter Qualitative Analysis was used. Extracted ion chromatograms (EICs) were generated using a ±20 ppm mass window unless otherwise specified. Peak areas were quantified using Extract Chromatograms function with manual baseline correction when needed. Compound identity was assigned by accurate mass within 20 ppm, retention-time match to an authentic or purified standard, and where indicated, MS/MS fragmentation.

#### Genome sequencing, assembly, and annotation of *M. tortuosum*

High molecular weight genomic DNA was isolated from *M. tortuosum* leaf tissues using the NucleoBond HMW DNA kit (TakaraBio) and quality-controlled via Femto Pulse (Agilent Technologies). HiFi SMRTbell libraries were prepared, size-selected to yield an average size of 18.5 kb with Short Read Eliminator (SRE), and sequenced on a PacBio Revio, producing 118.17 Gb of HiFi reads (~137-fold raw read coverage of the final 860.7 Mb assembly). In parallel, chromatin conformation capture libraries were prepared from *M. tortuosum* leaf tissues using the Dovetail Omni-C kit according to the manufacturer's protocol and sequenced on Illumina NovaSeq 6000 to obtain 40,168,039 read pairs.

The initial contig assembly of *M. tortuosum* was generated from HiFi reads with hifiasm (v0.24.0-r703)(*55*). Chromatin-contact scaffolding of the primary contig assembly was performed by Dovetail Genomics using the HiRise pipeline (*56*), yielding a final assembly of

860,735,218 bp across 2,280 sequences, comprising 10 chromosome-scale scaffolds (scaffold N50 = 74.7 Mb). Repeat-masking was performed using Earl Grey (v5.1.1) (*57*), a pipeline for transposable element annotation that uses RepeatModeler (v2.0.6), RepeatScout (v1.0.7), RepeatMasker (v4.1.5), and rmblast (v2.14.1+). Repeat analysis of the *M. tortuosum* genome revealed that 62.2% of the genome consisted of repetitive elements (fig. S4). The repeat-masked genome was annotated using the BRAKER3 pipeline (v3.0.8) (*58–60*), a combination of GeneMark-ETP, AUGUSTUS, and TSEBRA and a combination of transcriptome and proteome data to train the gene prediction tools. RNA-seq reads of *M. tortuosum* (24 libraries; generated as described below) were provided to BRAKER3 along with protein data from the Viridiplantae clade generated from OrthoDB (v11)(*61*). This resulted in annotation of 42,934 genes (57,591 transcripts). Genome assembly and its related annotation files were loaded onto JBrowse 2(*62*). Functional annotations were performed using the funannotate (v1.8.1) pipeline(*63*), incorporating eggNOG-mapper (*64*) and InterProScan/Pfam (*65*, *66*) domain assignment together with BLASTP(*67*) evidence against the Swiss-Prot, Araport11, and *M. crystallinum* proteomes(*68*, *69*).

Assembly completeness was assessed with BUSCO (v5.8.2) (*70*) and compleasm (v0.2.9) (*71*) against the eudicots_odb10 lineage datasets. Proteome completeness and consistency were evaluated using OMArk (v0.5.0) (*72*) against the Viridiplantae OMAmer database, yielding 96.0% conserved-ortholog (HOG) completeness and 89.5% proteome consistency.

Since there are various species within the *Mesembryanthemum* genus that phenotypically appear similar to *M. tortuosum*, we validated the species authenticity of the plant that was sequenced for genome assembly. Reference sets of Angiosperms353 marker genes from *Mesembryanthemum tortuosum* (https://treeoflife.kew.org/tree-of-life/5455; KBG414/93 (NBG)) and 5 other *Mesembryanthemum* species (*M. crystallinum*, *M. digitatum*, *M. prasinum*, *M. schlichtianum*, *M. splendens*) from the Kew Royal Botanic Gardens, Plant and Fungal Trees of Life (PAFTOL) project were used to query the assembled *M. tortuosum* coding sequences by BLAST. The genome showed highest mean pairwise identity to *M. tortuosum* (98.28%) of the 6 references.

RNA extraction, library preparation, sequencing and transcriptome assembly

*M. tortuosum* seedling tissues frozen at -80 ºC were homogenized to a fine powder under liquid nitrogen using a mortar and pestle that were pre-treated with RNAse Zap RNAse Decontamination Solution (Thermo Fisher Scientific). RNA was extracted using the RNeasy Plant Mini Kit (Qiagen), with on-column DNase digestion. RNA quantification for each sample was performed via Nanodrop at 260 nm, and quality was further assessed by measuring size distribution using an Agilent 2100 Bioanalyzer. This was performed for all tissue types (root 1, root 2, root 3, hypocotyl, cotyledons, true leaf 1, true leaf 2, true leaf 3; **Fig. 1C**) with three biological replicates. These samples were then used to perform short-read sequencing via library generation using NEBNext Ultra II with poly(A) selection and sequenced on an Illumina HiSeq platform to generate 150 bp paired-end reads (~20 million reads per sample). Upon sequencing, adapters were trimmed using Trimmomatic (*73*) (ILLUMINACLIP:adapters.fasta:2:30:10, SLIDINGWINDOW:4:5, LEADING:5, TRAILING:5, MINLEN:25), then pseudoaligned to either the annotated transcript set from the *M. tortuosum* genome or CD-HIT-collapsed Iso-Seq transcriptome using kallisto (v0.46.2) (*74*). For gene co-expression analysis, Pearson's *r* calculations were manually performed in Microsoft Excel using the $log_2$-transformed TPM estimations from kallisto across all samples.

For long-read sequencing, RNA was extracted from whole plant tissue of soil-grown *M. tortuosum* using the same approach as above and prepared into cDNA Iso-Seq library using the SMRTbell prep kit 3.0 (PacBio). This whole plant library sample was then sequenced on a PacBio Sequel IIe system (SMRT Cell 8M), yielding 4,773,258 HiFi reads. Upon sequencing, the recommended Iso-Seq pipeline (https://isoseq.how/) was used to process the data, including cDNA primer removal using lima, poly(A) tail and concatemer removal using isoseq refine, isoform-level clustering using isoseq cluster2. After clustering, 173,308 high-quality transcripts were obtained, which were collapsed at 99% sequence identity using CD-HIT-EST (v4.8.1) (*75*) to generate a non-redundant set of 70,211 transcripts. All scripts used for transcriptome assembly, quantification, and downstream analyses are available at **Supplementary Code**.

Preparation and sequencing of snRNA-seq of *M. tortuosum*

Nuclei were isolated from hypocotyl and root tissues of 4-week-old *M. tortuosum* seedlings. All extraction procedures were performed at 4 ºC and with wide-bore pipette tips. Tissues were physically disrupted by two double edge razor blades (Feather) on a glass petri dish for 2 mins in 300 µL nuclei isolation buffer (NIB). NIB consisted of 0.3 M sucrose, 1.25% Ficoll, 2.5% Dextran T40, 15 mM Tris-HCl pH 8, 20 mM MES, 10 mM MgCl2, 60 mM KCl, 15 mM NaCl, 0.5 mM spermine, 0.5 mM spermidine, and 0.1% Triton X-100, with 5 mM DTT, 1 mM PMSF, 1% Plant Protease Inhibitors (Sigma P9599), 0.8% BSA, and 0.2 U/µL Protector RNase Inhibitor (Roche) added immediately before use. The resulting solution containing chopped tissues was transferred to a pre-chilled 2 mL Dounce tissue grinder (Kimble, 885302), along with 400 µL of NIB used to wash and obtain the remaining tissues on the petri dish. Pestle A was used to further grind the samples by 10 up-down movements. The solution in the Dounce was incubated on ice to finish the lysis of the cells for 5 mins. 5 additional up-down movements with Pestle A were performed to release the rest of the nuclei. To remove debris, disrupted tissue solution was then passed through a pre-wet 20-µm cell strainer (CellTrics, 04-0042-2315) into a prechilled 1.5 mL Eppendorf with 500 µL of NIB to wash the remains of the Dounce tissue grinder. Nuclei were gently pelleted at 500 × *g* for 10 mins using a swinging bucket rotor. Supernatant was removed and nuclei were resuspended with 50 µL of nuclei resuspension buffer (NRB). NRB consisted of 0.3 M sucrose, 15 mM Tris-HCl pH 8, 60 mM KCl, 15 mM NaCl, 0.5 mM spermine, 0.5 mM spermidine, and 15 mM MES, with 5 mM DTT, 1% Plant Protease Inhibitors (Sigma P9599), 2% BSA, and 0.2u/µL Protector RNase Inhibitor (Roche) added immediately before use. After nuclei isolation, Acridine Orange (AO)/Propidium Iodide (PI) dye was added to stain the nuclei. An automated cell counter (LUNA-FX7™ Logos Biosystems) was used to count the number of nuclei and quality control for nuclei integrity before loading into the 10x Genomics Chromium controller. Libraries were generated using the Chromium GEM-X Single Cell 3' Reagent Kits v4. Libraries were then sequenced on the NovaSeq X Series (Illumina).

For data processing, adapter and poly(A) trimming were performed on cDNA reads using Cutadapt (v5.2) (*76*)(-q 30 3' quality trim, --trim-n, -g AAGCAGTGGTATCAACGCAGAGTACATGGG to remove the 10x TSO adapter, -a "A{20}" to remove poly(A), -m 30 minimum length). Bases with Phred quality score < 30 were trimmed and reads shorter than 30 nucleotides were discarded, then read pairs were synchronized using Seqkit (v2.4.0) (*77*). STARsolo (v2.7.11b) (*78*, *79*) was used to align the read pairs against the *M. tortuosum* genome. A STAR reference was generated using options --runMode genomeGenerate --sjdbOverhang 89 --genomeSAindexNbases 13, and read alignment was conducted using options --alignIntronMax 1 --alignSJDBoverhangMin 999 --soloBarcodeReadLength 0 --soloUMIlen 12 --soloStrand Forward --soloCellFilter EmptyDrops_CR --soloFeatures Gene GeneFull --soloMultiMappers EM --soloType CB_UMI_Simple. The resulting GeneFull count matrix, which includes intronic reads, was used for all downstream analyses. Downstream analyses were conducted using the R package Seurat (v5.5.1) (*80*, *81*). Ambient RNA contamination was removed using SoupX (v1.6.2) (*82*), then the data were normalized using the SCTransform method (*83*). Nuclei with fewer than 200 or greater than 5,400 detected features (nFeature_RNA) were filtered out. For further processing, the first 20 principal components were used for neighbor-graph construction and DoubletFinder (*84*) removed doublets in the Seurat object. 15,629 nuclei classified as singlets with a median of 3,386 UMIs and 2,151 genes detected per nucleus, were retained for all downstream analyses and resolved into 17 clusters. All scripts and parameters are provided to ensure reproducibility (**Supplementary Code**).

Because no cell-type reference exists for *M. tortuosum*, cluster identities were inferred by cross-species marker transfer from two independent reference atlases: the *M. crystallinum* leaf snRNA-seq atlas (*69*), comprising 86 literature-curated cell-type markers and the top 50 differentially expressed genes for each of its 17 clusters, and an Arabidopsis root/hypocotyl single-cell marker panel (*85–87*) comprising 44 literature-verified marker genes across 12 root-anatomy categories. Orthologs between *M. tortuosum* and each reference species were identified by reciprocal-best-hit DIAMOND (*88*) BLASTP (e-value < 1 × 10-6) and cross-validated against OrthoFinder (v2.5.5)-inferred orthogroups. Reference marker sets were mapped onto their *M. tortuosum* orthologs and scored per nucleus, then averaged per cluster, using UCell (v2.10.1) (*89*). Final cell-type calls were assigned from the convergence of enrichment significance, UCell score, and the functional annotation of each cluster's own top marker genes. Mapping statistics and scoring for each marker set transfer are summarized in **Supplementary Code**.

<u>*Agrobacterium*-mediated transient expression in *N. benthamiana*</u>

Biosynthetic gene candidates were first cloned into the pEAQ-HT vector. Gene fragments were obtained from PCR amplification using template cDNA libraries synthesized from RNA extracted from root 3 or hypocotyl tissues via SuperScript™ III First-Strand Synthesis Supermix (Invitrogen). In parallel, coding sequences from the annotated *M. tortuosum* genome were synthesized by Twist Bioscience with codon optimization for *N. benthamiana*. Gene fragments were assembled into the digested pEAQ-HT vector (AgeI/XhoI) using NEBuilder HiFi DNA Assembly Master Mix (New England Biolabs). The resulting constructs were then transformed into *E. coli* NEB 10-beta using the heat shock protocol provided by the manufacturer, plated on LB agar containing kanamycin (50 µg/mL), and incubated overnight at 37 ºC. After confirming gene insertion via colony PCR followed by agarose gel electrophoresis, positive colonies were grown in LB containing kanamycin overnight at 37 ºC for plasmid purification. Plasmid sequences were verified by whole-plasmid sequencing (Genewiz), transformed into *A. tumefaciens* GV3101 by freeze-thaw transformation, and selected on LB agar containing kanamycin and gentamicin (30 µg/mL) for 2 days at 30 ºC.

*A. tumefaciens* GV3101 strain carrying verified pEAQ-HT plasmid construct containing each biosynthetic gene candidate was grown overnight at 30 °C in LB with kanamycin, harvested by centrifugation and resuspended in induction medium (10 mM MES pH 5.6, 10 mM $MgCl_2$, 150 µM acetosyringone). Suspensions were incubated at room temperature for at least 1 hour prior to infiltration. Optical density at 600 nm ($OD_{600}$) was measured for each strain and appropriate strains were combined such that each strain's final $OD_{600}$ was 0.3. To express genes, strain mixtures were infiltrated into the abaxial side of 4-5 week old *N. benthamiana* leaves. Leaves were harvested 3-5 days post-infiltration for further processing. For substrate co-infiltrations, 100 µM substrate was dissolved in water and infiltrated a second time 3 days after the infiltration of the *A. tumefaciens* strains. Leaves were harvested 1 day post co-infiltration of substrates for further processing.

#### *In vitro* microsomal CYP72A1709 assays

*CYP72A1709* gene fragment from *M. tortuosum* was codon optimized for expression in *S. cerevisiae* and synthesized by Twist Bioscience. This gene fragment, along with its highest co-expressing cytochrome P450 reductase gene (*CPR-1*) were both cloned into the pESC-leu2d vector (yeast expression vector with a bidirectional promoter) digested with BamHI + HindIII and PacI + NotI restriction enzyme pairs. Sequence-verified plasmid construct was then transformed into *S. cerevisiae* strain INVSc1 (Invitrogen) using the Frozen-EZ Yeast Transformation II Kit (Zymo Research) and plated on agar containing Sc-Leu minimal media (6.7 g/L yeast nitrogen base without amino acids, 1.62 g/L Drop-out Mix Synthetic without Leucine without YNB (US Biological), 20 g/L D-glucose) for 2 days at 30 °C.

To prepare CYP72A1709-enriched *S. cerevisiae* microsomes, 1 mL of Sc-Leu minimal media was inoculated with a positive transformant carrying the pESC-leu2d vector with *CYP72A1709* and *CPR-1*, and grown at 30 ºC, 200 rpm overnight. Then, 25 mL of YPD media (10 g/L yeast extract, 20 g/L peptone, 20 g/L D-glucose) was inoculated with overnight culture and incubated at the same conditions for 35 hrs. After centrifugation (3200 × *g*, 4 °C, 5 min), cells were resuspended and induced in 25 mL of YP-Gal media (same as YPD, but 20 g/L galactose instead of D-glucose) and incubated at 25 °C and 220 rpm for 18 hrs. After galactose induction, cells were centrifuged (7500 × *g*, 4 °C, 10 min) and resuspended in 7.5 mL TEK buffer (50 mM Tris-HCl pH 7.5, 1 mM EDTA, 100 mM KCl). Cells were centrifuged again, resuspended in 2 mL TES B buffer (50 mM Tris-HCl pH 7.5, 1 mM EDTA, 600 mM sorbitol, 10 g/L bovine serum albumin fraction V (freshly added) and 1 mM DTT (freshly added)), and lysed by vigorous shaking with ~2 mL glass disruption beads (0.5 mm diameter, RPI) for 10 min (30 sec of shaking, 30 sec on ice, 10 cycles). The crude lysate was recovered by washing the beads with 5 mL TES B buffer four times. The combined lysate was then centrifuged (7500 × *g*, 4 °C,

10 min), after which only the supernatant was transferred to an ultracentrifuge tube and ultracentrifuged (115,000 × *g*, 4 °C, 90 min). Post-ultracentrifugation, the supernatant was removed, and the microsome pellet was gently washed with 2.5 mL TES B buffer, then 2.5 mL TEG buffer (50 mM Tris-HCl pH 7.5, 1 mM EDTA, 20% glycerol), and finally homogenized with a polypropylene pestle for microcentrifuge tubes (USA Scientific) in 500 µL TEG buffer. Aliquots were flash frozen in liquid nitrogen and stored at -80 °C for future use.

To perform *in vitro* microsomal assays, reactions were typically carried out in 50 µL total reaction volumes containing 20 µg of CYP72A1709-enriched microsomes, 10 µM substrate, and 1 mM NADPH in 20 mM potassium phosphate buffer (pH 7.5). Assays were incubated at 30 ºC overnight, quenched by adding 1:1 (v/v) methanol, vortexed for 5 min, centrifuged (16000 × g, 5 min), supernatant filtered through 0.45 µm PTFE filters before LC-MS analysis. Control assays were established by either omitting NADPH or assaying microsomes harboring an empty pESC-leu2d vector.

*In vitro* assays using recombinant ATH, 4CL, *Mc*AT-1, *Pa*BauA

Codon-optimized synthetic genes for ATH and 4CL, *Mc*AT-1, and *Pa*BauA were synthesized by Twist Bioscience for recombinant expression in *E. coli* strain BL21. Coding sequences were cloned into the pET-28a(+) vector using NdeI and XhoI restriction digest sites (N-terminal 6xHis-tag). ATH mutants were generated according to the protocol described in the Q5 Site-Directed Mutagenesis Kit (New England Biolabs) using plasmid pET-28a::*Mt*ATH as the template and the primer sequences incorporating the desired mutations. Sequence-verified constructs were transformed into *E. coli* BL21 (DE3) for recombinant protein production. Starter cultures grown overnight were used to inoculate 1 L of TB media containing kanamycin (50 µg/mL) and incubated at 37 ºC; 200 rpm until reaching an $OD_{600}$ of 0.6–0.8. Cultures were cold-shocked, induced with 0.5 mM IPTG (isopropyl β-*D*-1-thiogalactopyranoside), and incubated at 16 ºC for 18 hours; 200 rpm. Cells were harvested via centrifugation (~10 g cell pellet per L) and resuspended in lysis buffer (100 mM Tris-HCl pH 8.0, 200 mM NaCl, 20 mM imidazole, 10% v/v glycerol, 1 mM DTT) supplemented with lysozyme (1 mg/mL), 1 mM PMSF, trace DNaseI. Then, cell suspensions were lysed via sonication (45% amplitude, 30 sec on and 30 sec off cycles for 5 min) using a 1/2-inch disruptor horn (Branson Sonifier SFX550). Lysates were clarified by centrifugation (48,400 × *g*, 1 hour, 4 ºC) and loaded onto Ni-NTA agarose resin (Cube Biotech) for gravity-flow affinity purification. Ni-NTA resin was washed with 20 column volumes of lysis buffer, and target proteins were eluted using 3 column volume of elution buffer (100 mM Tris-HCl pH 8.0, 200 mM NaCl, 250 mM imidazole, 10% v/v glycerol, 1 mM DTT). Eluants were desalted using PD-10 desalting columns packed with Sephadex™ G-25 (Cytiva) for rapid buffer exchange into storage buffer (50 mM Tris-HCl pH 8.0, 5% v/v glycerol). Purified proteins were concentrated using Vivaspin Polyethersulfone Centrifugal Concentrators (Sartorius). N-terminal 6xHis-tag cleavage was not necessary to observe *in vitro* activity. SDS-polyacrylamide gel electrophoresis was conducted throughout the purification process to evaluate quality and purity of the recombinant proteins.

Assays for *Mc*AT-1 or ATH and its mutants with each substrate combinations (*p*-coumaric acid (**3**) and *p*-tyramine (**9**); ferulic acid (**6**) and *p*-tyramine (**9**); *p*-coumaric acid (**3**)

and *N*-methyltyramine (**10**); ferulic acid (**6**) and *N*-methyltyramine (**10**); including other substrate pairs in fig. S18) were carried out in 20 mM HEPES with 1 mM substrates, 1 mM ATP, 1 mM CoA, 0.5 mM $CaCl_2$, and 5 mM $MgCl_2$. 10 μM of 4CL was added to each assay 3 mins before the addition of the respective enzyme at 10 μM. For combined assays with CYP72A1709-enriched microsomes, 20 μg of microsomes and 1 mM NADPH were added. For malonate semialdehyde (**24**) byproduct trapping experiments, 5 μM *Pa*BauA, 1 mM PLP, and 1 mM (S)-(−)-α-Methylbenzylamine (**25**) were added. For enzyme assays in $H_2{}^{18}O$, reaction was carried out in 70% $H_2{}^{18}O$ by preparing necessary stock solutions in 97% $H_2{}^{18}O$. All assays were incubated at 30 ºC overnight, quenched by adding 1:1 (v/v) methanol, vortexed for 5 min, centrifuged (16,000 × g, 5 min), supernatant filtered through 0.45 μm PTFE filters before LC-MS analysis. For MS/MS analysis of these assays, fragmentation of **12** ($[M+H]^+$ = *m/z* 302.1387) produced diagnostic fragments at *m/z* 165.0546 and *m/z* 121.0648 (fig. S13I-K), which were used to localize the $^{18}O$ label. Dansylated β-alanine (**26**) and its isotopologues were identified by MS/MS against a dansylated β-alanine standard ($[M+H]^+$ = *m/z* 323.1060; fragments *m/z* 157.0884, 170.0962, 234.0580, 252.0683, 308.0822) (fig. S16D–G).

<u>Chemical derivatization for LC-MS detection</u>

To derivatize β-alanine (**26**) for LC-MS detection (Fig. 5E, fig. S16), dansyl chloride (DNS) solution was first generated by making 50 mM DNS in 100% acetonitrile, then mixing 1:1 (v/v) of 50 mM DNS with carbonate buffer (100 mM $Na_2CO_3$/$NaHCO_3$, pH 9.8). This DNS solution was then added 2:1 (v/v; DNS solution:reaction mixture) instead of methanol to the reaction, vortexed, incubated for 1 hour at room temperature stored away from light. DNS derivatization was quenched by adding 1:10 (v/v; 10% ammonium hydroxide:reaction mixture) 10% ammonium hydroxide, incubated at room temperature for 5 mins, centrifuged (16000 × g, 5 min), supernatant filtered through 0.45 μm PTFE filters before LC-MS analysis.

To capture potential acyl thioester intermediates (fig. S13E, G), hydroxylamine derivatization was utilized. To reactions containing intermediates, hydroxylamine (1 M stock in 20 mM HEPES, pH 8.5) was added 1:1 (v/v) with reaction mixture, incubated at 30 ºC for 30 mins, quenched with 1:1 (v/v) methanol, centrifuged (16000 × g, 5 min), supernatant filtered through 0.45 μm PTFE filters before LC-MS analysis.

<u>Activity-guided fractionation and proteomic MS</u>

Aerial tissue from soil-grown six-month old *M. tortuosum* (~34 g fresh weight) was ground to fine powder under liquid nitrogen using a mortar and pestle, then extracted in ice-cold extraction buffer (100 mM Tris-HCl pH 8, 10 mM DTT, 10% glycerol, 1% w/v PVPP, protease inhibitor cocktail) (fig. S10). The homogenate was clarified by centrifugation (20,000 × *g*, 4 ºC, 45 min), and the supernatant was ultracentrifuged (115,000 × *g*, 4 °C, 90 min). The resulting soluble fraction was concentrated using the 10 kDa MWCO Vivaspin Polyethersulfone Centrifugal Concentrator (Sartorius) prior to chromatography.

Reductase activity in each fraction was assayed by incubating 5 μL of fraction with 1 μM mesembrenone (**19**) and 1 mM NADPH in 50 mM Tris-HCl pH 8 at 30ºC for 1 hour. Reactions were quenched with 1:1 (v/v) methanol. Mesembrine (**20**, $[M+H]^+$ = *m/z* 290.1751) production and residual **19** ($[M+H]^+$ = *m/z* 288.1594) were detected using the C18 - 9 min LC-MS method. Control reactions lacked NADPH.

All chromatography was performed at 4 ºC on the GE ÄKTA Purifier fast protein liquid chromatography (FPLC) system (Cytiva), with $A_{280}$ and conductivity monitored. The following sequential rounds were performed:

*Round 1: Size exclusion.* Size-exclusion chromatography (SEC) on the Superdex 200 10/300 GL column (Cytiva) column equilibrated in size-exclusion buffer (50 mM Tris-HCl, pH 8, 1 mM DTT) (fig. S10C). The column was calibrated with a gel filtration chromatography standard mixture (BioRad). Each run was eluted isocratically over 1.2 column volumes (CV) at 0.5 mL/min and collected in 0.5 mL fractions.

*Round 2: Anion exchange.* Active fractions from round 1 were pooled and further separated by anion-exchange chromatography (AEC) on the HiTrapQ HP (Cytiva) column using buffers A (50 mM Tris-HCl, pH 8) and B (50 mM Tris-HCl, pH 8, 1 M NaCl) with a linear gradient from 0 to 30% buffer B (equivalent to 0 to 300 mM NaCl) over 20 CV. Activity eluted at ~11-15 mS/cm, which is approximately 125-170 mM NaCl (fig. S10D).

*Round 3: Polishing size exclusion.* For the final round of fractionation, the active fractions from round 2 were pooled and run on the same Superdex 200 10/300 GL column (Cytiva) under the Round 1 condition, except at 0.25 mL/min with 0.25 mL fractions (fig. S10E).

Four fractions from round 3 (fig. S10F) were submitted for protein mass spectrometry (pMS). The Proteomics Core Facility of the Harvard Center for Mass Spectrometry performed tryptic digestion and LC-MS/MS. Raw files were processed in Proteome Discoverer 3.2 (Thermo Fisher Scientific). Detected MS/MS spectra were searched with Sequest HT (*90*) and MS Amanda 3.0 (*91*) against the *M. tortuosum* proteome and transcriptome, combined with a database of common contaminants. Peptide-spectrum matches (PSMs) from both search engines were rescored with Percolator (*92*) using a concatenated target-decoy strategy and filtered at a 1% false discovery rate (FDR, q-value) at the PSM and peptide levels. Only high-confidence peptides were kept. Proteins were grouped under strict parsimony, and master proteins with at least one peptide were reported at a 1% protein-level FDR. The number of PSMs per protein served as a semi-quantitative measure of abundance across fractions. All matched proteins were annotated by Pfam domain content and by BLASTP against *A. thaliana* and *M. crystallinum* proteome. Candidates were further narrowed by three criteria: 1) a PSM profile that tracked activity in fig. S10F, 2) annotation as an oxidoreductase, and 3) co-expression with known mesembrenone pathway genes. This ultimately led to the identification of KDR-1 (*g1529*) from pMS.

#### Metabolite purification and structural elucidation via NMR spectroscopy

For preparative metabolite isolation, pathway genes were transiently expressed in *N. benthamiana* for 7 days. Infiltrated tissue was harvested, lyophilized, and processed as described above. For all metabolites purified (**12**, **13**, **14**, **16**, and **17**), tissue was extracted with 100% methanol overnight at room temperature with shaking at 30 rpm. Extraction was repeated twice in half the initial methanol volume for 1 hour each. Extracts were combined, filtered, and dried by rotary evaporation at 30 ºC. Dried residues were resuspended in water + 0.1% formic acid for subsequent preparative HPLC purifications. Resuspended extracts were injected onto a Infinity Lab Poroshell 120 SB C-18 column (120Å, 4 µm, 21.2 mm × 150 mm) (Agilent) and separated using water + 0.1% formic acid (solvent A) and acetonitrile + 0.1% formic acid (solvent B); 5-40% solvent B gradient on a Pure C-850 FlashPrep instrument (Buchi) with fractions collected based on UV absorbance at 272 nm. Fractions were screened by LC-MS and appropriate fractions were pooled for injection onto a XBridge BEH HILIC OBD column (130Å, 5 µm, 19 mm × 150 mm) (Waters) and separated using water + 0.1% formic acid and 10 mM ammonium formate (solvent A) and 90% acetonitrile, 10% water + 0.1% formic acid and 10 mM ammonium formate (solvent B); 100-70% solvent B gradient with fractions collected based on $\lambda_{272}$ and screened via LC-MS. This process was repeated until a final fraction containing pure compound was obtained. Each of the purified metabolites yielded approximately 1 mg except **14** which yielded ~0.2 mg.

For structure determination using NMR, each purified compound was resuspended in either deuterated methanol (MeOD) or deuterated DMSO (DMSO-$d_6$). All spectra were acquired on a Bruker Avance 600 MHz spectrometer. NMR data were processed using MestReNova (Mestrelab Research). Chemical shifts were referenced to residual MeOD or DMSO-$d_6$. Automatic phase correction was applied. For baseline correction, the Whittaker baseline correction was applied. Assignments supporting the structures of each metabolite were established by standard analysis of 1D and 2D spectra.

#### Protein structure prediction and structural comparison

Structural models of ATH (*g4212.t1*) and *Mc*AT-1 (*Mcr-012483*) in complex with **12** were generated with Boltz-2 (v2.2.1) using the protein sequence and ligand SMILES as input, with 5 diffusion samples and the default 5 recycling steps. Both *S*- and *R*-diastereomer ligand complexes were modeled with ATH and *Mc*AT-1. The structural models were superimposed in PyMOL (v3.1.3.1) and the active-site pocket defined as residues within 5 Å of the docked ligand. Residues differing between ATH and McAT-1 within this pocket were identified from the structure-guided alignment in Fig. 4E and targeted for mutagenesis. Figures were rendered in PyMOL.

#### Phylogenetic tree construction, phylogenomic analysis and divergence time estimation

For each enzyme family of interest (ATH, CYP98A286, KDR-1), homologs were retrieved by BLASTp (v2.15.0+) of the *M. tortuosum* query protein against the NCBI non-redundant protein database (*e*-value cutoff $1 \times 10^{-20}$, top 500 hits), supplemented with the top 5 BLASTp hits from each genome listed in table S1, and if relevant, with previously characterized enzymes of each family. Redundant sequences were removed using CD-HIT at 99% identity and sequences outside the expected length range were excluded. Sequences were aligned with MAFFT (v7.526) (*93*) using the L-INS-i algorithm (--localpair --maxiterate 1000 --leavegappyregion) and trimmed with ClipKIT (v2.12.0) (*94*) (default smart-gap mode). Maximum-likelihood trees were inferred with IQ-TREE (v3.1.1) with 1,000 bootstrap replicates. Trees were visualized with iTOL. The BAHD tree in Fig. 4D comprises ATH and 182 orthologous proteins and includes characterized members of BAHD clades I–V and tyramine hydroxycinnamoyltransferase (THT) from *Oryza sativa*. The CYP98 tree (fig. S7A) includes characterized C3H enzymes and non-CYP98 outgroups; the KDR/OPR tree (fig. S12B) includes characterized 12-oxophytodienoate reductases. All characterized sequences were obtained from the UniProt database.

Genome assemblies and annotations for 13 species in table S1 were used for comparative genomic analysis. For each species, redundant sequences were removed using CD-HIT at 90% identity prior to orthology inference. Orthogroups were inferred with OrthoFinder (v2.5.5; -S diamond -M msa -A muscle -T raxml-ng -I 1.5) from the primary protein sequence of each gene, yielding 919 single-copy orthologs. Each single-copy ortholog was aligned at the protein level with MAFFT (v7.526, L-INS-i: --localpair --maxiterate 1000 --leavegappyregion), back-translated to a codon-level alignment with PAL2NAL (*95*), and trimmed with trimAl (v1.5.1) (*96*). Then, a maximum-likelihood gene tree was inferred for each alignment with IQ-TREE (v3.1.2; -m MFP, 1,000 bootstraps) (*97*), rooted on the two monocots in the dataset (*Hordeum vulgare*, *Lycoris radiata*) as outgroup. Based on these gene trees, a coalescent-based species tree was inferred using ASTRAL-IV/ASTER (v1.25.4.8). Divergence times were estimated with MCMCtree (PAML v4.10.10) using three soft-bound calibrations: 1) monocot–eudicot split at 130–180 millon years ago (MYA) (*98*, *99*), 2) Caryophyllales node at 100–130 MYA (*100*, *101*), and 3) Caryophyllaceae stem at 72–86 MYA (*99*).

### Supplementary Figures

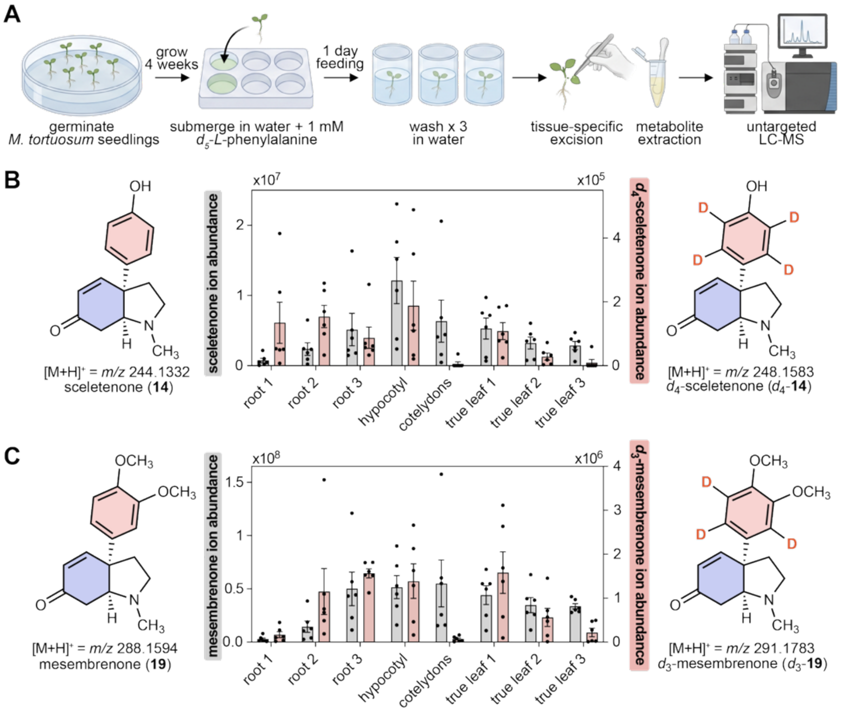


**fig. S1. Stable isotope tracing using $d_5$-$L$-phenylalanine in *M. tortuosum* seedlings.**
(**A**) Schematic of the $d_5$-$L$-phenylalanine (**1**) tracing experiment in *M. tortuosum*, generated using BioRender. (**B**) LC-MS profiling of tentative sceletenone (**14**) ($[M+H]^+$ = *m/z* 244.1332; gray, left) and $d_4$-sceletenone ($d_4$-**14**) ($[M+H]^+$ = *m/z* 248.1583; pink, right). (**C**) LC-MS profiling of mesembrenone (**19**) ($[M+H]^+$ = *m/z* 288.1594; gray, left) and $d_3$-mesembrenone ($d_3$-**19**) ($[M+H]^+$ = *m/z* 291.1783; pink, right). For panels B and C, the bar graphs represent the mean of six biological replicates across eight tissues as outlined in Figure 1C with error bars representing SEM.

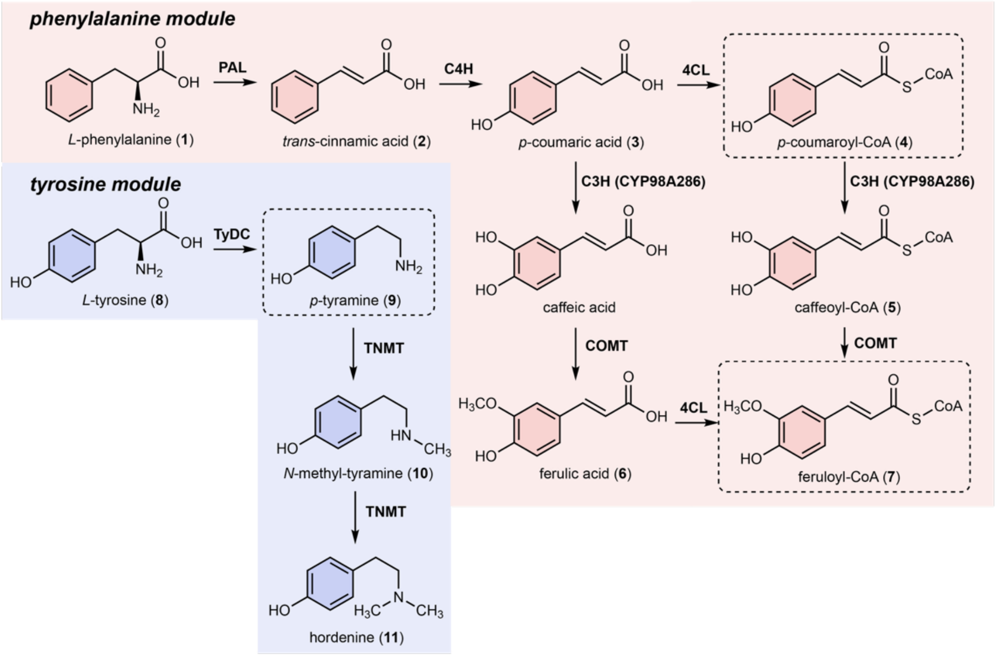


**fig. S2. Proposed pre-scaffold pathway modules discussed in this study.**
The phenylalanine module (pink) ultimately converts *L*-phenylalanine (**1**) to *p*-coumaroyl-CoA (**4**) and feruloyl-CoA (**7**), which are used for Kanna alkaloid scaffold formation. The tyrosine module (blue) converts *L*-tyrosine (**8**) to *p*-tyramine (**9**), which enter the scaffolding modules in **fig. S3**.

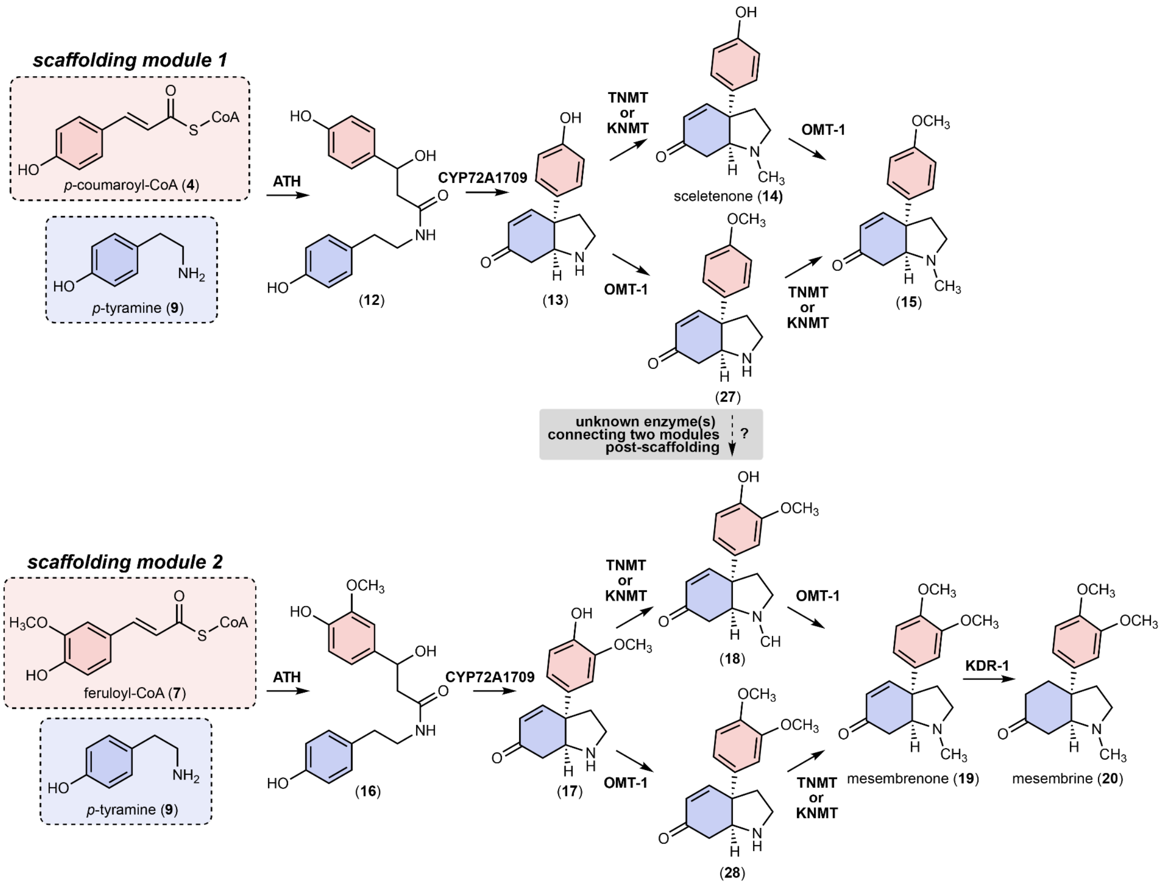


**fig. S3. Two major proposed scaffolding pathway modules in Kanna alkaloid biosynthesis.** Scaffolding module 1 (top) uses ATH to condense and hydrate *p*-coumaroyl-CoA (**4**) and *p*-tyramine (**9**) to generate a linear amide **12**, which CYP72A1709 converts to **13**. Scaffolding module 2 (bottom) uses ATH to condense and hydrate feruloyl-CoA (**7**) and *p*-tyramine (**9**) to generate a linear amide **16**, which CYP72A1709 converts to **17**. The gray box indicates unidentified enzyme(s) that could connect the two modules post-scaffolding.

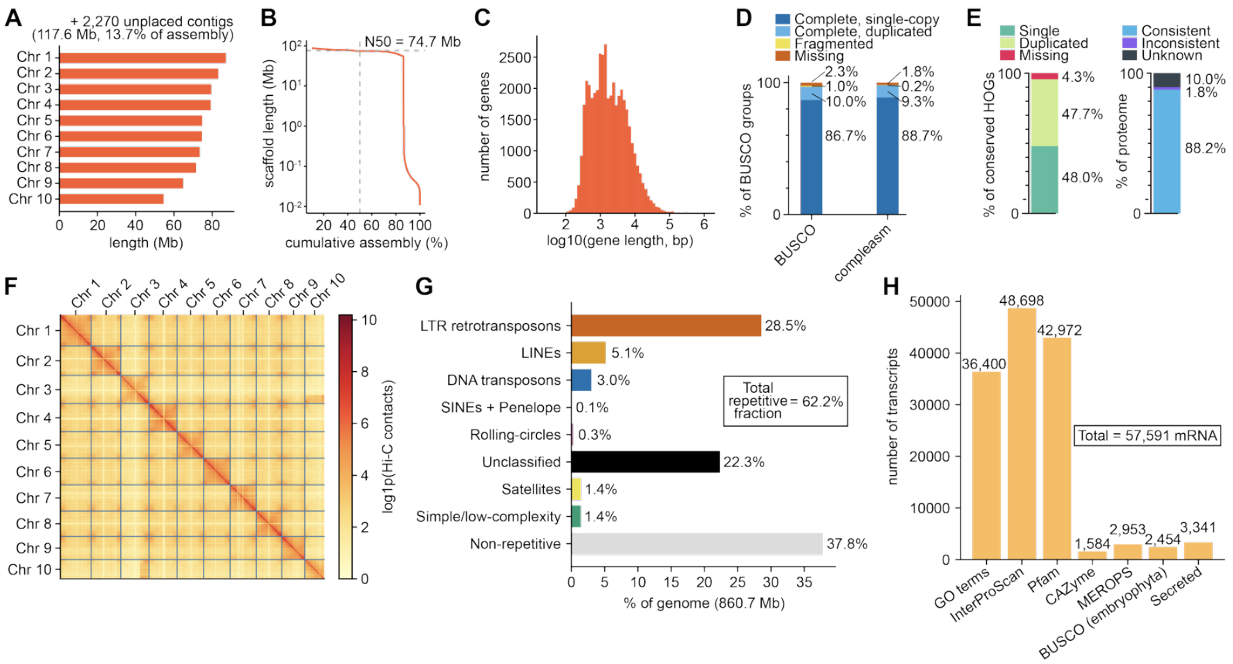


**fig. S4. Assembly and annotation of the *M. tortuosum* genome.**
(**A**) Length of each chromosome in *M. tortuosum* genome assembly from this study (743.2 Mb, 86.3% of assembly). (**B**) Nx contiguity plot with N50 = 74.7 Mb. (**C**) Length distribution of 42,934 annotated genes from BRAKER3 with median of 1,569 bp and mean of 4,349 bp. (**D**) *M. tortuosum* genome assembly completeness assessed by BUSCO and compleasm using eudicots_odb10 database. (**E**) Proteome completeness (left) and consistency (right) as assessed using OMArk compared to Viridiplantae OMAmer database. (**F**) Hi-C contact map with chromosome-scale scaffolds (500 Kb bins). (**G**) Repeat landscape annotated by RepeatMasker and EarlGrey with total repetitive fraction of 62.2%. (**H**) Summary of functional annotation coverage of 57,591 mRNAs.

**A**

| gene | PAL r | PAL rank | TyDC r | TyDC rank | characterized function | Pfam annotation | BLASTp annotation | identity % M. crystallinum | identity % A. thaliana |
|---|---|---|---|---|---|---|---|---|---|
| *g30732* | 1.000 | 0 | 0.219 | 42 | PAL | Aromatic amino acid lyase | Phenylalanine ammonia-lyase | 99.3 | 85.5 |
| *g29247* | 0.354 | 1 | 0.312 | 5 | C4H | Cytochrome P450 | Trans-cinnamate 4-monooxygenase | 97.6 | 84.5 |
| *g37187* | 0.338 | 2 | 0.323 | 2 | 4CL | AMP-binding enzyme | 4-coumarate--CoA ligase 1 | 97.3 | 68.2 |
| *g19370* | 0.302 | 4 | 0.339 | 1 | CYP72A1709 | Cytochrome P450 | Cytochrome P450 CYP72A219 | 32.2 | 50.4 |
| ⋮ | | | | | | | | | |
| *g4212* | 0.282 | 7 | 0.318 | 3 | ATH-1 | Transferase family | Pelargonidin 3-O-(6-caffeoylglucoside) 5-O- (6-O-malonylglucoside) 4'''-malonyltransferase | 75.0 | 33.0 |
| *g22226* | 0.280 | 8 | 0.244 | 22 | C3H (CYP98A286) | Cytochrome P450 | Cytochrome P450 98A2 | 97.7 | 68.5 |
| ⋮ | | | | | | | | | |
| *g272* | 0.240 | 17 | 0.268 | 12 | OMT-1 | O-methyltransferase domain | Cathecol O-methyltransferase 1 | 78.0 | 54.2 |
| ⋮ | | | | | | | | | |
| *g35047* | 0.219 | 28 | 1.000 | 0 | TyDC | Pyridoxal-dependent decarboxylase conserved domain | Tyrosine decarboxylase | 98.5 | 53.4 |
| ⋮ | | | | | | | | | |
| *g20977* | 0.173 | 85 | 0.219 | 41 | TNMT | Mycolic acid cyclopropane synthetase | (S)-coclaurine N-methyltransferase | 76.7 | 54.7 |
| ⋮ | | | | | | | | | |
| *g20975* | 0.084 | 623 | 0.120 | 289 | KNMT | Mycolic acid cyclopropane synthetase | (S)-coclaurine N-methyltransferase | 74.4 | 55.2 |
| ⋮ | | | | | | | | | |
| *g1529* | -0.063 | 25,770 | -0.013 | 23,484 | KDR-1 | NADH:flavin oxidoreductase / NADH oxidase family | Putative 12-oxophytodienoate reductase 11 | 87.3 | 74.7 |

**B**

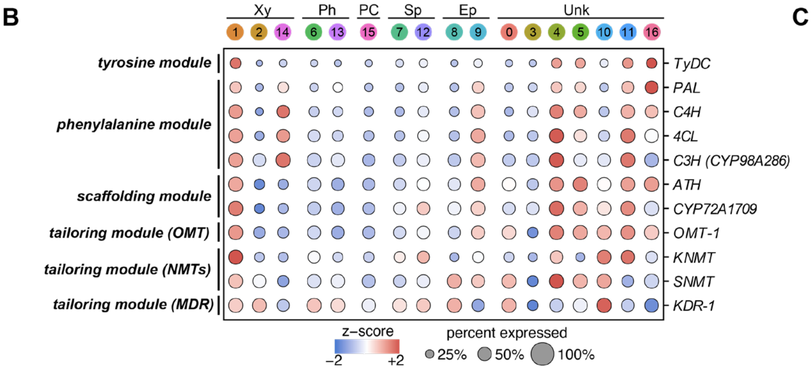


**C**

| cell type | clusters | # cells |
|---|---|---|
| xylem | 1, 2, 14 | 4,537 |
| phloem | 6, 13 | 1,582 |
| phloem-companion | 15 | 300 |
| S-phase | 7, 12 | 1,241 |
| epidermis | 8, 9 | 835 |
| unknown | 0, 3, 4, 5, 10, 11, 16 | 7,134 |

(annotated based on *M. crystallinum*)

**fig. S5. *M. tortuosum* snRNA-seq co-expression analysis and cell-type annotation.**
(**A**) Table of mesembrine (**20**) biosynthetic genes ranked according to Spearman's correlation to *PAL* expression across 15,629 nuclei. Also shown in this table are the corresponding ranks to *TyDC* expression, the characterized function of each gene, Pfam and BLASTp annotations, and amino-acid identity (%) to the closest orthologs in *M. crystallinum* and *A. thaliana*. (**B**) Dot plot showing cluster-specific expression of mesembrine biosynthetic genes, grouped by pathway module. Dot color indicates z-scored mean expression, and dot size represents the percent of nuclei expressing each gene. (Xy, xylem; Ph, phloem; PC, phloem-companion; Sp, S-phase; Ep, epidermis; Unk, unknown) (**C**) Nuclei per cluster and annotated cell type in the *M. tortuosum* snRNA-seq dataset. Clusters are annotated based on *M. crystallinum* snRNA-seq atlas.

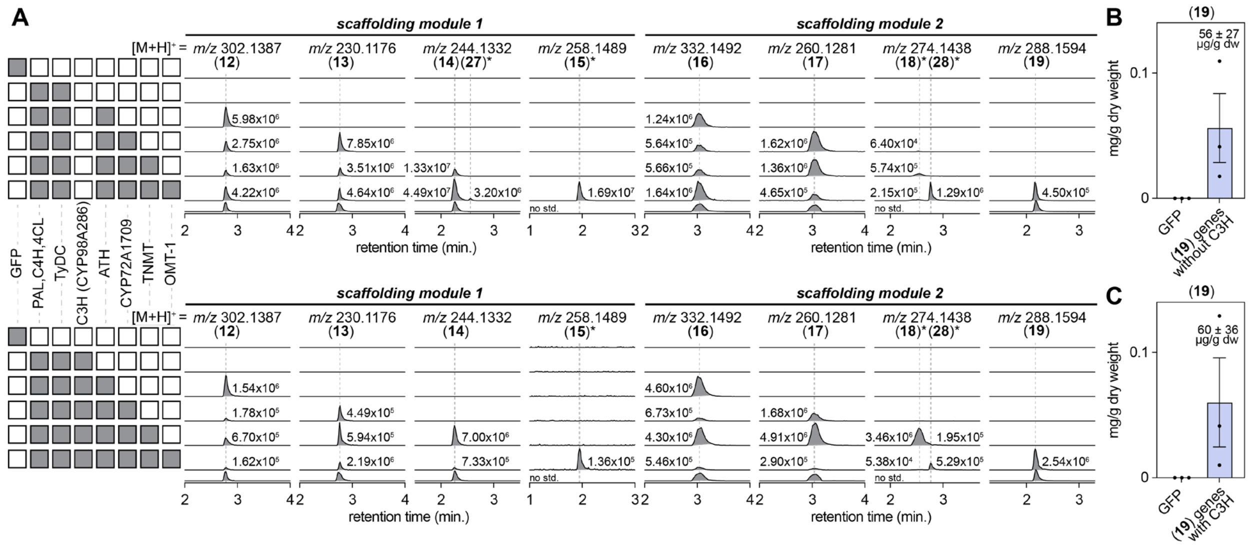


**fig. S6. Transient expression of mesembrenone biosynthetic genes in *N. benthamiana*.**
(**A**) LC-MS chromatograms of mesembrenone (**19**) pathway build-up samples, shown without (top) and with (bottom) C3H (CYP98A286). Filled squares on the left indicate genes co-infiltrated in each sample and peak areas are annotated above each EIC peak. Columns show scaffolding module 1 (**12** = *m/z* 302.1387; **13** = *m/z* 230.1176; **14** and **27*** = *m/z* 244.1332; **15** = *m/z* 258.1489; all $[M+H]^+$) and scaffolding module 2 (**16** = *m/z* 332.1492; **17** = *m/z* 260.1281; **18** and **28** = *m/z* 274.1438; **19** = *m/z* 288.1594; all $[M+H]^+$). An asterisk indicates predicted structures with no purified or authentic standard available; these structures are inferred from LC-MS exact mass and biosynthetic pathway logic. (**B**) Quantification of mesembrenone (**19**) in samples containing the (**19**) gene set without C3H (56 ± 27 µg/g dry weight). The calibration curve for mesembrenone was generated from its authentic standard at 10 concentrations spanning 0.25 nM to 1 µM and fitted by linear regression ($R^2$ = 0.9989). Mesembrenone was quantified using the following equation [mesembrenone in nM] = (ion count - 107892)/27940. Bars show mean ± SEM with three biological replicates. GFP is shown as negative control. (**C**) Quantification of mesembrenone (**19**) in samples containing the (**19**) gene set with C3H (60 ± 36 µg/g dry weight). Mesembrenone was quantified using the same equation as above. Bars show mean ± SEM with three biological replicates. GFP is shown as negative control.

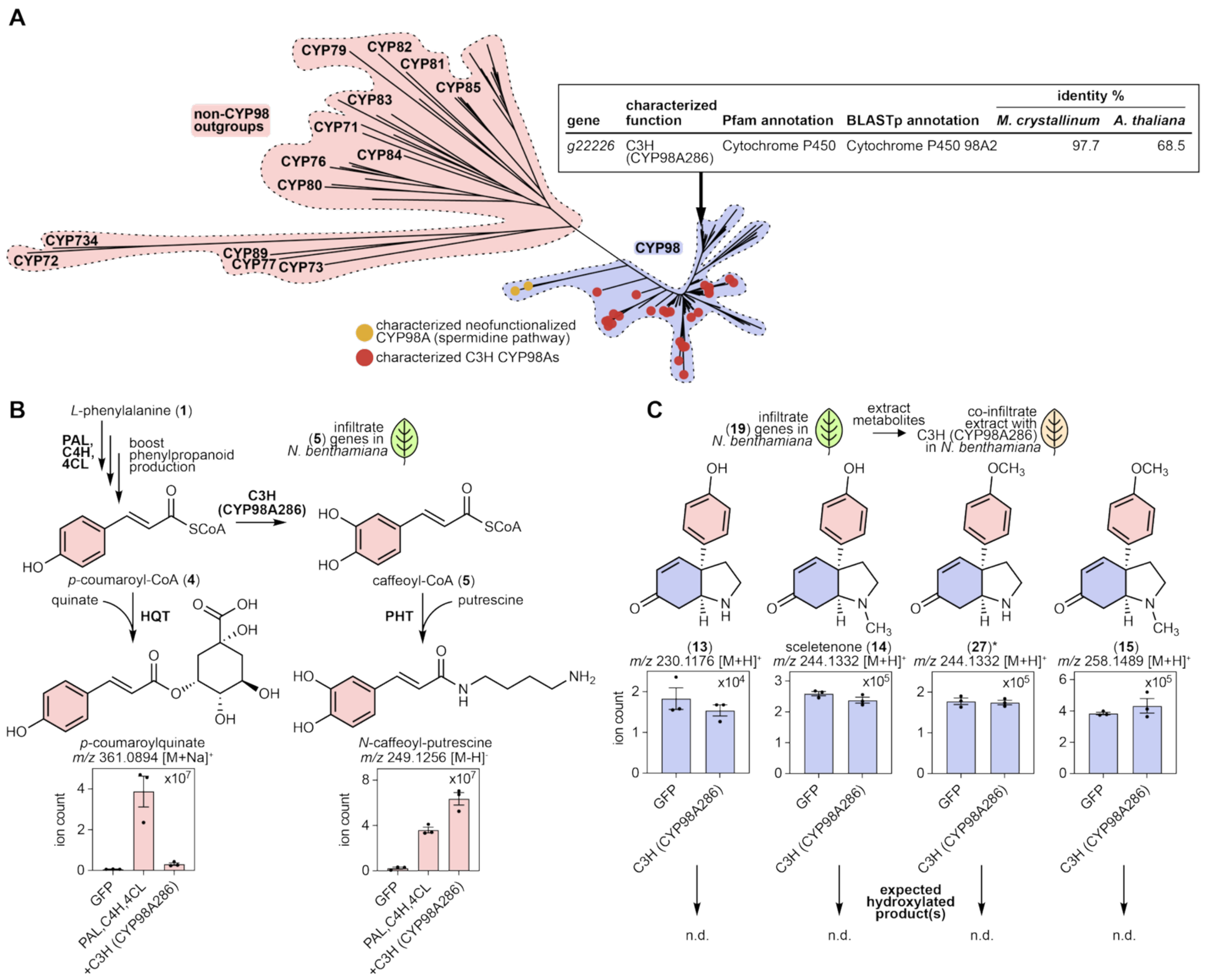


**fig. S7. Verification of CYP98A286 as a C3H for phenylpropanoid metabolism.**
(**A**) Maximum-likelihood phylogenetic tree of CYP98A286, which belongs to the characterized C3H clade that hydroxylates phenylpropanoid substrates. Red circles mark characterized C3H CYP98As, yellow circles mark the characterized neofunctionalized CYP98A in the spermidine pathway, and non-CYP98 CYP families are shown as outgroups. Table portraying functional annotation of CYP98A286 and the amino acid identity to its orthologs found in *M. crystallinum* (97.7%) and *A. thaliana* (68.5%). Arrow points to C3H (CYP98A286) from *M. tortuosum*. (**B**) Expression of CYP98A286 in *N. benthamiana* along with phenylpropanoid metabolic enzymes PAL, C4H, and 4CL. CYP98A286 activity leads to a decrease in *p*-coumaroyl-quinate ($[M+Na]^+$ = *m/z* 361.0894) and an increase in *N*-caffeoyl-putrescine ($[M-H]^-$ = *m/z* 249.1256). Each were validated with their respective chemical standards. (**C**) Expression of CYP98A286 in *N. benthamiana* co-infiltrated with crude metabolites extracted from *N. benthamiana* leaf expressing all (**19**)-producing genes without CYP98A286 (*PAL*, *C4H*, *4CL*, *TyDC*, *ATH*, *CYP72A1709*, *TNMT* and *OMT-1*). CYP98A286 does not consume intermediates from the scaffolding module 1 pathway (**13** = *m/z* 230.1176; **14** = *m/z* 244.1332; **27** = *m/z* 244.1332; **15** = *m/z* 258.1489, all $[M+H]^+$) and no production of the expected hydroxylated product(s) is observed (n.d. = not detected). Bars show mean ± SEM with three biological replicates.

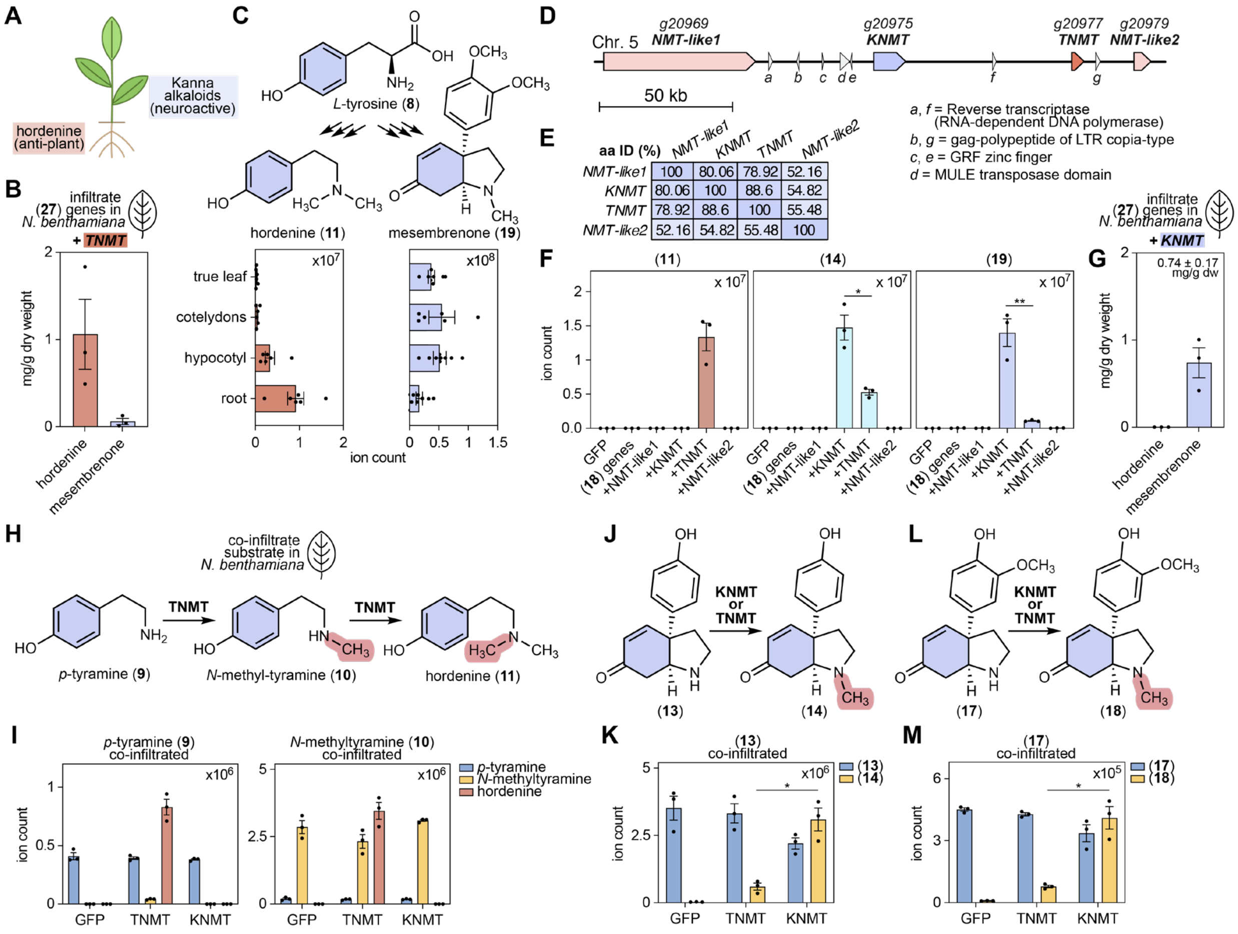


**fig. S8. TNMT and KNMT have distinct biosynthetic functions.**

(**A**) Schematic representation of *M. tortuosum* and the preferred site of accumulation for hordenine (**11**) and Kanna alkaloids. (**B**) Quantification of hordenine (**11**) (1.06 ± 0.40 mg/g of dry leaf weight) and mesembrenone (**19**) (60 ± 36 µg/g of dry leaf weight) produced in *N. benthamiana* when *TNMT* is expressed with (**27**) pathway genes (*PAL*, *C4H*, *4CL*, *TyDC*, *ATH*, *CYP72A1709*, and *OMT-1*), reported as mg/g dry weight. (**C**) Reaction schematic of the competing *N*-methylation routes from *L*-tyrosine (**8**) to hordenine (**11**) and to mesembrenone (**19**), and LC-MS ion counts of **11** ($[M+H]^+$ = *m/z* 166.1226; left) and **19** ($[M+H]^+$ = *m/z* 288.1594; right) across the following *M. tortuosum* tissues: true leaf, cotyledon, hypocotyl and roots. (**D**) Organization of the *N*-methyltransferase locus on chromosome 5 of the *M. tortuosum* genome, containing *NMT-like1* (*g20969*), *KNMT* (*g20975*), *TNMT* (*g20977*) and *NMT-like2* (*g20979*), interspersed with transposon-derived genes. (**E**) Pairwise amino-acid identity (%) among the four paralogous NMTs. (**F**) Ion counts of hordenine (**11**; $[M+H]^+$ = *m/z* 166.1226), sceletenone (**14**; $[M+H]^+$ = *m/z* 244.1332) and mesembrenone (**19**; $[M+H]^+$ = *m/z* 288.1594) in *N. benthamiana* expressing the (**27**) gene set together with *NMT-like1*, *KNMT*, *TNMT* or *NMT-like2*. (**G**) Quantification of hordenine (**11**) and mesembrenone (**19**) produced in *N. benthamiana* when *KNMT* is expressed with (**27**) pathway genes. 0.74 ± 0.30 mg of mesembrenone (**19**) per g of dry leaf weight is produced. (**H**) Reaction schematic of the sequential *N*-methylation of *p*-tyramine (**9**) to *N*-methyl-tyramine (**10**) and hordenine (**11**) by TNMT. (**I**) Ion counts of **9** ($[M+H]^+$ = *m/z* 138.0913), **10** ($[M+H]^+$ = *m/z* 152.1070) and **11** ($[M+H]^+$ = *m/z* 166.1226) in *N. benthamiana* expressing GFP, *TNMT* or *KNMT* and co-infiltrated with **9** (left) or **10** (right). (**J**) Reaction schematic of the *N*-

methylation of **13** to sceletenone (**14**) by KNMT or TNMT. (**K**) Ion counts of **13** ($[M+H]^+$ = *m/z* 230.1176) and **14** ($[M+H]^+$ = *m/z* 244.1332) in *N. benthamiana* expressing GFP, *TNMT* or *KNMT* and co-infiltrated with **13**. (**L**) Reaction schematic of the *N*-methylation of **17** to **18** by KNMT or TNMT. (**M**) Ion counts of **17** ($[M+H]^+$ = *m/z* 260.1281) and **18** ($[M+H]^+$ = *m/z* 274.1438) in *N. benthamiana* expressing GFP, *TNMT* or *KNMT* and co-infiltrated with **17**. All bars show mean ± SEM with three biological replicates and GFP is shown as negative control. Asterisks indicate paired t-test on lognormal-transformed peak areas (* = $p < 0.05$; ** = $p < 0.01$).

| gene | *KNMT* r | *KNMT* rank | *TNMT* r | *TNMT* rank | characterized function | Pfam annotation | BLASTp annotation | identity % *M. crystallinum* | identity % *A. thaliana* |
|---|---|---|---|---|---|---|---|---|---|
| *g20975* | 1.000 | 0 | 0.241 | 124 | KNMT | Mycolic acid cyclopropane synthetase | (S)-coclaurine N-methyltransferase | 74.4 | 55.2 |
| *g272* | 0.403 | 1 | 0.398 | 17 | OMT-1 | O-methyltransferase domain | Cathecol O-methyltransferase 1 | 78.0 | 54.2 |
| *g19370* | 0.350 | 2 | 0.494 | 4 | CYP72A1709 | Cytochrome P450 | Cytochrome P450 CYP72A219 | 32.2 | 50.4 |
| *g4212* | 0.342 | 3 | 0.351 | 32 | ATH | Transferase family | Pelargonidin 3-O-(6-caffeoylglucoside) 5-O- (6-O-malonylglucoside) 4'''-malonyltransferase | 75.0 | 33.0 |
| *g20969* | 0.294 | 6 | -0.182 | 25,922 | NMT-like1 | Mycolic acid cyclopropane synthetase | (S)-coclaurine N-methyltransferase | 63.0 | 51.6 |
| *g1529* | 0.281 | 9 | 0.162 | 302 | KDR-1 | NADH:flavin oxidoreductase / NADH oxidase family | Putative 12-oxophytodienoate reductase 11 | 87.3 | 74.7 |
| *g20977* | 0.241 | 23 | 1.000 | 0 | TNMT | Mycolic acid cyclopropane synthetase | (S)-coclaurine N-methyltransferase | 76.7 | 54.7 |
| *g37187* | 0.195 | 59 | 0.400 | 16 | 4CL | AMP-binding enzyme | 4-coumarate--CoA ligase 1 | 97.3 | 68.2 |
| *g29247* | 0.160 | 133 | 0.322 | 45 | C4H | Cytochrome P450 | Trans-cinnamate 4-monooxygenase | 97.6 | 84.5 |
| *g22226* | 0.135 | 240 | 0.339 | 36 | C3H (CYP98A286) | Cytochrome P450 | Cytochrome P450 98A2 | 97.7 | 68.5 |
| *g35047* | 0.120 | 325 | 0.219 | 152 | TyDC | Pyridoxal-dependent decarboxylase conserved domain | Tyrosine decarboxylase | 98.5 | 53.4 |
| *g20979* | 0.094 | 577 | 0.288 | 73 | NMT-like2 | Mycolic acid cyclopropane synthetase | (S)-coclaurine N-methyltransferase | 65.2 | 51.9 |
| *g30732* | 0.084 | 715 | 0.173 | 264 | PAL | Aromatic amino acid lyase | Phenylalanine ammonia-lyase | 99.3 | 85.5 |

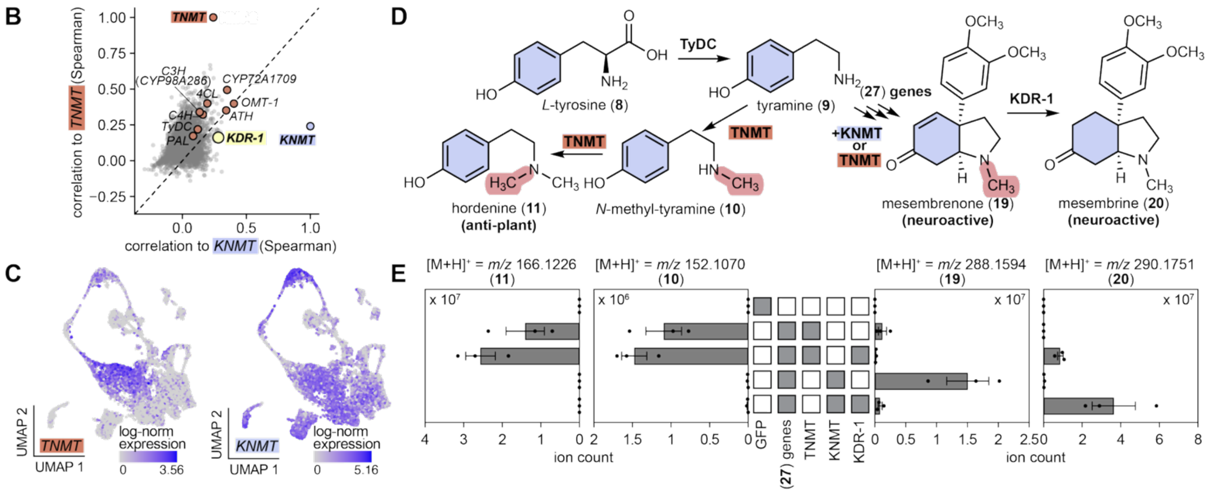


**fig. S9. snRNA-seq co-expression analysis using *KNMT* and *TNMT* as bait genes.**
(**A**) Table of mesembrine biosynthetic genes ranked according to Spearman's correlation to *KNMT* expression across 15,629 nuclei. Also shown in this table is the corresponding rank to *TNMT* expression, characterized gene function, Pfam and BLASTp annotation, and amino acid identity (%) to the closest orthologs in *M. crystallinum* and *A. thaliana*. (**B**) Co-expression plot showing the correlation of *M. tortuosum* genes to *TNMT* and *KNMT*. Spearman correlation coefficients are used. Orange dots indicate mesembrenone biosynthetic genes and yellow dot indicate *KDR-1* gene involved in the conversion of mesembrenone (**19**) into mesembrine (**20**). (**C**) Expression of *TNMT* (left) and *KNMT* (right) (ln(expression + 1)) in *M. tortuosum* nuclei. (**D**) Proposed pathway bifurcation from *L*-tyrosine (**8**) showing TNMT-dependent hordenine (**11**) production versus KNMT-dependent conversion of the **27** gene-set products to mesembrenone (**19**). Mesembrenone (**19**) is then converted to mesembrine (**20**) by KDR-1. (**E**) Ion counts of **11** ($[M+H]^+$ = *m/z* 166.1226), **10** ($[M+H]^+$ = *m/z* 152.1070), **19** ($[M+H]^+$ = *m/z* 288.1594) and **20** ($[M+H]^+$ = *m/z* 290.1751) in *N. benthamiana* expressing the (**27**) gene set with *TNMT* or *KNMT*, and with *KDR-1*. **27** gene set includes *PAL*, *C4H*, *4CL*, *TyDC*, *ATH*, *CYP72A1709*, and *OMT-1*. Bars show mean ± SEM with three biological replicates and GFP is shown as negative control.

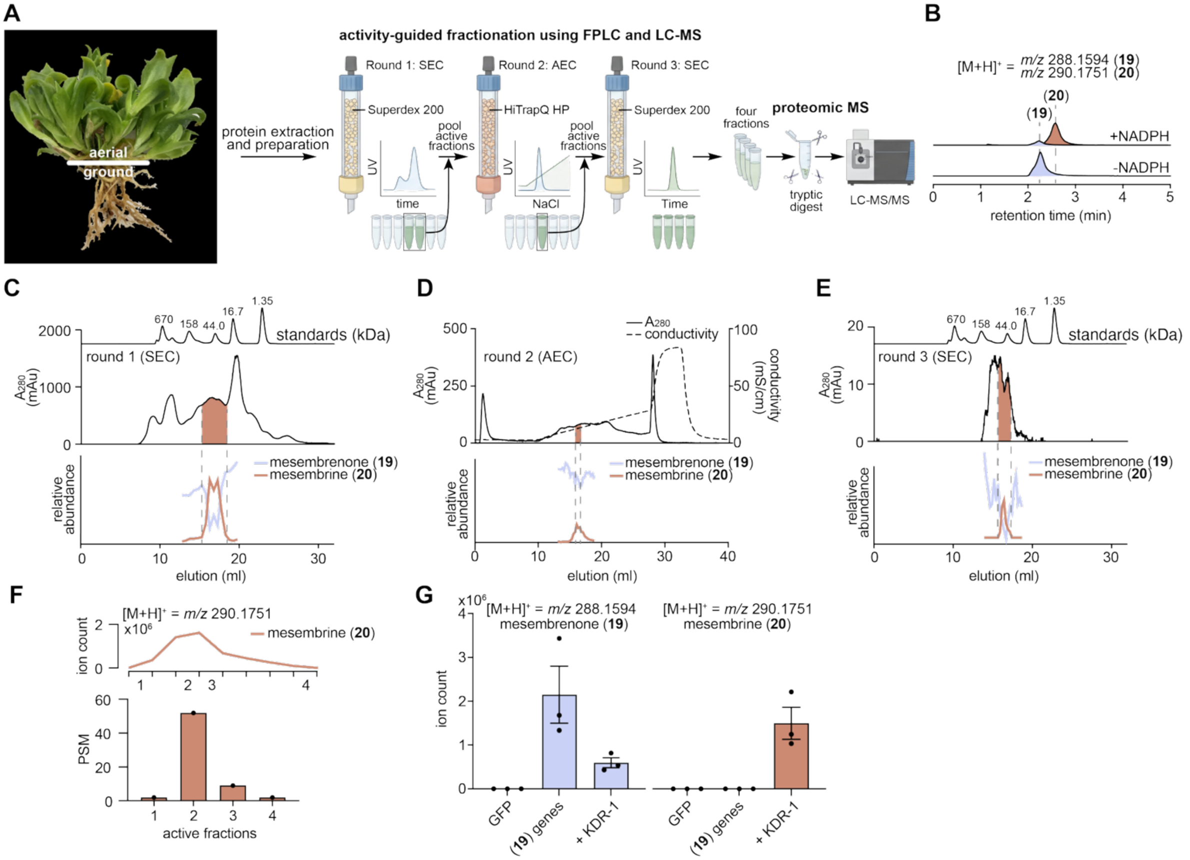


**fig. S10. Activity-guided fractionation of *M. tortuosum* protein lysate identifies KDR-1.**
(**A**) Image of *M. tortuosum* plant collected for crude protein extraction and a schematic of the subsequent activity-guided fractionation, generated using BioRender. (**B**) *In vitro* NADPH-dependent conversion of mesembrenone (**19**) to mesembrine (**20**) using total protein lysate isolated from aerial *M. tortuosum* tissues. (EICs show **19**, $[M+H]^+$ *m/z* 288.1594 and **20**, $[M+H]^+$ = *m/z* 290.1751). (**C**) Round 1, size-exclusion chromatography (SEC) of the crude extract. $A_{280}$ trace (middle) with the elution of protein standards of known size (670, 158, 44.0, 16.7 and 1.35 kDa; top), and relative abundance of mesembrenone (**19**) and mesembrine (**20**) in the corresponding *in vitro* assays (bottom). The shaded region marks the active fractions pooled for the next round. (**D**) Round 2, anion-exchange chromatography (AEC) of the pooled active fractions, showing $A_{280}$ and conductivity (top) and assay activity (bottom). (**E**) Round 3, SEC of the pooled active fractions, showing $A_{280}$ (top) and assay activity (bottom). (**F**) LC-MS EIC of mesembrine (**20**) showing the activity assays from fractions collected in round 3 with four fractions spanning the final activity peak (top). The corresponding peptide-spectrum matches (PSM) for KDR-1 identified by proteomic mass spectrometry in each fraction (bottom). (**G**) LC-MS showing the mean of the peak area for EICs of **19** and **20** in *N. benthamiana* leaves expressing the (**19**) gene set with and without *KDR-1*. Bars show mean ± SEM with three biological replicates and GFP is shown as negative control.

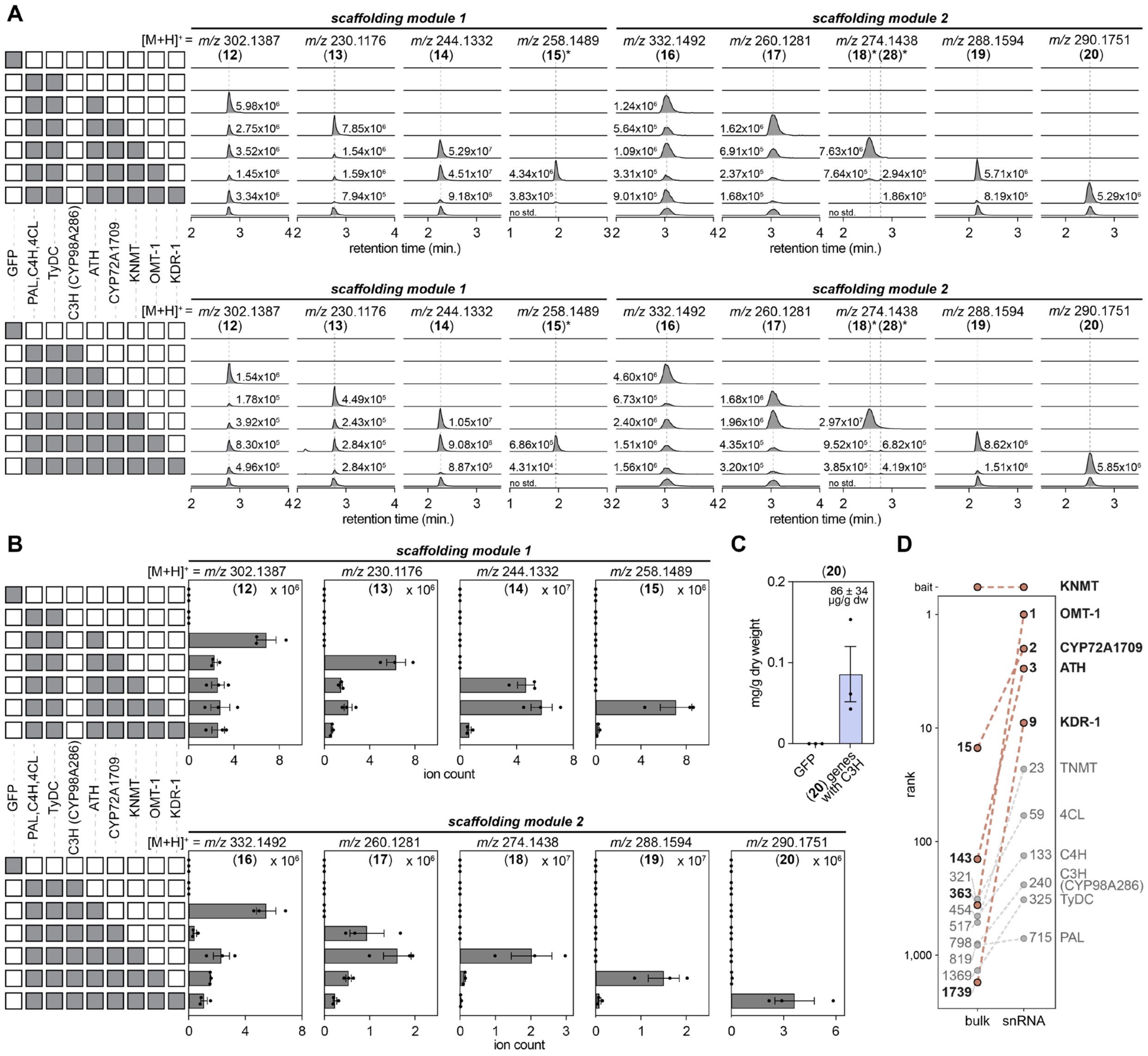


**fig. S11. Transient expression of mesembrine biosynthetic genes in *N. benthamiana*.** (**A**) LC-MS chromatograms of mesembrine (**20**) pathway build-up samples, shown without (top) and with (bottom) C3H (CYP98A286). Peak areas are annotated above each EIC. Columns show scaffolding module 1 (**12** = *m/z* 302.1387; **13** = *m/z* 230.1176; **14**/**27** = *m/z* 244.1332; **15** = *m/z* 258.1489; all $[M+H]^+$) and scaffolding module 2 (**16** = *m/z* 332.1492; **17** = *m/z* 260.1281; **18**/**28** = *m/z* 274.1438; **19** = *m/z* 288.1594; **20** = *m/z* 290.1751; all $[M+H]^+$). An asterisk indicates predicted structures with no purified or authentic standard available, but inferred from LC-MS exact mass, MS/MS fragmentation, and biosynthetic pathway logic. (**B**) Quantification of the mean peak area for EICs of each molecule in scaffolding module 1 (top) and module 2 (bottom), corresponding to the gene combinations in (A). (**C**) Quantification of mesembrine (**20**) in samples containing the **20** gene set with *C3H* (86 ± 34 µg/g dry weight). Bars show mean peak area ± SEM with three biological replicates. (**D**) Rank of each characterized mesembrine (**20**) pathway gene correlation to *KNMT* using either bulk (n = 24 samples; Pearson's *r*) or snRNA (n = 15,629 nuclei; by Spearman rank correlation) co-expression data.

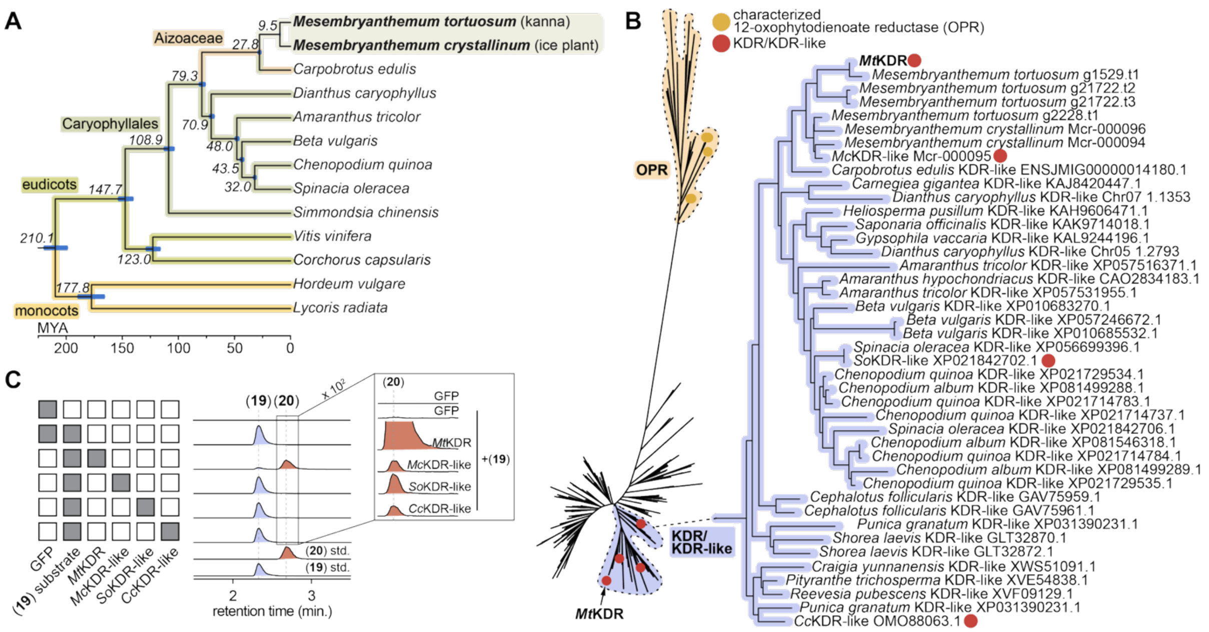


**fig. S12. Assessing the evolutionary history of KDR-1.**
(**A**) Phylogenomic analysis of *M. tortuosum* and 12 select plant species described in **table S1**. Node labels indicate estimated divergence times in million years ago (MYA). *M. tortuosum* and *M. crystallinum* diverged 9.5 MYA. Blue bars indicate 95% confidence intervals for estimated divergence times at each node. (**B**) Maximum-likelihood phylogenetic tree of the KDR/KDR-like clade with characterized 12-oxophytodienoate reductases (OPR) as outgroup. Yellow circles indicate characterized OPRs and red circles indicate the KDR/KDR-like enzymes assayed in panel C. *Mt*KDR is indicated by an arrow. (**C**) *In planta* assay of KDR-1 orthologs using transient expression in *N. benthamiana*. Leaves expressing genes for GFP, *Mt*KDR, *Mc*KDR-like (*M. crystallinum*), *So*KDR-like (*Spinacia oleracea*) or *Cc*KDR-like (*Corchorus capsularis*) were co-infiltrated with mesembrenone (**19**). EICs are shown for **19** ($[M+H]^+$ = *m/z* 288.1594 and **20** ($[M+H]^+$ = *m/z* 290.1751), with authentic standards shown below. Magnified view of the EIC for **20** showing trace activity for all three non-*M. tortuosum* orthologs (right).

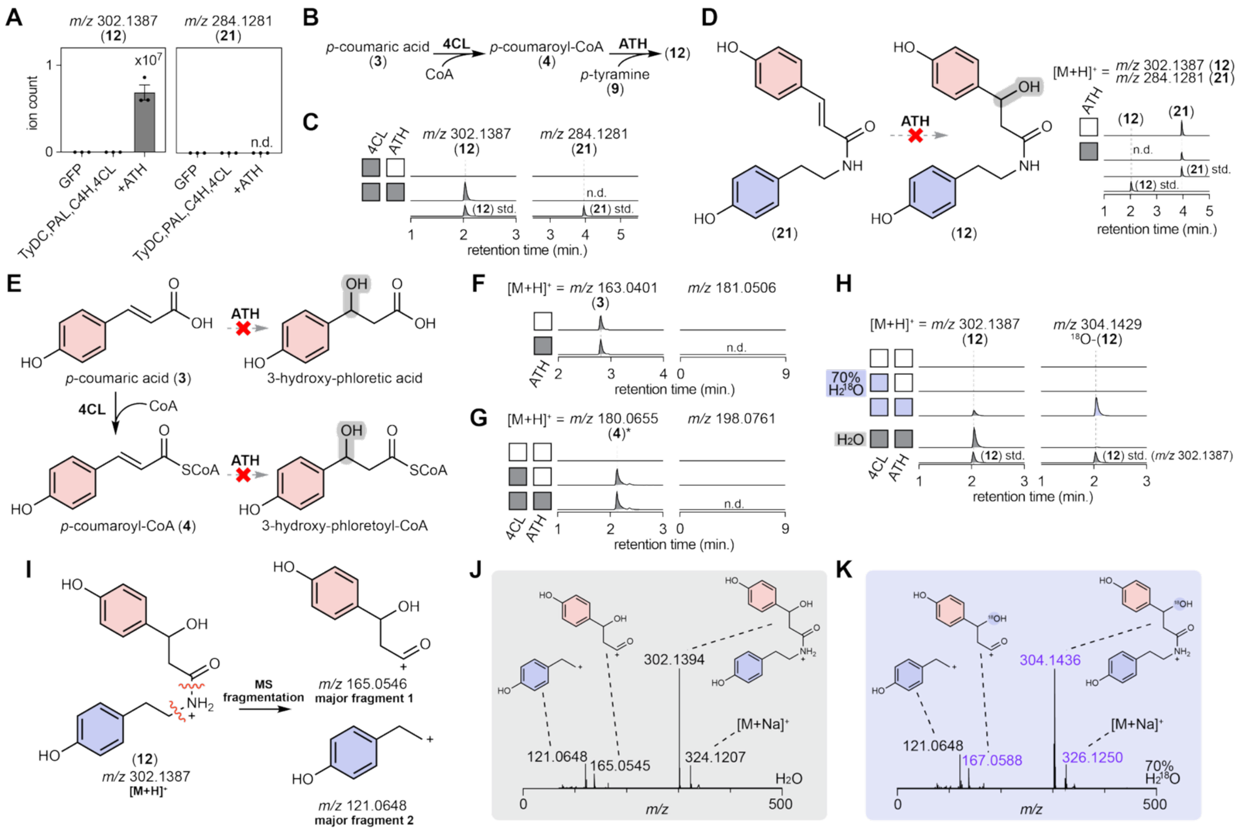


**fig. S13. Characterizing the non-canonical activity of ATH.**
(**A**) Comparison of **12** ($[M+H]^+ = m/z$ 302.1387) and **21** ($[M+H]^+ = m/z$ 284.1281) ion abundance in mesembrine pathway reconstruction in *N. benthamiana*. **21** is not detected (n.d.). (**B**) Schematic of ATH *in vitro* assay set-up. First, 4CL generates *p*-coumaroyl-CoA (**4**) from *p*-coumaric acid (**3**) and CoA. Then, ATH uses **4** and *p*-tyramine (**9**) as substrates to produce **12**. (**C**) *In vitro* assay of ATH produces **12** but not **21**. Authentic standards are shown below each LC-MS EIC. (**D**) ATH does not have activity on **21** directly *in vitro* (combined EICs for **12** and **21** with authentic standards shown). (**E**) Schematic of ATH *in vitro* assay on phenylpropanoid precursors prior to *N*-acyltransferase conjugation. (**F**) ATH does not convert *p*-coumaric acid (**3**; $[M\text{-}H]^- = m/z$ 163.0401) into 3-hydroxy-phloretic acid (expected $[M\text{-}H]^- = m/z$ 181.0506). (**G**) ATH not is observed to act on *p*-coumaroyl-CoA (**4***; $[M+H]^+ = m/z$ 180.0655, derivatized by hydroxylamine) to yield 3-hydroxy-phloretoyl-CoA (expected $[M+H]^+ = m/z$ 198.0761). (**H**) *In vitro* assay in 70% $H_2{}^{18}O$ leads to the production of $^{18}O$-**12** ($[M+H]^+ = m/z$ 304.1429). (**I**) $MS^1$ in-source fragmentation of **12** leading to major observed fragments. (**J**) $MS^1$ spectrum of **12** from ATH *in vitro* assay performed in water. (**K**) $MS^1$ spectrum of $^{18}O$-**12** from ATH *in vitro* assay performed in 70% $H_2{}^{18}O$. The +2 *m/z* shift is retained in phenylpropanoid fragment (*m/z* 167.0588), but not in the *p*-tyramine (**9**)-derived fragment (*m/z* 121.0648), localizing the $^{18}O$ to the β-hydroxy group.

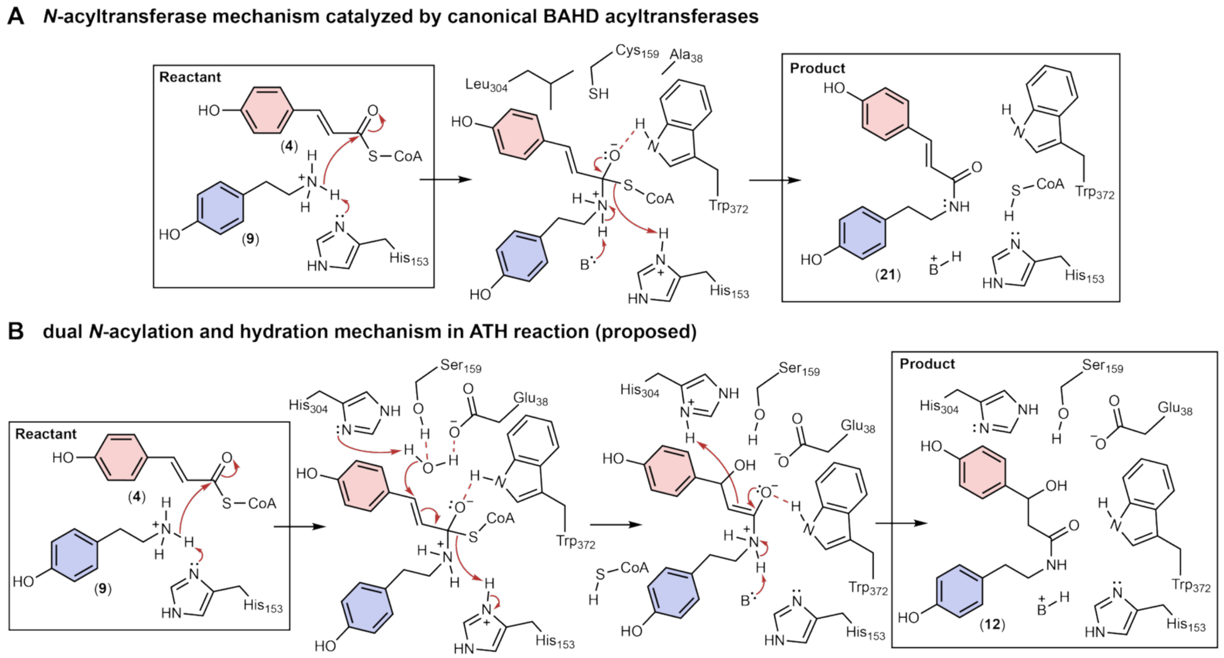


**fig. S14. Proposed mechanism of ATH compared to canonical *N*-acyltransferase mechanism.** (**A**) Canonical *N*-acyltransferase mechanism to conjugate **4** and **9** to yield **21**. The counterpart residues of ATH active-site residues are shown ($Ala_{38}$, $Cys_{159}$, $Leu_{304}$). (**B**) Proposed mechanism of dual *N*-acylation and hydration catalyzed by ATH as illustrated in Figure 4G.

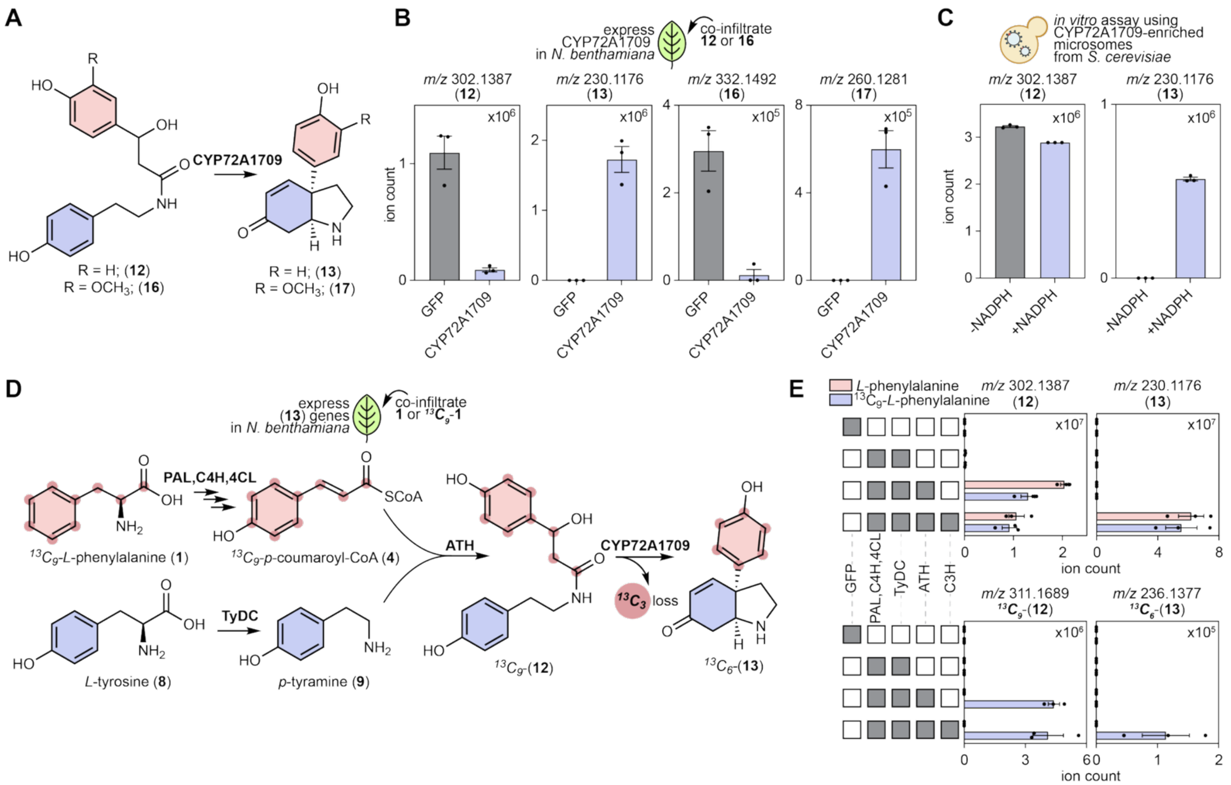


**fig. S15. Characterizing the CYP72A1709 scaffold-forming reaction.**
(**A**) Schematic of the CYP72A1709 scaffolding reaction, which converts the linear amides **12** (R = H) and **16** (R = $OCH_3$) to the cyclized scaffolds **13** and **17**, respectively. (**B**) *In planta* production of **13** or **17** when **12** or **16**, respectively, are co-infiltrated into *N. benthamiana* leaves expressing CYP72A1709 (**12** = *m/z* 302.1387, **13** = *m/z* 230.1176, **16** = *m/z* 332.1492 and **17** = *m/z* 260.1281; all $[M+H]^+$). (**C**) *In vitro* assay using CYP72A1709-enriched microsomes isolated from *S. cerevisiae*, showing NADPH-dependent consumption of **12** and production of **13**. (**D**) $^{13}C_9$-*L*-phenylalanine (**1**) tracing in *N. benthamiana* leaves expressing the biosynthetic genes to produce **13**. The nine $^{13}C$ are shown in gray circles and three are lost during scaffold formation. (**E**) LC-MS of $^{13}C_9$-*L*-phenylalanine (**1**) tracing showing the mean of the peak area for EICs of each molecule. (**12** = *m/z* 302.1387; **13** = *m/z* 230.1176; $^{13}C_9$-**12** = *m/z* 311.1689; $^{13}C_6$-**13** = *m/z* 236.1377; all $[M+H]^+$). Bars show mean ± SEM with three biological replicates.

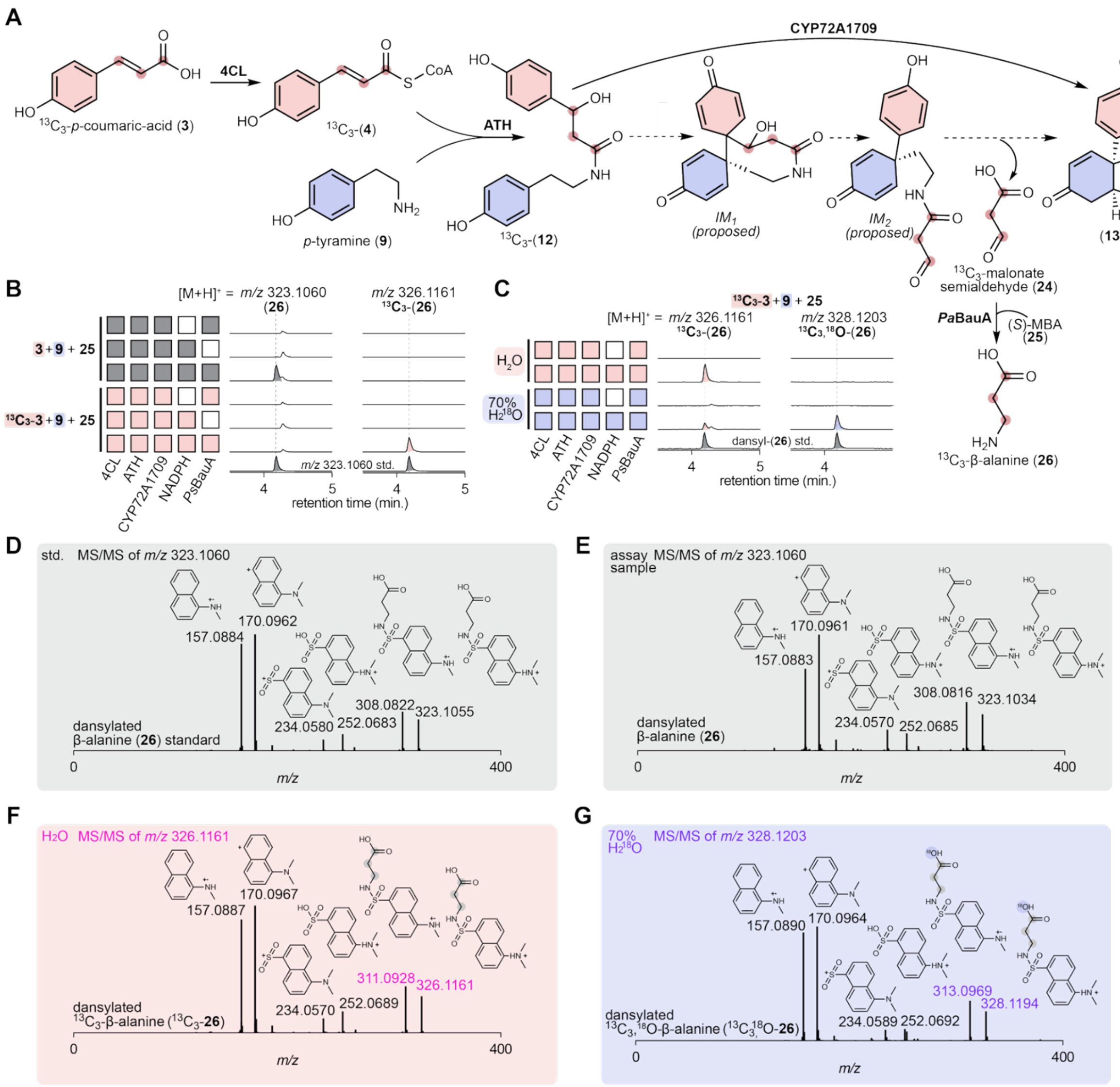


**fig. S16. Capture of malonate semialdehyde from the CYP72A1709 reaction.**
(**A**) Schematic of the coupled assay. 4CL activates $^{13}C_3$-*p*-coumaric acid ($^{13}C_3$-**3**) to $^{13}C_3$-**4**, which ATH condenses with *p*-tyramine (**9**) to give $^{13}C_3$-**12**; CYP72A1709 then converts $^{13}C_3$-**12** to **13** via *IM₁* and *IM₂* with release of $^{13}C_3$-malonate semialdehyde ($^{13}C_3$-**24**), which is captured by *Pa*BauA using (*S*)-α-methylbenzylamine ((*S*)-MBA, **25**) as an amine donor to yield $^{13}C_3$-β-alanine ($^{13}C_3$-**26**). (**B**) EICs of dansylated **26** ($[M+H]^+$ = *m/z* 323.1060) and dansylated $^{13}C_3$-**26** ($[M+H]^+$ = *m/z* 326.1161) from assays containing **3** + **9** + **25** (gray) or $^{13}C_3$-**3** + **9** + **25** (pink). Filled squares indicate the components present (4CL, ATH, CYP72A1709, NADPH, *Pa*BauA). An authentic dansylated β-alanine (**26**) standard is shown below. (**C**) The same $^{13}C_3$ assay performed in $H_2O$ (pink) or 70% $H_2^{18}O$ (blue), showing the +2 *m/z* shift from dansylated $^{13}C_3$-**26** (*m/z* 326.1161) to dansylated $^{13}C_3$,$^{18}$O-**26** (*m/z* 328.1203). (**D**) MS/MS spectrum of *m/z* 323.1060 for the dansylated β-alanine (**26**) standard. (**E**) MS/MS spectrum of *m/z* 323.1060 from the assay sample. (**F**) MS/MS spectrum of *m/z* 326.1161 for dansylated $^{13}C_3$-**26** produced in $H_2O$. (**G**) MS/MS spectrum of *m/z* 328.1203 for dansylated $^{13}C_3$,$^{18}$O-**26** produced in 70% $H_2^{18}O$, confirming that the oxygen incorporated during amide cleavage derives from bulk water.

**A**

| gene | CYP72A1709 r | CYP72A1709 rank | ATH r | ATH rank | characterized function | Pfam annotation | BLASTp annotation | identity % M. crystallinum | identity % A. thaliana |
|---|---|---|---|---|---|---|---|---|---|
| *g19370* | 1.000 | 0 | 0.669 | 1 | CYP72A1709 | Cytochrome P450 | Cytochrome P450 CYP72A219 | 32.2 | 50.4 |
| *g4212* | 0.669 | 1 | 1.000 | 0 | ATH | Transferase family | Pelargonidin 3-O-(6-caffeoylglucoside) 5-O- (6-O-malonylglucoside) 4'''-malonyltransferase | 75.0 | 33.0 |
| *g37187* | 0.603 | 2 | 0.480 | 3 | 4CL | AMP-binding enzyme | 4-coumarate--CoA ligase 1 | 97.3 | 68.2 |
| ⋮ | | | | | | | | | |
| *g29247* | 0.549 | 5 | 0.466 | 4 | C4H | Cytochrome P450 | Trans-cinnamate 4-monooxygenase | 97.6 | 84.5 |
| *g272* | 0.542 | 6 | 0.567 | 2 | OMT-1 | O-methyltransferase domain | Cathecol O-methyltransferase 1 | 78.0 | 54.2 |
| ⋮ | | | | | | | | | |
| *g33715* | 0.516 | 9 | 0.416 | 11 | putative dehydrogenase (ALDH-like-1) | Aldehyde dehydrogenase family | Methylmalonate-semialdehyde dehydrogenase [acylating] | 99.0 | 85.2 |
| ⋮ | | | | | | | | | |
| *g20977* | 0.494 | 13 | 0.351 | 30 | TNMT | Mycolic acid cyclopropane synthetase | (S)-coclaurine N-methyltransferase | 76.7 | 54.7 |
| ⋮ | | | | | | | | | |
| *g22226* | 0.450 | 22 | 0.344 | 33 | C3H (CYP98A286) | Cytochrome P450 | Cytochrome P450 98A2 | 97.7 | 68.5 |
| ⋮ | | | | | | | | | |
| *g20975* | 0.350 | 55 | 0.342 | 34 | KNMT | Mycolic acid cyclopropane synthetase | (S)-coclaurine N-methyltransferase | 74.4 | 55.2 |
| ⋮ | | | | | | | | | |
| *g35047* | 0.339 | 63 | 0.318 | 46 | TyDC | Pyridoxal-dependent decarboxylase conserved domain | Tyrosine decarboxylase | 98.5 | 53.4 |
| ⋮ | | | | | | | | | |
| *g30732* | 0.302 | 103 | 0.282 | 69 | PAL | Aromatic amino acid lyase | Phenylalanine ammonia-lyase | 99.3 | 85.5 |
| ⋮ | | | | | | | | | |
| *g20979* | 0.139 | 671 | 0.071 | 1,820 | NMT-like2 | Mycolic acid cyclopropane synthetase | (S)-coclaurine N-methyltransferase | 65.2 | 51.9 |
| ⋮ | | | | | | | | | |
| *g1529* | -0.007 | 19,524 | 0.034 | 4,585 | KDR-1 | NADH:flavin oxidoreductase / NADH oxidase family | Putative 12-oxophytodienoate reductase 11 | 87.3 | 74.7 |
| ⋮ | | | | | | | | | |
| *g20969* | -0.213 | 25,887 | -0.029 | 23,079 | NMT-like1 | Mycolic acid cyclopropane synthetase | (S)-coclaurine N-methyltransferase | 63.0 | 51.6 |

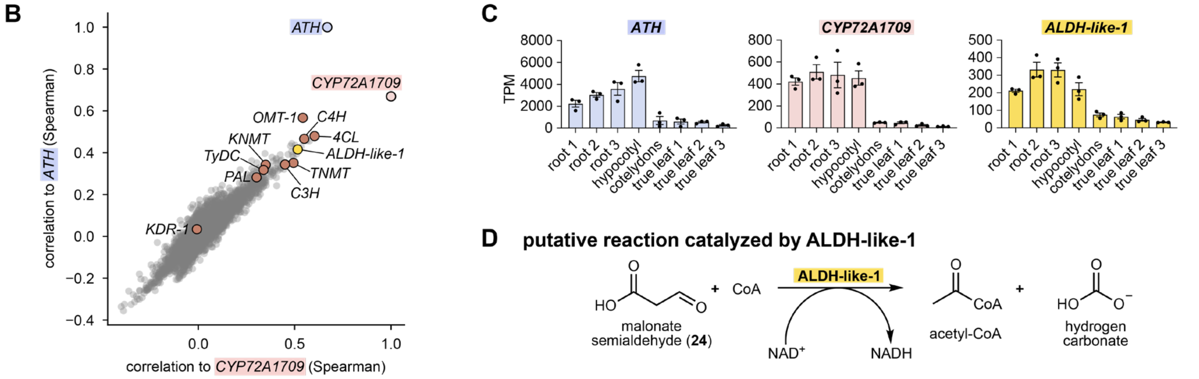


**fig. S17. snRNA-seq co-expression analysis using *CYP72A1709* and *ATH* as bait genes.**
(**A**) Table of mesembrine (**20**) biosynthetic genes ranked according to Spearman's correlation to *CYP72A1709* expression across 15,629 nuclei. Also shown in this table is the corresponding rank to *ATH* expression, characterized gene function, Pfam and BLASTp annotation, and amino acid identity (%) to the closest orthologs in *M. crystallinum* and *A. thaliana*. (**B**) Co-expression plot showing the correlation of *M. tortuosum* genes to *CYP72A1709* and *ATH*. Spearman correlation coefficients are used. Orange dots indicate mesembrine (**20**) biosynthetic genes and yellow dot indicate a putative dehydrogenase gene (*ALDH-like-1*, *g33715*) with a functional annotation to *Methylmalonate-semialdehyde dehydrogenase [acylating]* (*45–47*, *102*). (**C**) Transcript expression (TPM) of *ATH*, *CYP72A1709* and *ALDH-like-1* across *M. tortuosum* tissues shown in Figure 1C, demonstrating co-enrichment in root and hypocotyl tissue. Bars show mean ± SEM with individual replicates overlaid. (**D**) Putative reaction catalyzed by ALDH-like-1, converting malonate semialdehyde (**24**) and CoA substrates to acetyl-CoA and hydrogen carbonate with coupled reduction of $NAD^+$ to NADH.

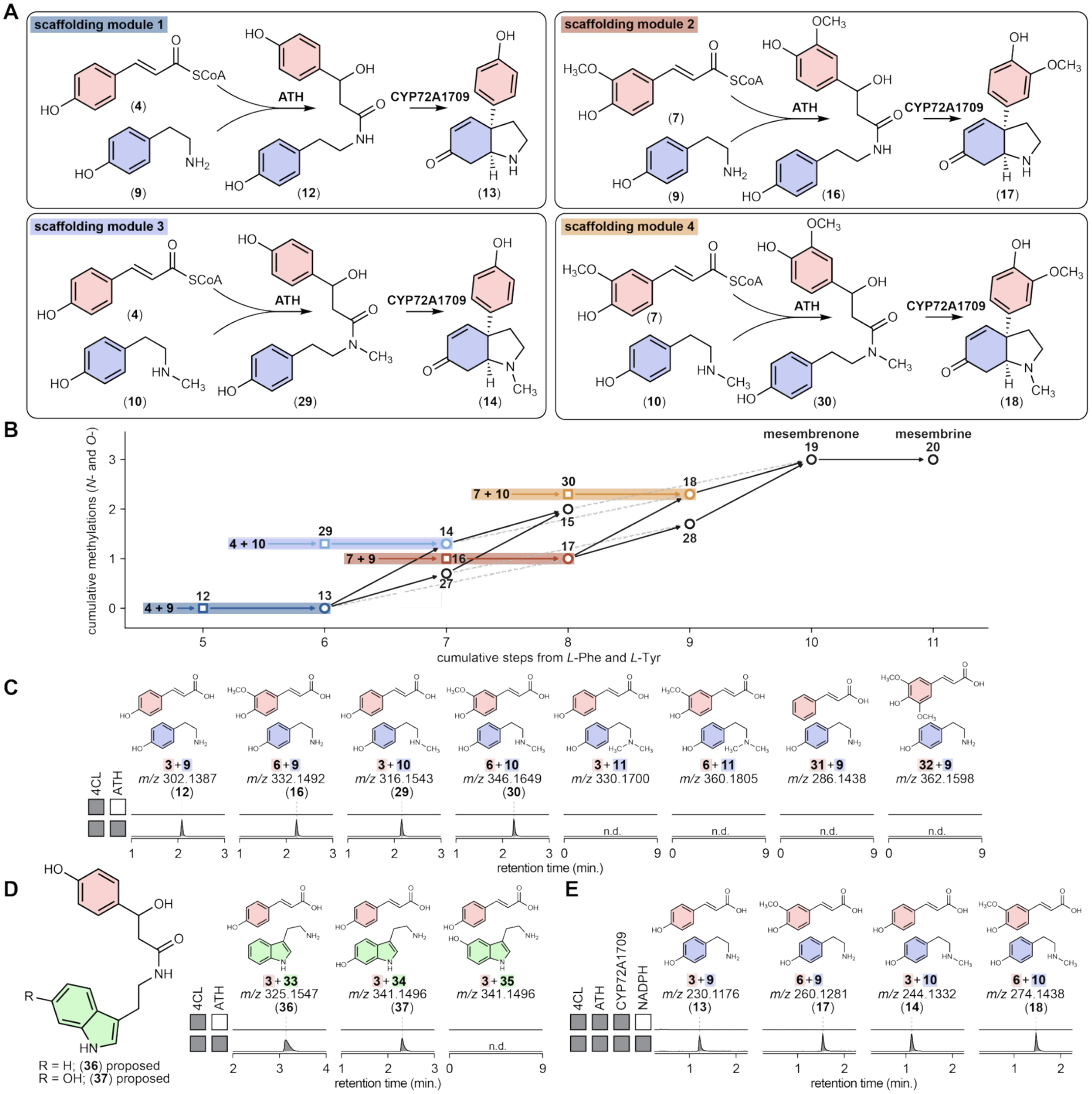


**fig. S18. Metabolic network of ATH and CYP72A1709 scaffolding reactions in Kanna alkaloid biosynthesis.** (**A**) Schematic of four possible scaffolding modules catalyzed by ATH and CYP72A1709 *in vitro*: **4** + **9** → **12** → **13** (module 1); **7** + **9** → **16** → **17** (module 2); **4** + **10** → **29** (proposed) → sceletenone (**14**) (module 3); **7** + **10** → **30** (proposed) → **18** (proposed) (module 4). (**B**) Metabolic network map plotting cumulative *N*- and *O*-methylations against cumulative enzymatic steps from *L*-phenylalanine (**1**) and *L*-tyrosine (**8**), showing the multiple routes by which the four modules converge on mesembrenone (**19**) and mesembrine (**20**). Shaded bars indicate the ATH and CYP72A1709 scaffolding steps for each acyl-donor and amine pair. (**C**) EICs of *in vitro* assays using various *L*-phenylalanine (**1**)- and *L*-tyrosine (**8**)-derived precursors with 4CL and ATH. Products detected: **3** + **9** → **12** (*m/z* 302.1387), **6** + **9** → **16** (*m/z* 332.1492), **3** + **10** → **29** (proposed) (*m/z* 316.1543), **6** + **10** → **30** (proposed) (*m/z* 346.1649). No product was detected for **3** + **11** (expected *m/z* 330.1700), **6** + **11** (*m/z* 360.1805), **31** + **9** (*m/z* 286.1438) or **32** + **9** (*m/z* 362.1598), all predicted $[M+H]^+$. These data indicate that ATH does not accept the tertiary amine hordenine (**11**) or the acyl donors (cinnamic acid, **31** and sinapic acid, **32**). (**D**) EICs of *in vitro* assays using indoleamine acceptors with 4CL and ATH. Proposed product structures (**36** and **37**) are shown on the left. Assay using **3** with tryptamine (**33**) and 6-hydroxytryptamine (**34**) yield the corresponding hydrated amides ($[M+H]^+$ = *m/z* 325.1547 (**36**; proposed) and 341.1496 (**37**; proposed)), whereas 5-hydroxytryptamine (a.k.a. serotonin; **35**) gives no detectable product (*m/z* 341.1496). (**E**) EICs of *in vitro* assays using 4CL, ATH, CYP72A1709-enriched microsomes and NADPH, showing scaffold formation for all four acyl-donor and amine pairs: **3** + **9** → **13** ($[M+H]^+$ = *m/z* 230.1176), **6** + **9** → **17** ($[M+H]^+$ = *m/z* 260.1281), **3** + **10** → **14** ($[M+H]^+$ = *m/z* 244.1332), **6** + **10** → **18** (proposed) ($[M+H]^+$ = *m/z* 274.1438).

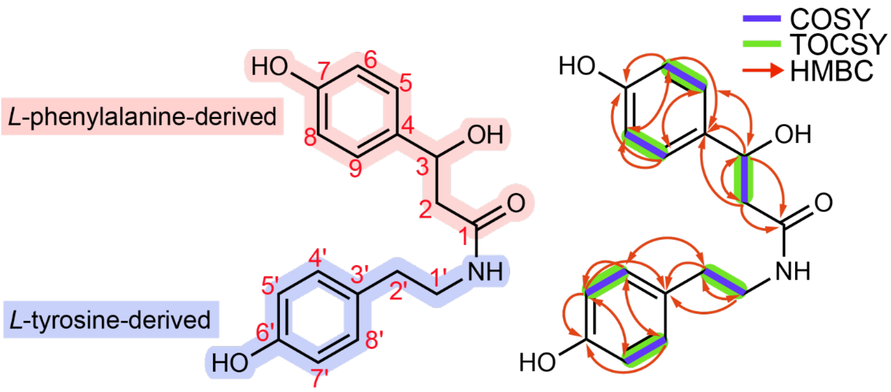


(**12**)
(*proposed*)

| carbon | | 13C δ (ppm) | 1H δ (ppm, J in Hz) |
|---|---|---|---|
| **1** | C | 173.42 | - |
| **2** | $CH_2$ | 40.80 | 2.48 (dd, J = 15.9, 10.1)<br>2.25 (dd, J = 15.9, 4.2) |
| **3** | CH | 58.61 | 4.00 (dd, J = 10.1, 4.2) |
| **4** | C | 129.61 | - |
| **5** | CH | 128.74 | 7.18 (d, J = 8.4) |
| **6** | CH | 115.35 | 6.72 (d, J = 8.4) |
| **7** | C | 157.26 | - |
| **8** | CH | 115.35 | 6.72 (d, J = 8.4) |
| **9** | CH | 128.74 | 7.18 (d, J = 8.4) |
| **1'** | $CH_2$ | 46.79 | 2.60 (m) |
| **2'** | $CH_2$ | 33.18 | 2.62 (m) |
| **3'** | C | 128.69 | - |
| **4'** | CH | 129.50 | 6.93 (d, J = 8.4) |
| **5'** | CH | 115.31 | 6.64 (d, J = 8.4) |
| **6'** | C | 155.95 | - |
| **7'** | CH | 115.31 | 6.64 (d, J = 8.4) |
| **8'** | CH | 129.50 | 6.93 (d, J = 8.4) |

**fig. S19. NMR assignment of 12 ($[M+H]^+$ = *m/z* 302.1387).** Assignments listed in the table above are based on the NMR spectra shown in figs. S20-S26. Final $^1H$ δ assignments are based on experiments collected in deuterated DMSO (DMSO-$d_6$). All experiments were recorded at 600 MHz in DMSO-$d_6$ unless specified.

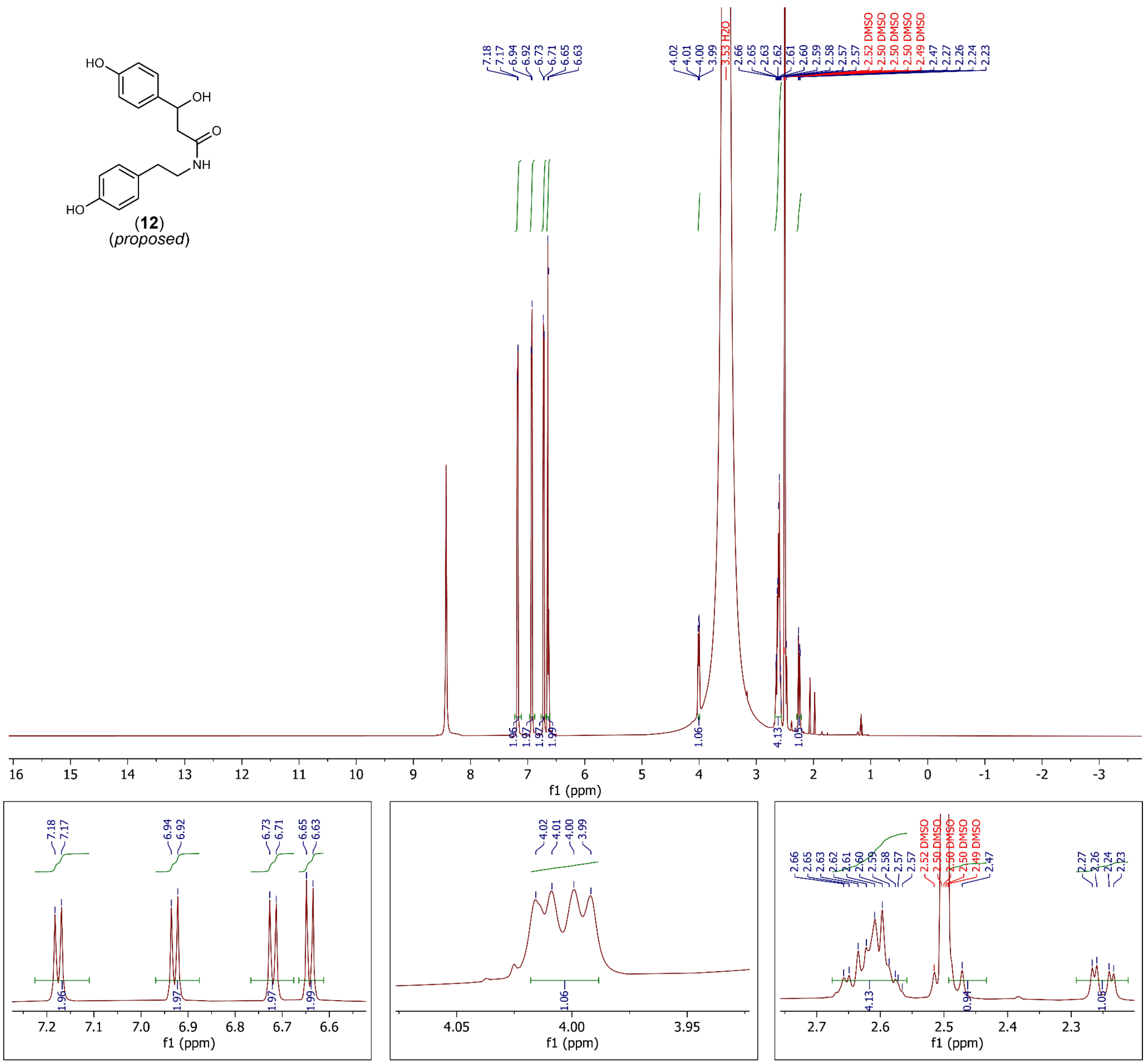


**fig. S20. $^1$H-NMR spectrum of 12 ([M+H]$^+$ = *m/z* 302.1387).** Spectrum collected in DMSO-$d_6$, 600 MHz, referenced to DMSO-$d_5$ = 2.50 ppm. Ammonium formate salt contamination is detected at 8.4277 ppm.

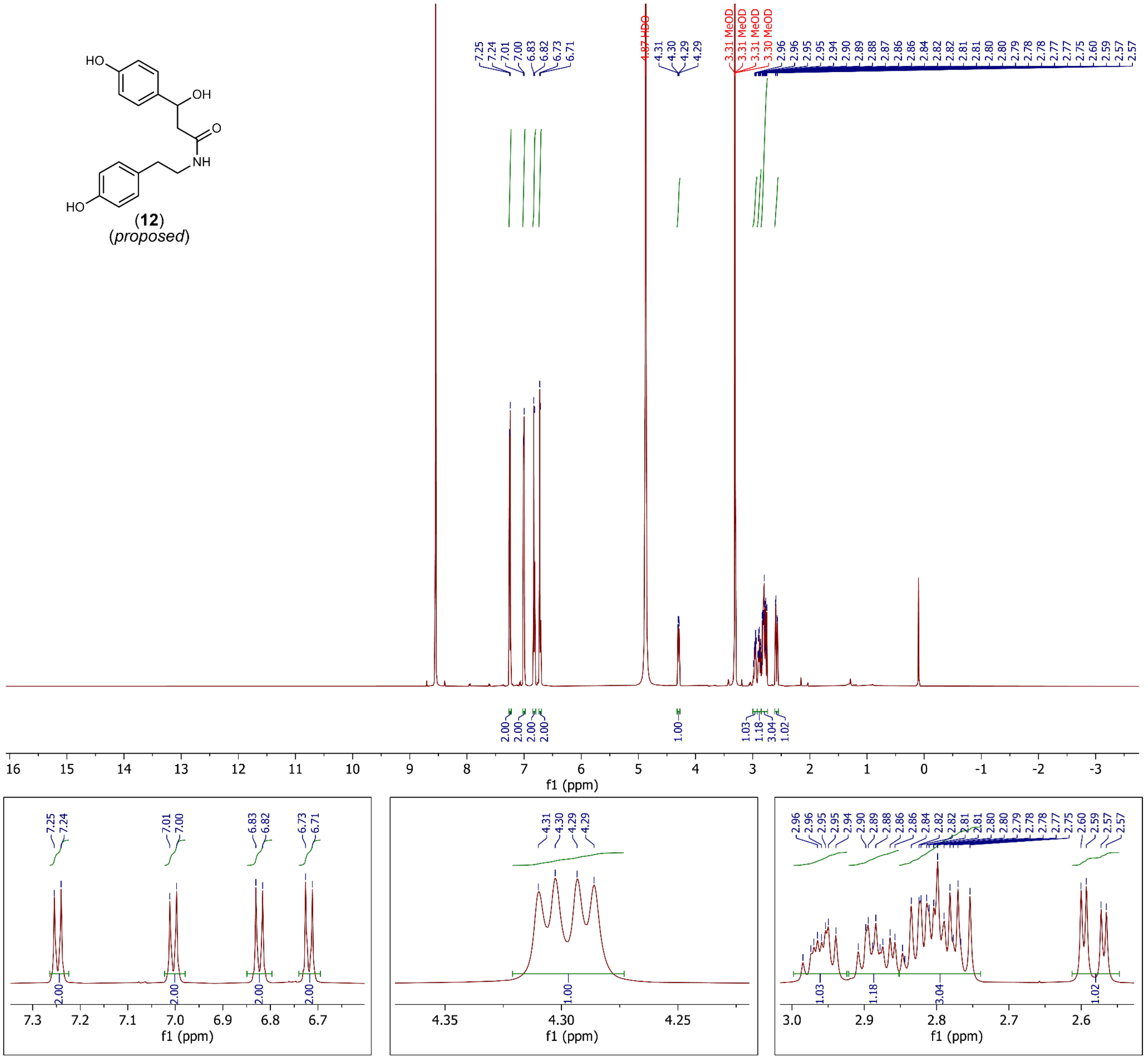


**fig. S21. ¹H-NMR spectrum of 12 ([M+H]⁺ = *m/z* 302.1387).** Spectrum collected in methanol-*d*₄, 600 MHz, referenced to methanol-*d*₃ = 3.31 ppm. Ammonium formate salt contamination is detected at 8.5496 ppm.

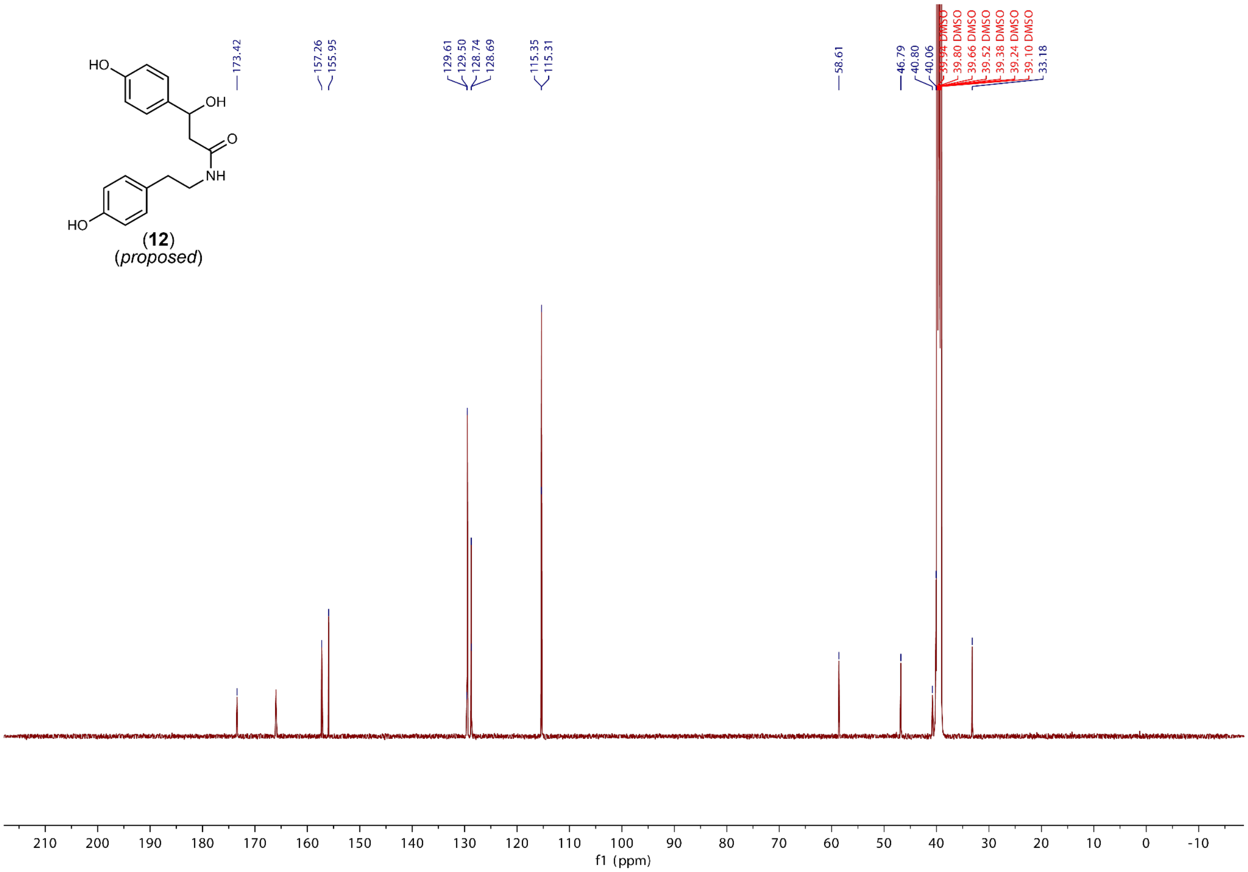


**fig. S22. $^{13}$C-NMR spectrum of 12 ($[M+H]^+$ = *m/z* 302.1387).** Ammonium formate salt contamination is detected at 165.9832 ppm.

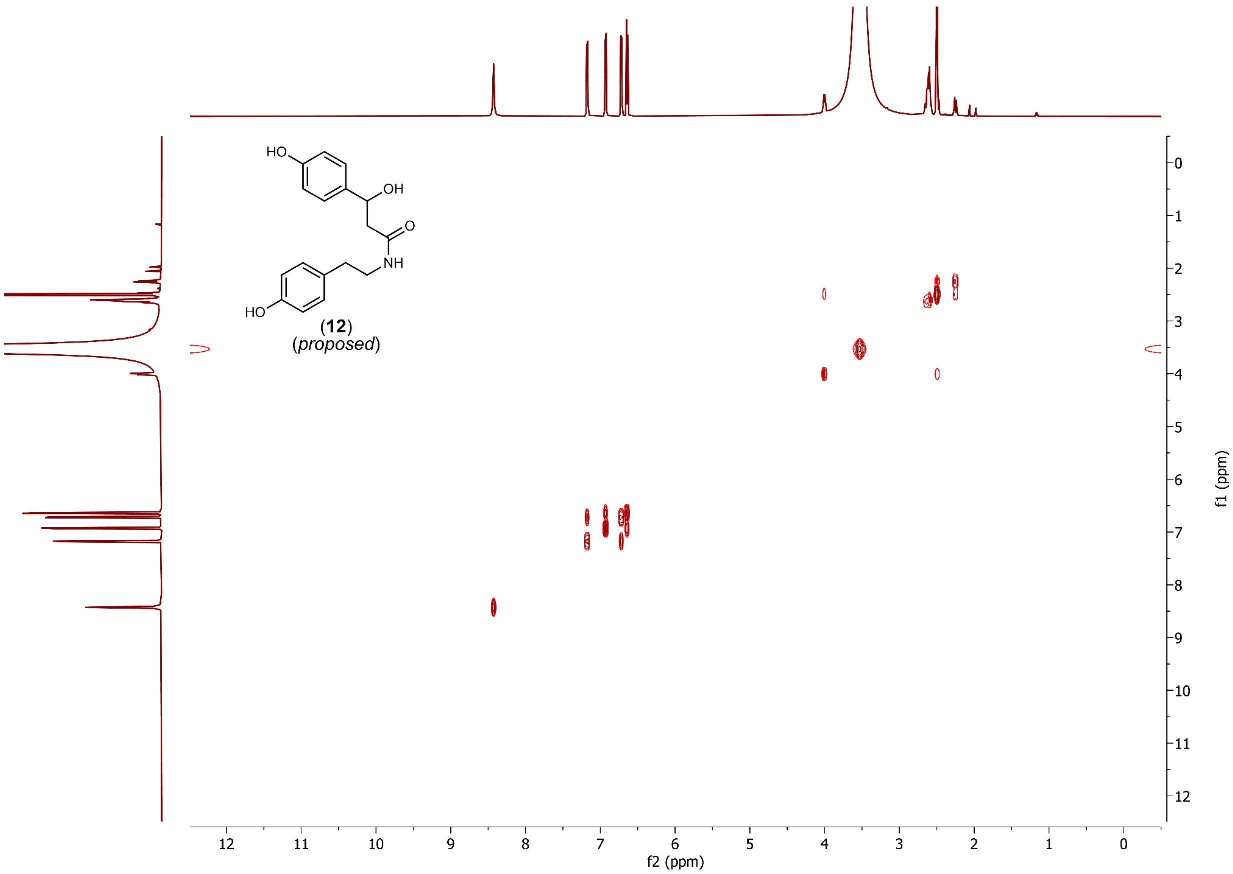


**fig. S23. COSY spectrum of 12 ([M+H]$^+$ = *m/z* 302.1387).**

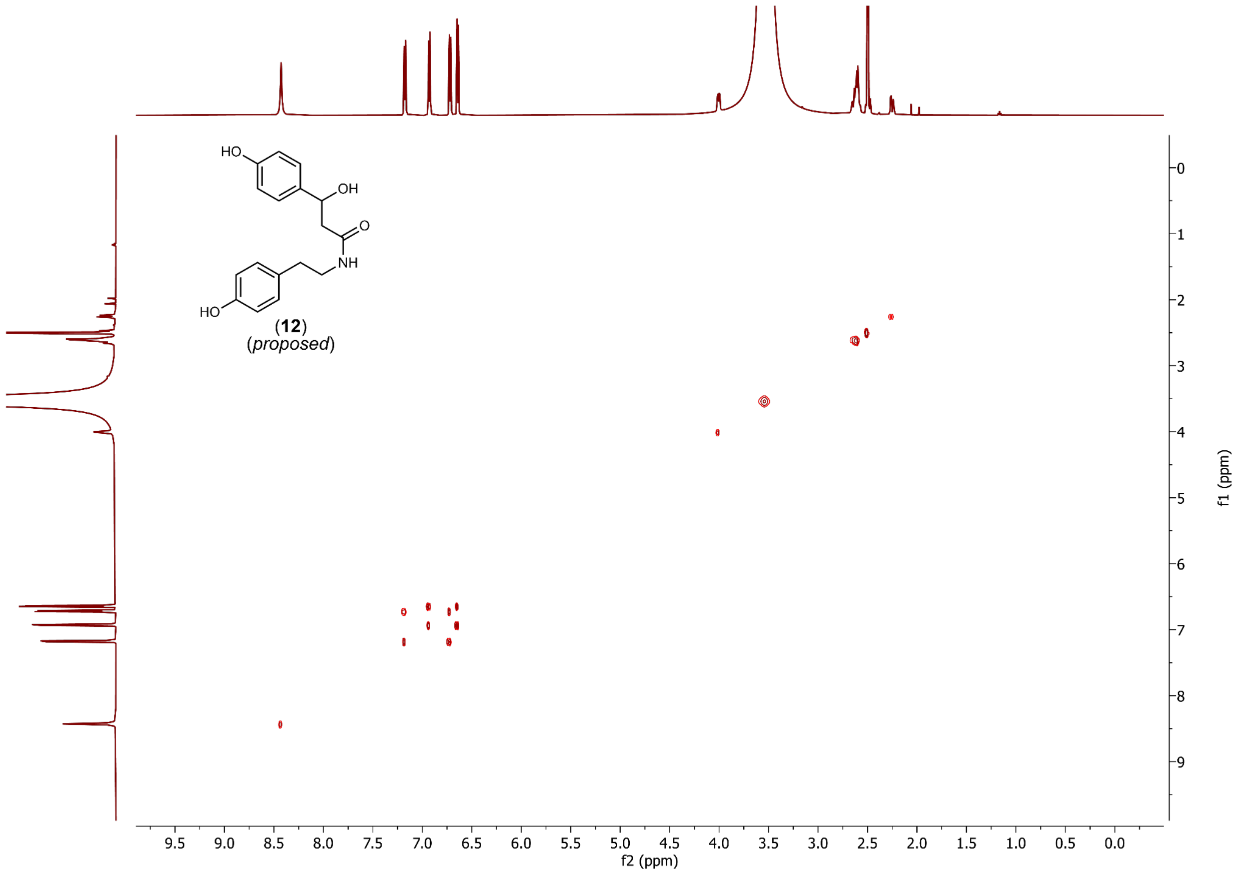


1556

1557 **fig. S24. TOCSY spectrum of 12 ([M+H]$^+$ = *m/z* 302.1387).**

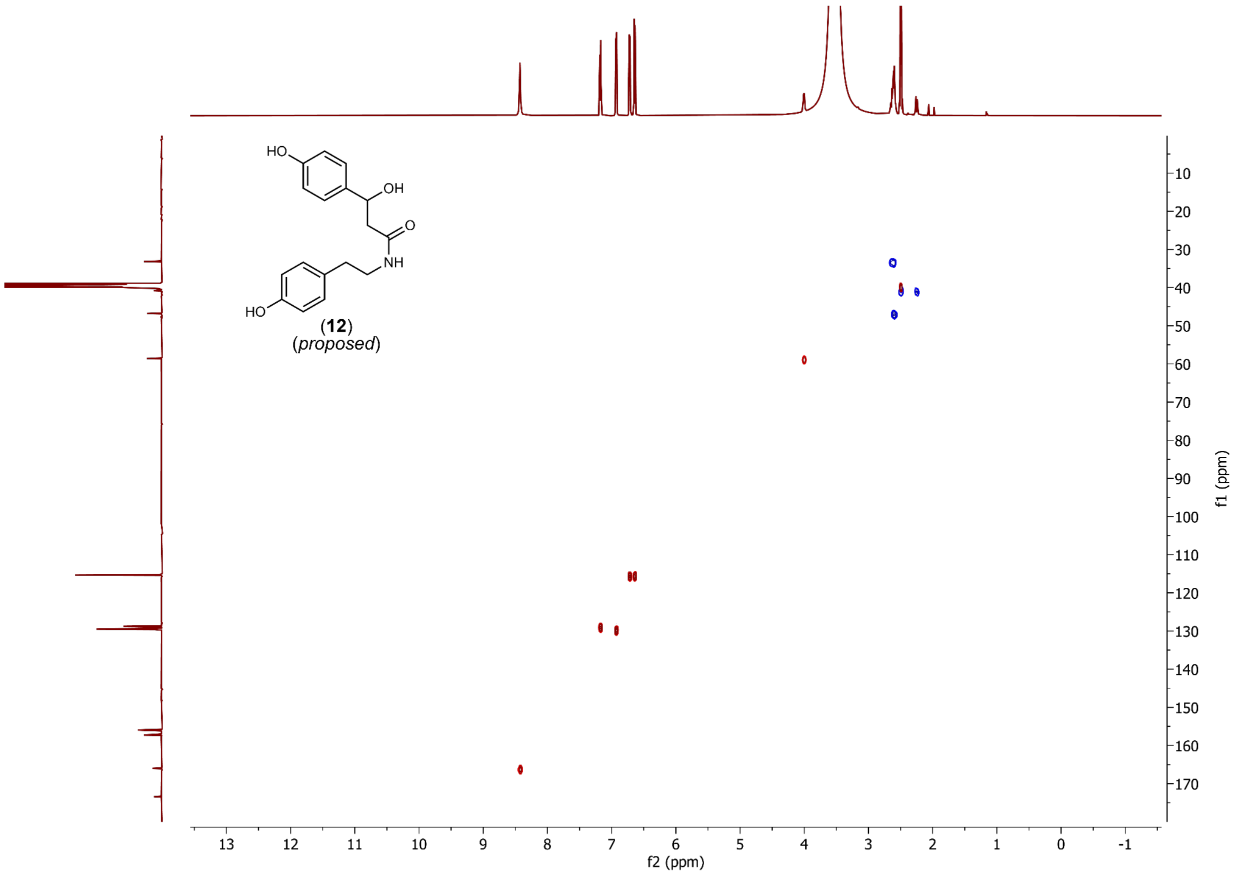


**fig. S25. HSQC spectrum of 12 ($[M+H]^+$ = *m/z* 302.1387).**

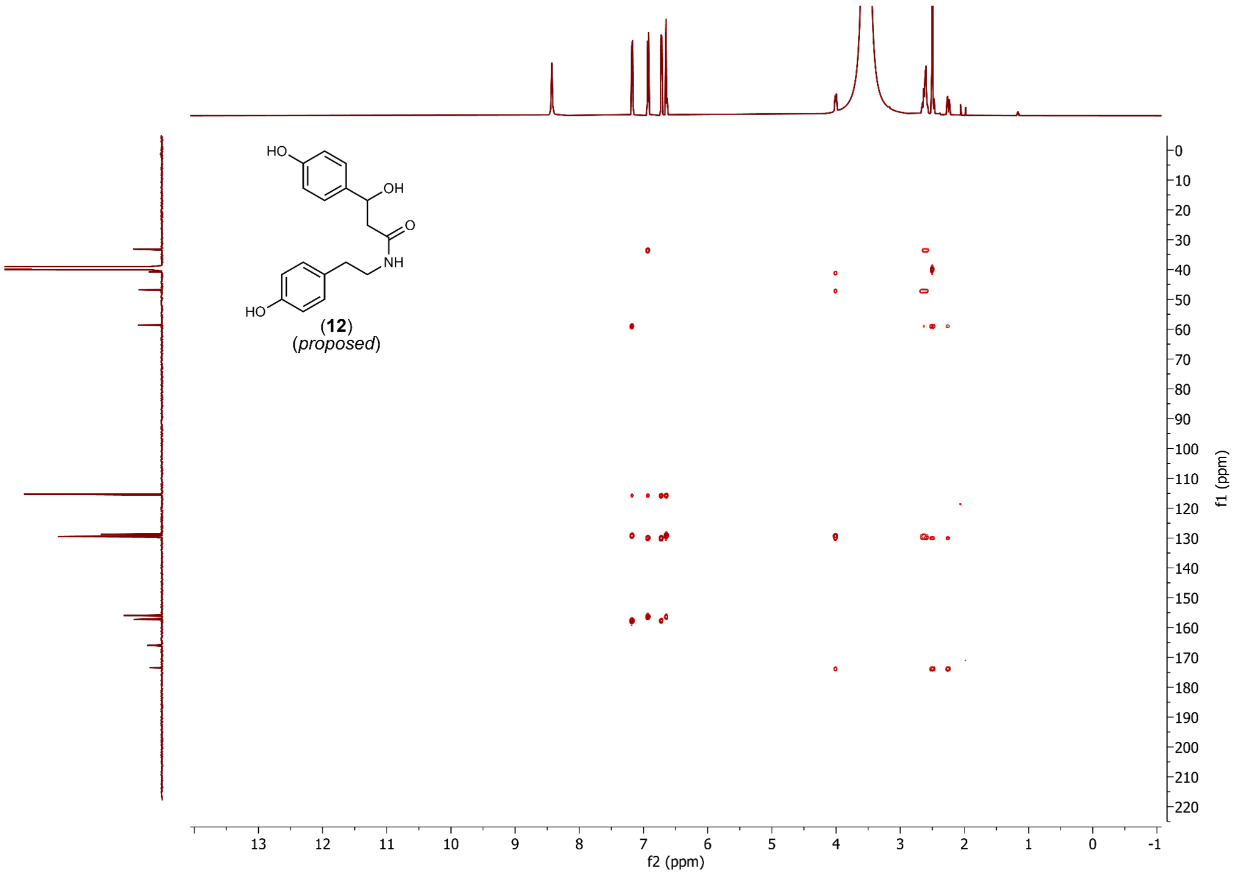


**fig. S26. HMBC spectrum of 12 ([M+H]$^{+}$ = *m/z* 302.1387).**

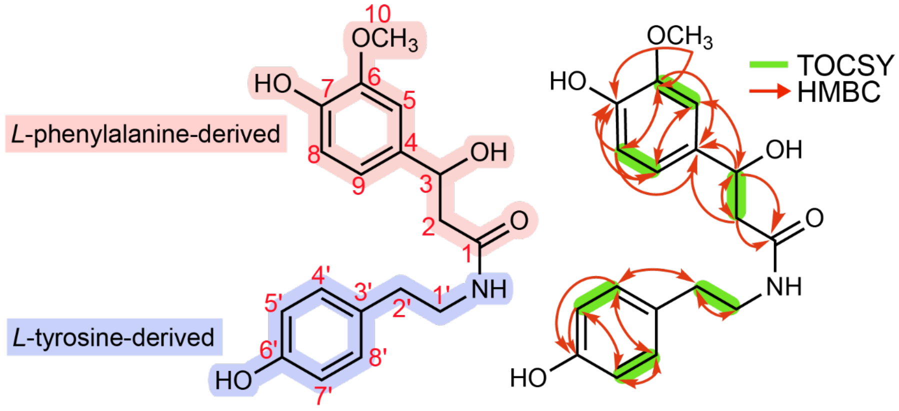


(**16**)
(*proposed*)

| carbon | | 13C δ (ppm) | 1H δ (ppm, J in Hz) |
|---|---|---|---|
| **1** | C | 178.10 | - |
| **2** | CH2 | 41.66 | 2.755 (dd, J = 16.5, 9.8)<br>2.587 (dd, J = 16.5, 4.5) |
| **3** | CH | 61.57 | 4.268 (dd, J = 9.7, 4.4) |
| **4** | C | 128.99 | - |
| **5** | CH | 112.17 | 6.982 (d, J = 1.8) |
| **6** | C | 149.49 | - |
| **7** | C | 148.46 | - |
| **8** | CH | 116.57 | 6.814 (d, J = 8.1) |
| **9** | CH | 121.92 | 6.852 (dd, J = 8.1, 1.9) |
| **10** | OCH3 | 56.44 | 3.857 (s) |
| **1'** | CH2 | 48.04 | 2.948 (m)<br>2.882 (m) |
| **2'** | CH2 | 33.50 | 2.819 (m) |
| **3'** | C | 129.13 | - |
| **4'** | CH | 130.75 | 7.004 (d, J = 8.4) |
| **5'** | CH | 116.65 | 6.717 (d, J = 8.5) |
| **6'** | C | 157.53 | - |
| **7'** | CH | 116.65 | 6.717 (d, J = 8.5) |
| **8'** | CH | 130.69 | 7.004 (d, J = 8.4) |

**fig. S27. NMR assignment of 16 ([M+H]$^+$ = *m/z* 332.1492).** Assignments, as listed in the table above are based on the NMR spectra shown in figs. S28-S32. All experiments were recorded at 600 MHz in deuterated methanol (methanol-$d_4$,) unless specified.

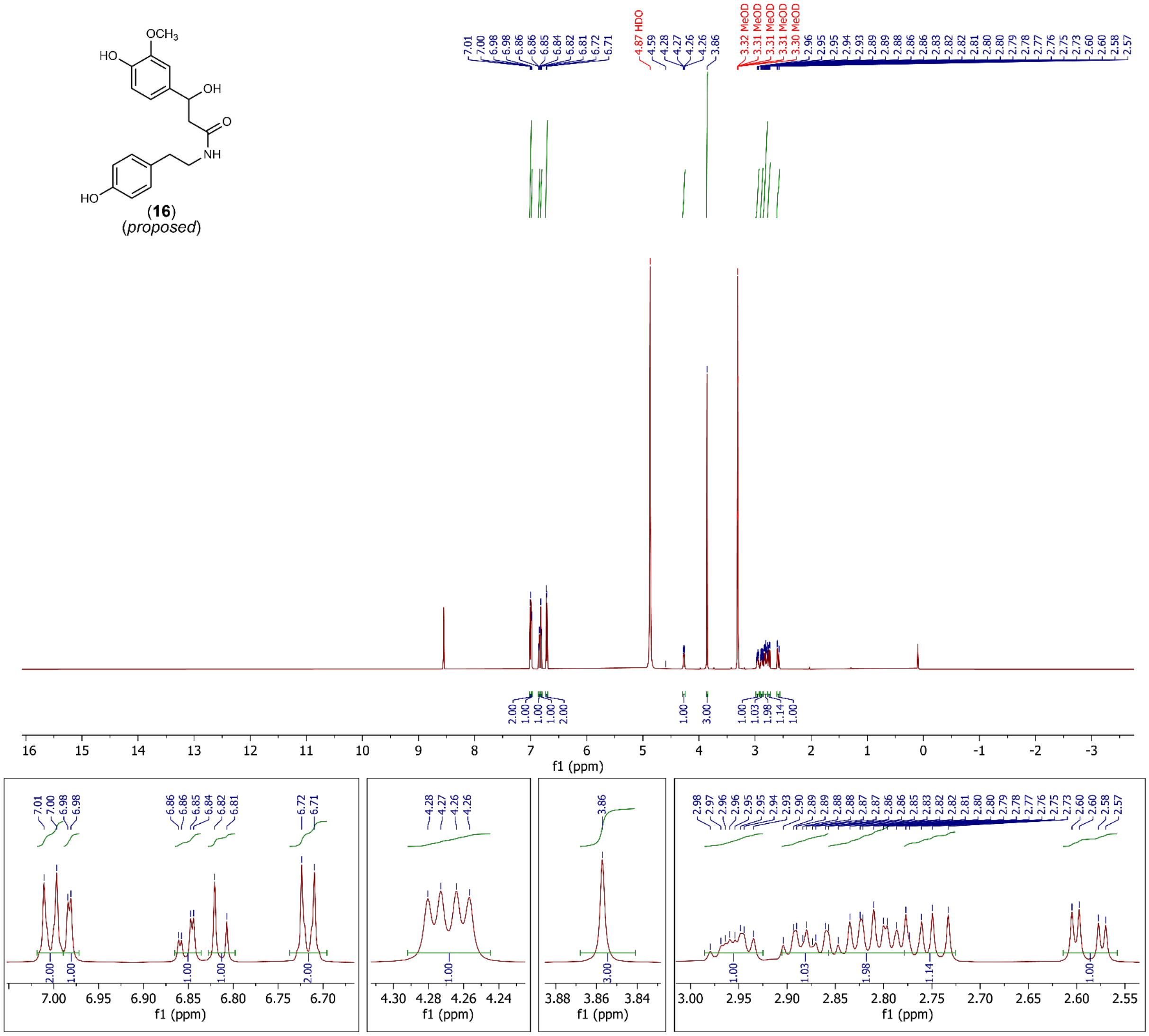


**fig. S28. $^1$H-NMR spectrum of 16 ($[M+H]^+$ = *m/z* 332.1492).** Spectrum collected in methanol-$d_4$, 600 MHz, referenced to methanol-$d_3$ = 3.31 ppm. Ammonium formate salt contamination is detected at 8.5511 ppm.

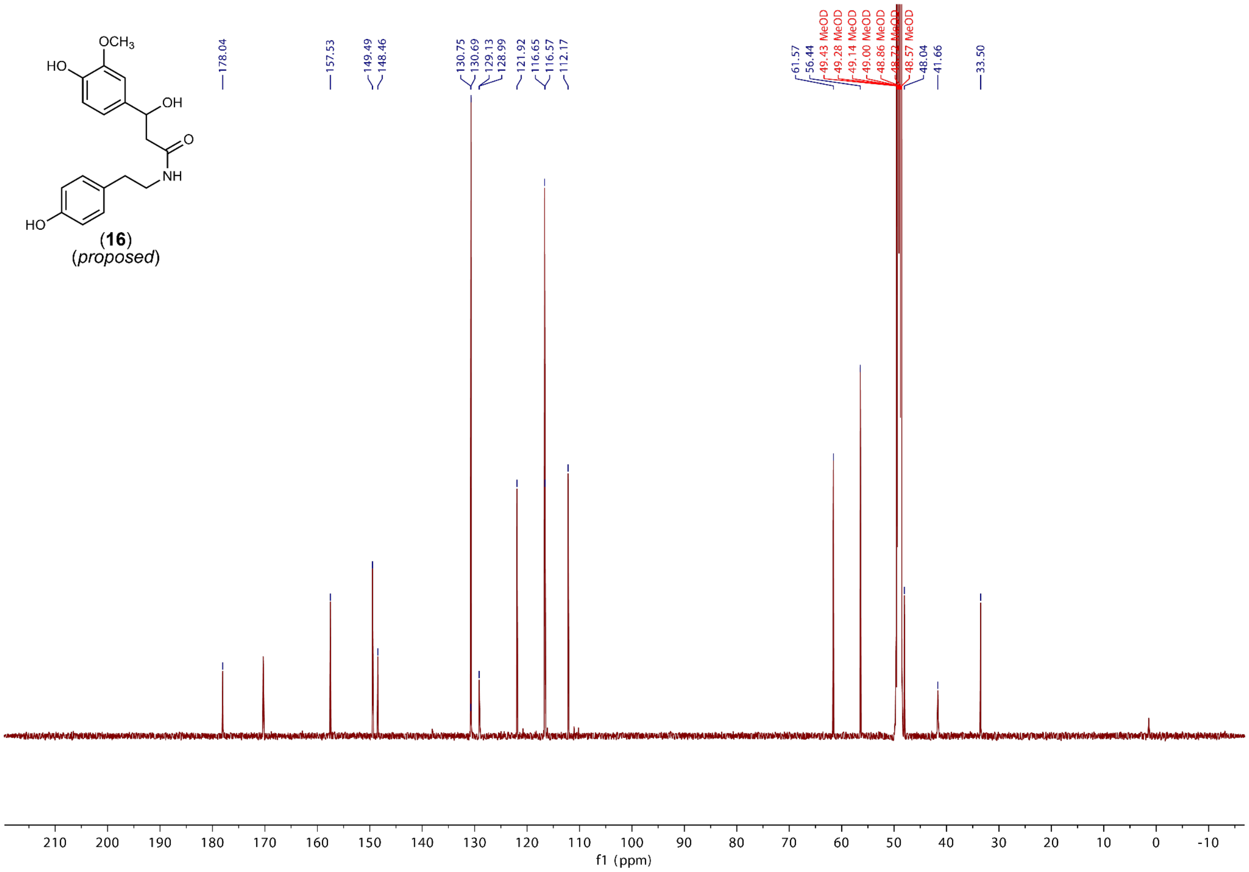


**fig. S29. $^{13}$C-NMR spectrum of 16 ([M+H]$^+$ = *m/z* 332.1492).** Spectrum collected in methanol-$d_4$, 600 MHz, referenced to methanol-$d_3$ = 49.00 ppm. Ammonium formate salt contamination is detected at 170.3243 ppm.

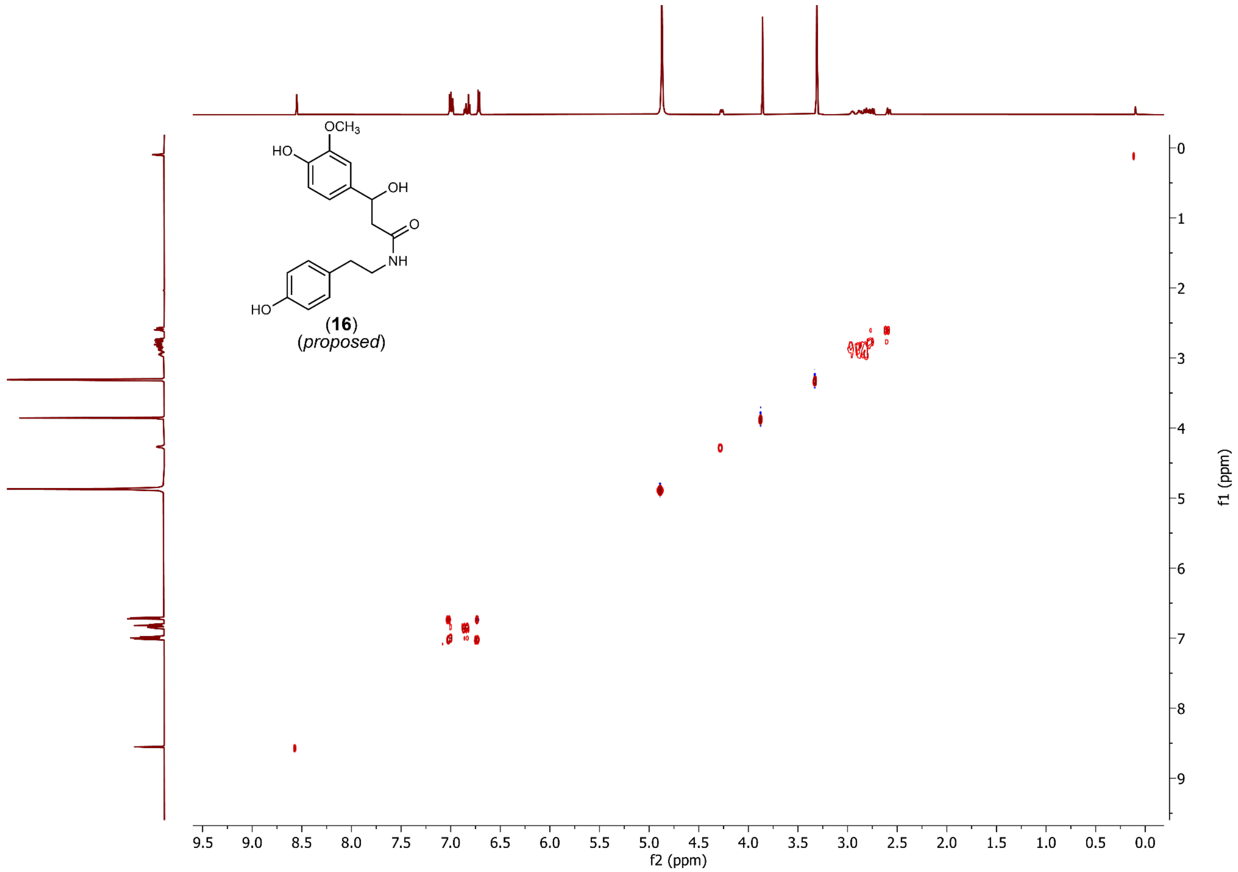


**fig. S30. TOCSY spectrum of 16 ([M+H]$^+$ = *m/z* 332.1492).** Spectrum collected in methanol-$d_4$, 600 MHz.

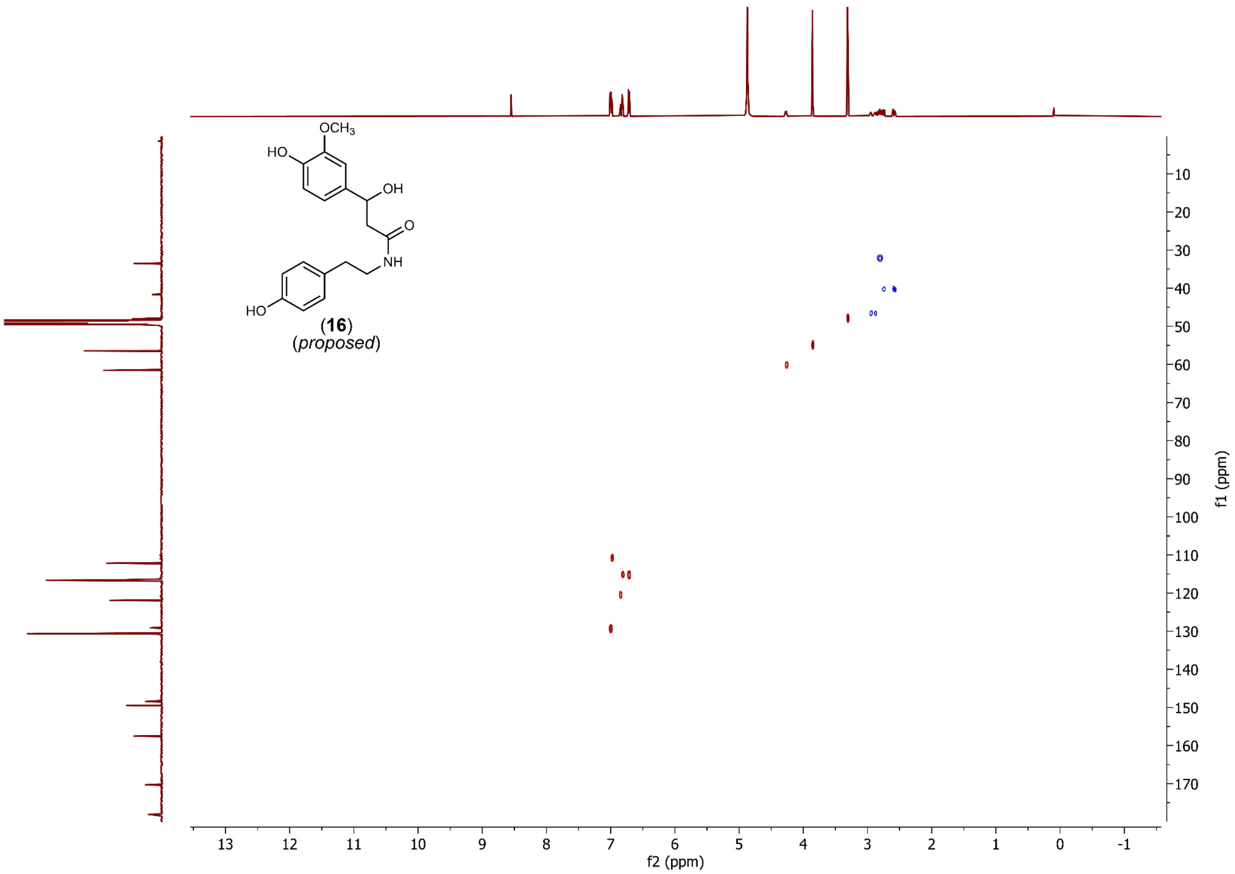


**fig. S31. HSQC spectrum of 16 ([M+H]$^+$ = *m/z* 332.1492).** Spectrum collected in methanol-$d_4$, 600 MHz.

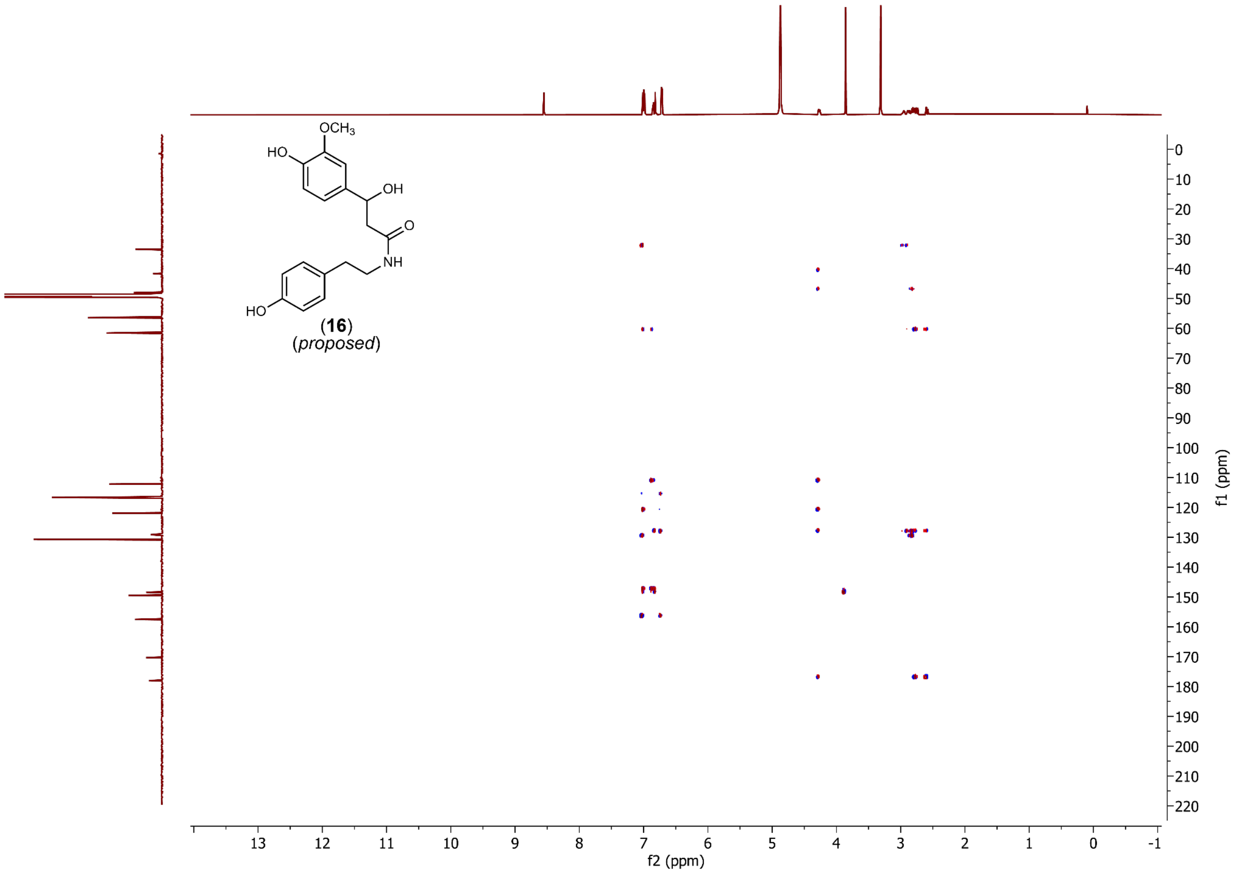


**fig. S32. HMBC spectrum of 16 ($[M+H]^+$ = *m/z* 332.1492).** Spectrum collected in methanol-$d_4$, 600 MHz.

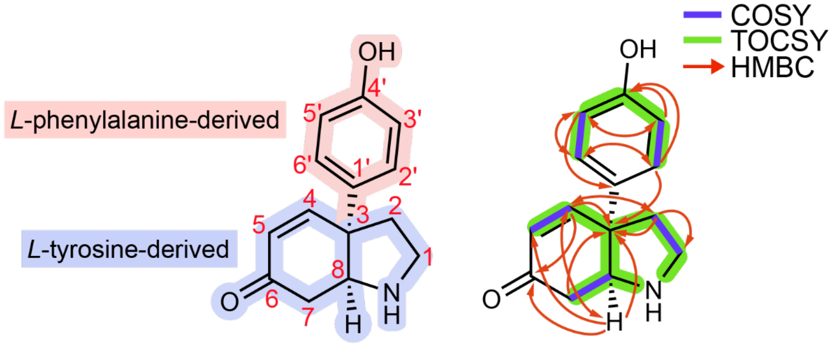


(**13**)
(*proposed*)

| carbon | | 13C δ (ppm) | 1H δ (ppm, J in Hz) |
|---|---|---|---|
| **1** | CH2 | 45.43 | 3.320 m<br>3.000 m |
| **2** | CH2 | 40.22 | 2.530 m<br>2.210 m |
| **3** | C | 51.21 | - |
| **4** | CH | 154.25 | 6.815 (d, J = 10.3) |
| **5** | CH | 130.25 | 6.135 (d, J = 10.4) |
| **6** | C | 197.57 | - |
| **7** | CH2 | 36.64 | 2.530 m<br>2.160 m |
| **8** | CH | 64.81 | 3.78 m |
| **1'** | C | 131.70 | - |
| **2'** | CH | 128.84 | 7.17 (d, J = 8.6) |
| **3'** | CH | 116.85 | 6.72 (d, J = 8.6) |
| **4'** | C | 158.14 | - |
| **5'** | CH | 116.82 | 6.72 (d, J = 8.6) |
| **6'** | CH | 128.77 | 7.17 (d, J = 8.6) |

**fig. S33. NMR assignment of 13 ($[M+H]^+$ = *m/z* 230.1176).** Assignments, as listed in the table above are based on the NMR spectra shown in figs. S34-S39. All experiments were recorded at 600 MHz in deuterated DMSO (DMSO-$d_6$) unless specified.

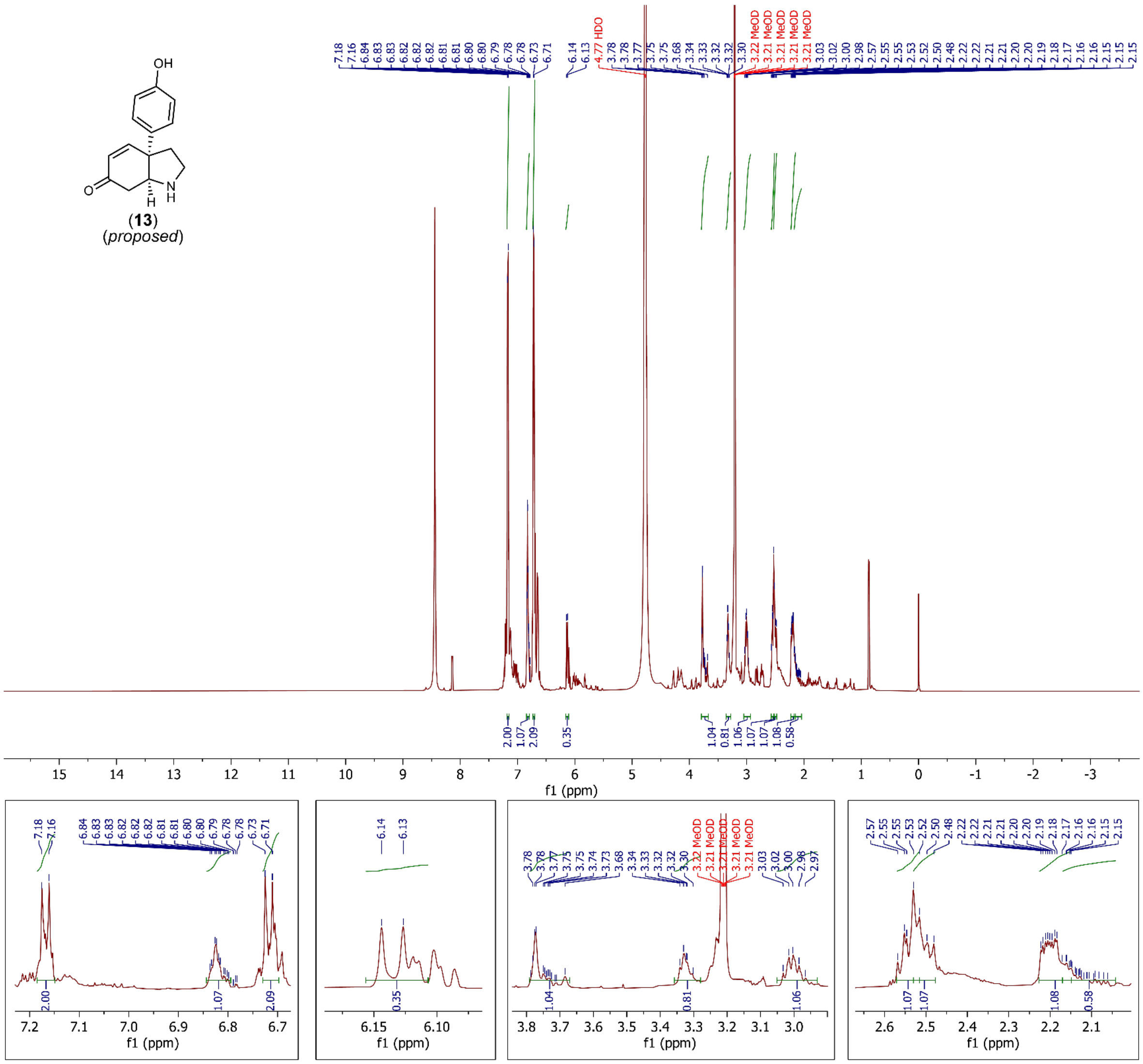


**fig. S34. $^1$H-NMR spectrum of 13 ([M+H]$^+$ = *m/z* 230.1176).** Ammonium formate salt contamination is detected at 8.4442 ppm.

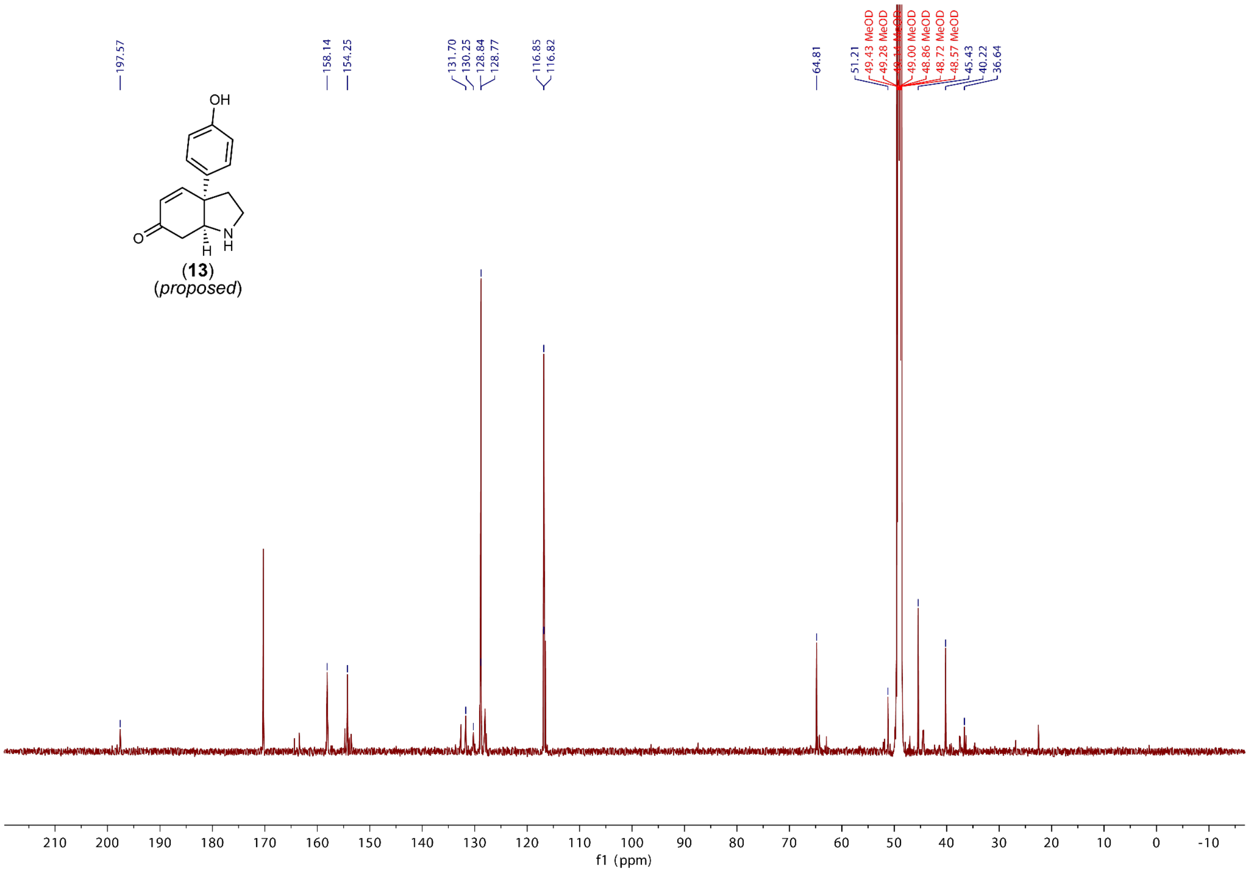


**fig. S35. $^{13}$C-NMR spectrum of 13 ($[M+H]^+$ = *m/z* 230.1176).** Ammonium formate salt contamination is detected at 170.2671 ppm.

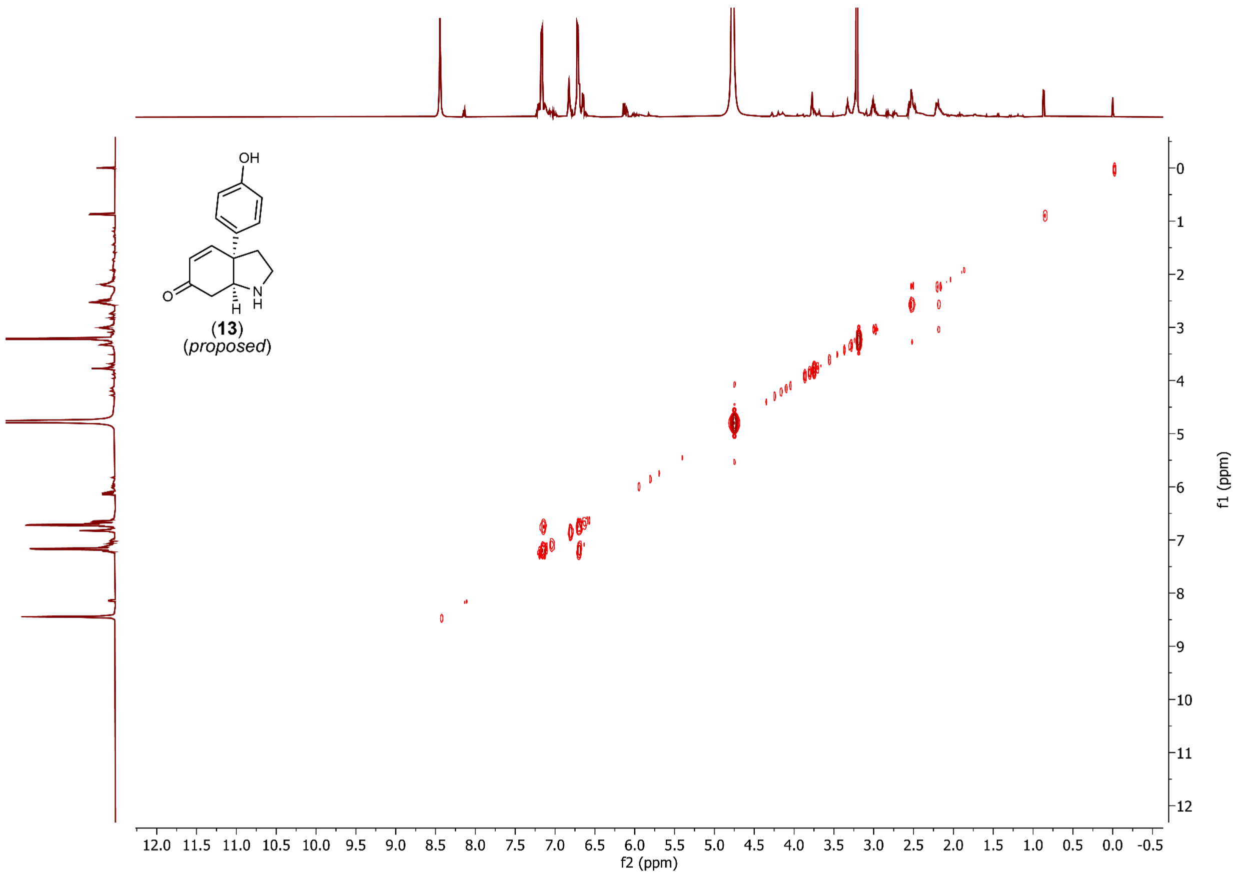


**fig. S36. COSY spectrum of 13 ($[M+H]^+$ = *m/z* 230.1176).**

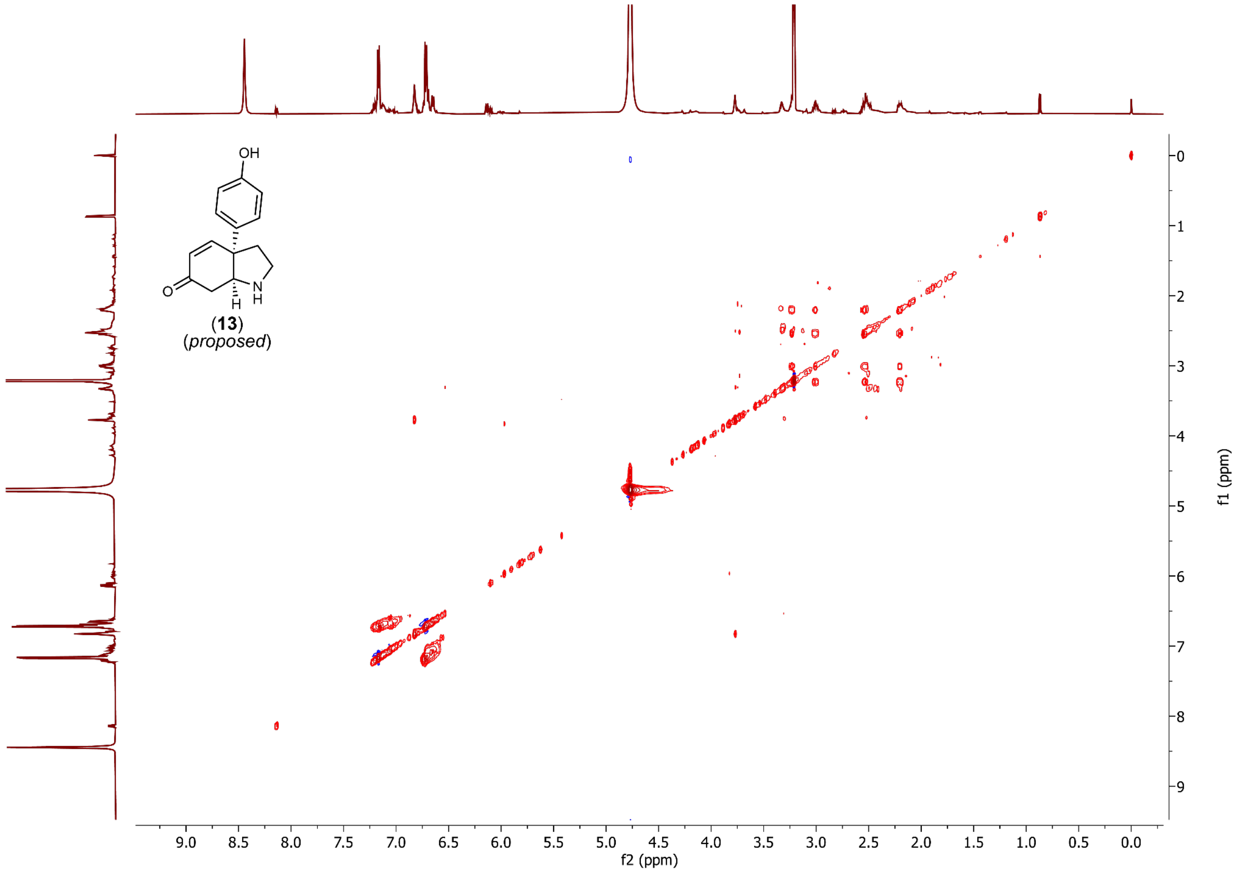


**fig. S37. TOCSY spectrum of 13 ([M+H]$^+$ = *m/z* 230.1176).**

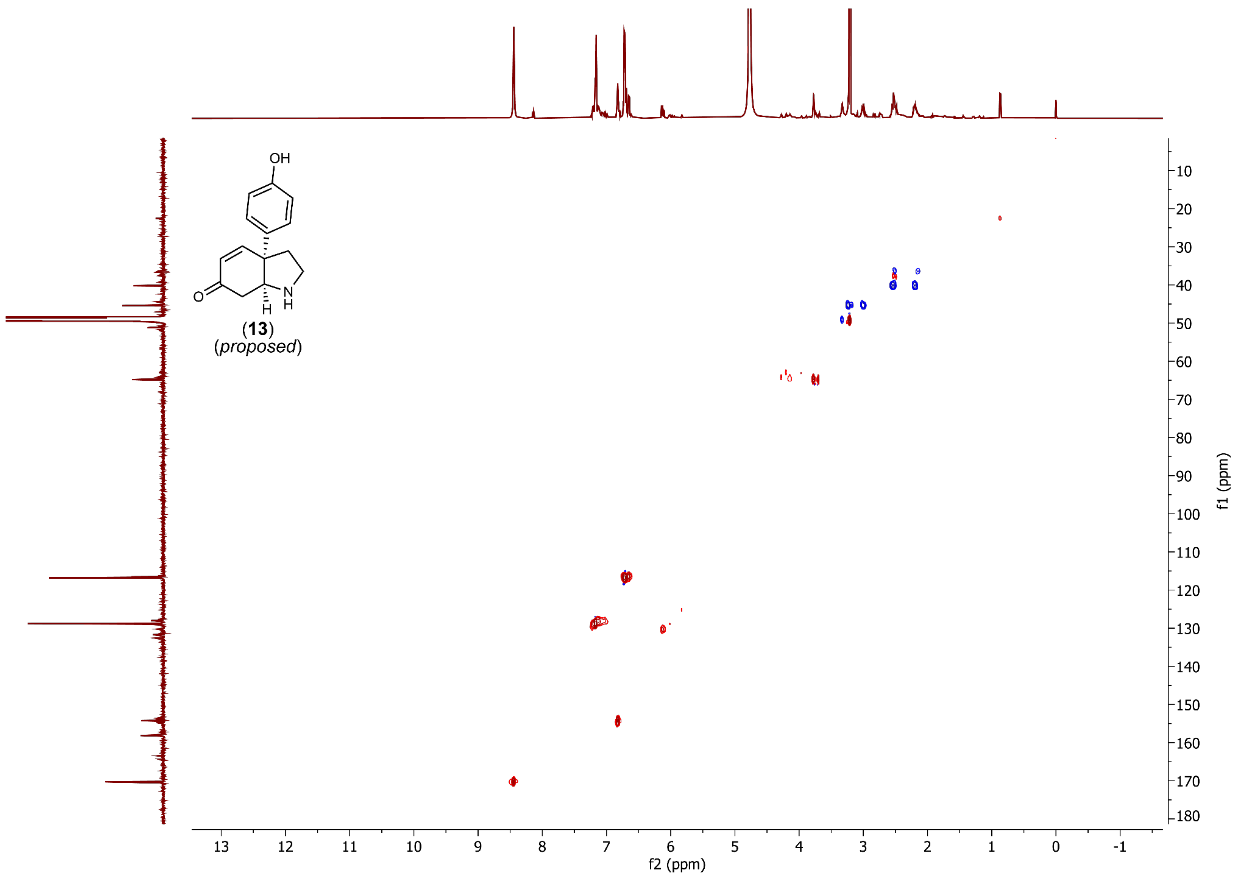


**fig. S38. HSQC spectrum of 13 ($[M+H]^+$ = *m/z* 230.1176).**

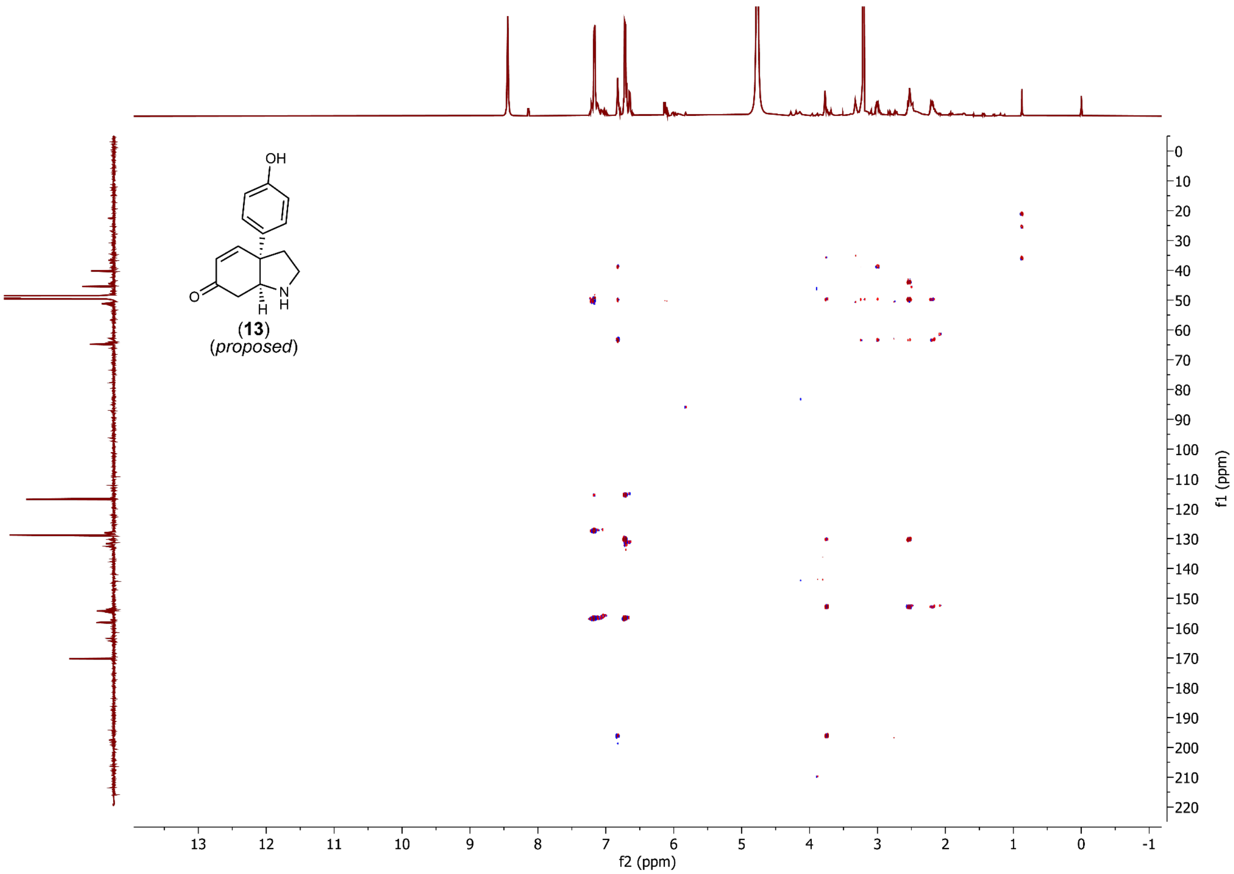


**fig. S39. HMBC spectrum of 13 ([M+H]$^{+}$ = *m/z* 230.1176).**

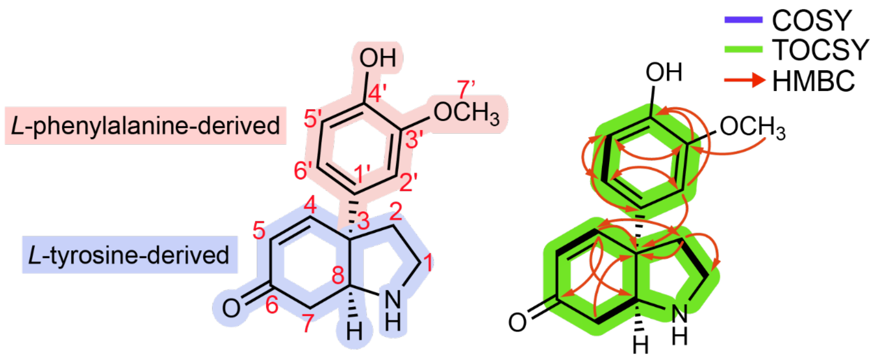


(**17**)
(*proposed*)

| carbon | | 13C δ (ppm) | 1H δ (ppm, J in Hz) |
|---|---|---|---|
| **1** | CH2 | 45.42 | 3.407 m<br>3.174 m |
| **2** | CH2 | 39.75 | 2.705 m<br>2.373 m |
| **3** | C | 51.42 | - |
| **4** | CH | 153.50 | 6.970 (J = 10.3) |
| **5** | CH | 130.79 | 6.277 (d, J = 10.3) |
| **6** | C | 197.80 | – |
| **7** | CH2 | 36.90 | 2.67 m<br>2.60 m |
| **8** | CH | 64.89 | 3.986 m |
| **1'** | C | 132.12 | - |
| **2'** | CH | 111.36 | 6.978 (d, J = ~2) |
| **3'** | C | 149.53 | - |
| **4'** | C | 147.64 | - |
| **5'** | CH | 116.66 | 6.825 (d, J = 8.3) |
| **6'** | CH | 120.49 | 6.879 (dd, J = 8.3, 2.2) |
| **7'** | OCH3 | 56.56 | 3.869 (s) |

**fig. S40. NMR assignment of 17 ($[M+H]^+$ = *m/z* 260.1281).** Assignments, as listed in the table above are based on the NMR spectra shown in figs. S41-S46. All experiments were recorded at 600 MHz in methanol-$d_4$.

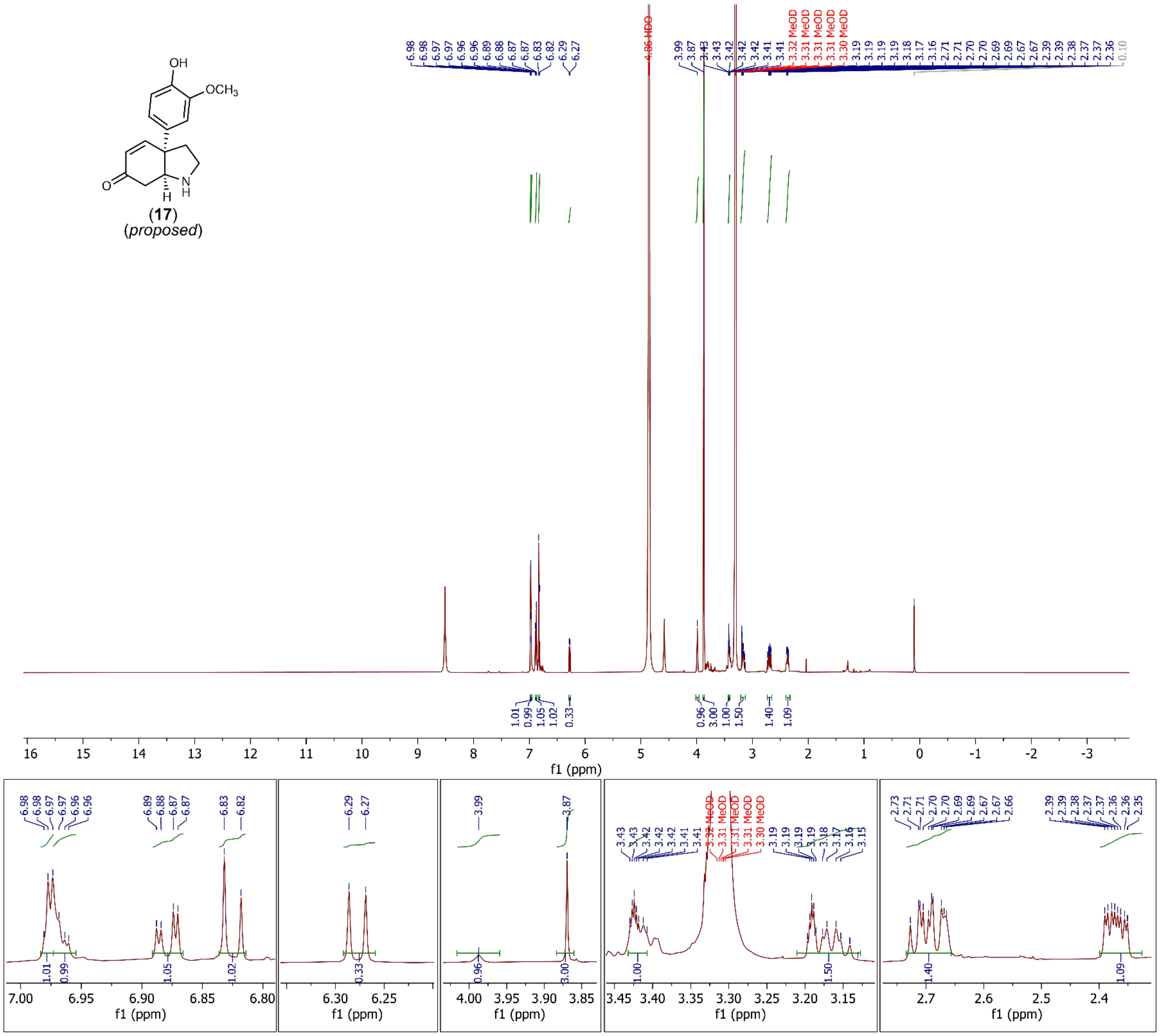


**fig. S41. $^{1}$H-NMR spectrum of 17 ($[M+H]^{+}$ = *m/z* 260.1281).** Ammonium formate salt contamination is detected at 8.5115 ppm.

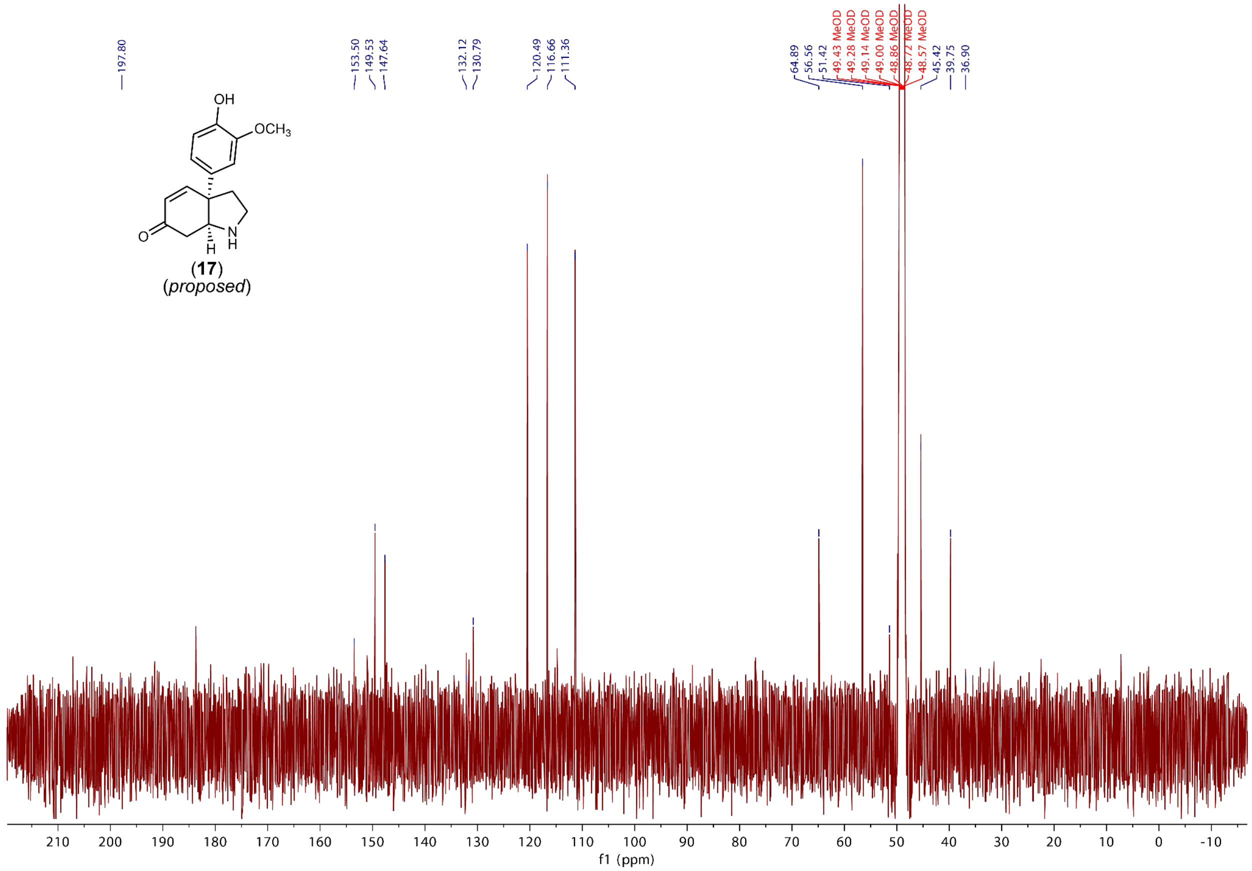


**fig. S42. $^{13}$C-NMR spectrum of 17 ([M+H]$^+$ = *m/z* 260.1281).** Carbon-6 signal was inferred from HMBC spectrum of **17** in fig. S46 and $^{13}$C spectrum of **13** in fig. S35.

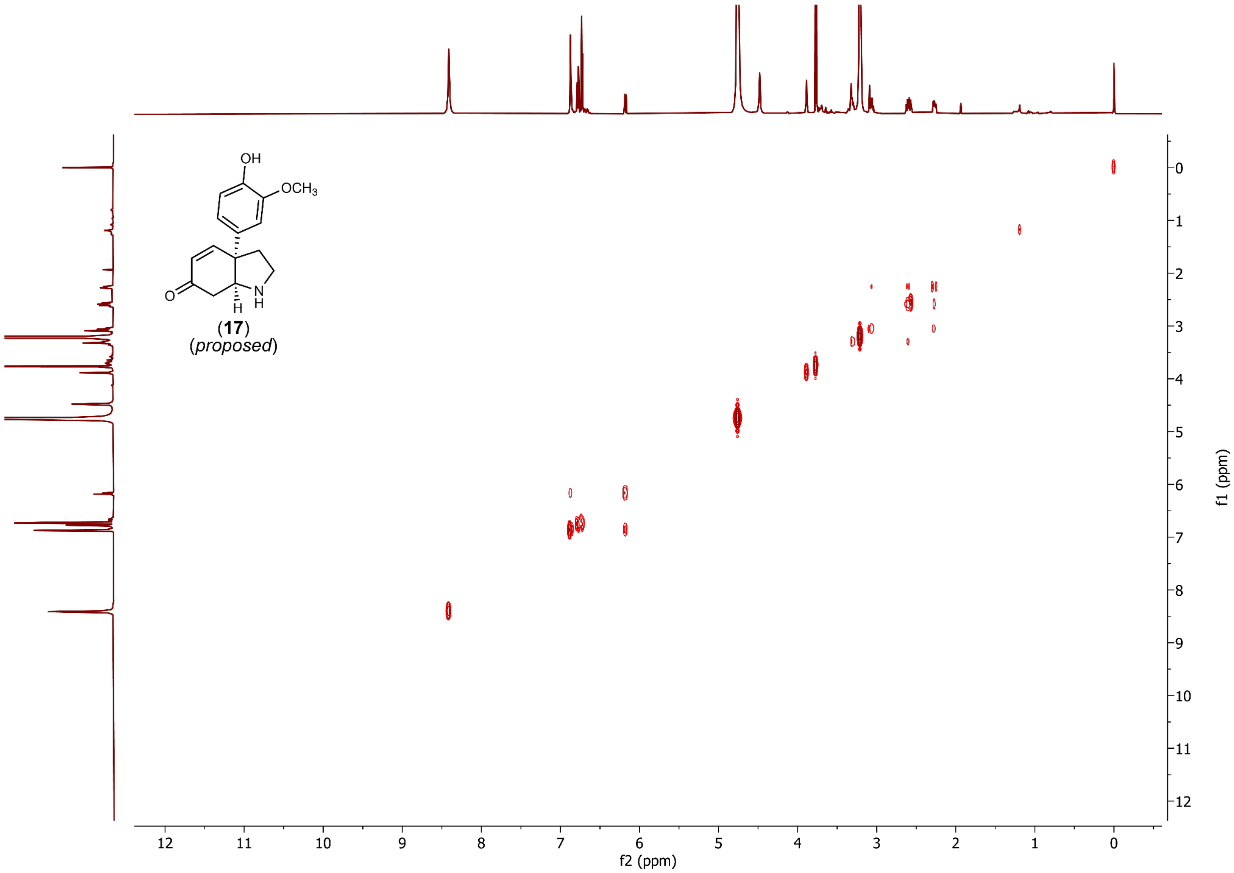


**fig. S43. COSY spectrum of 17 ([M+H]$^+$ = *m/z* 260.1281).**

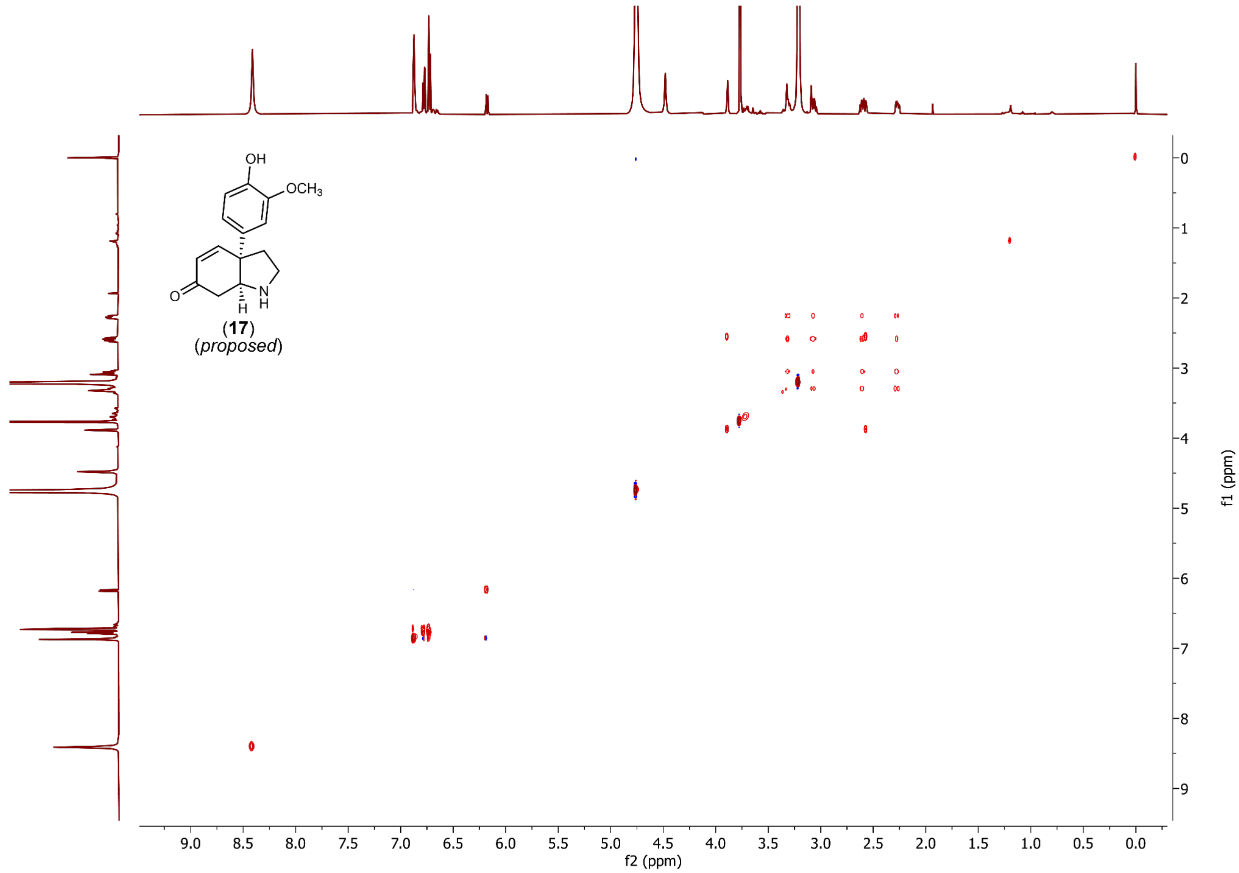


**fig. S44. TOCSY spectrum of 17 ([M+H]$^+$ = *m/z* 260.1281).**

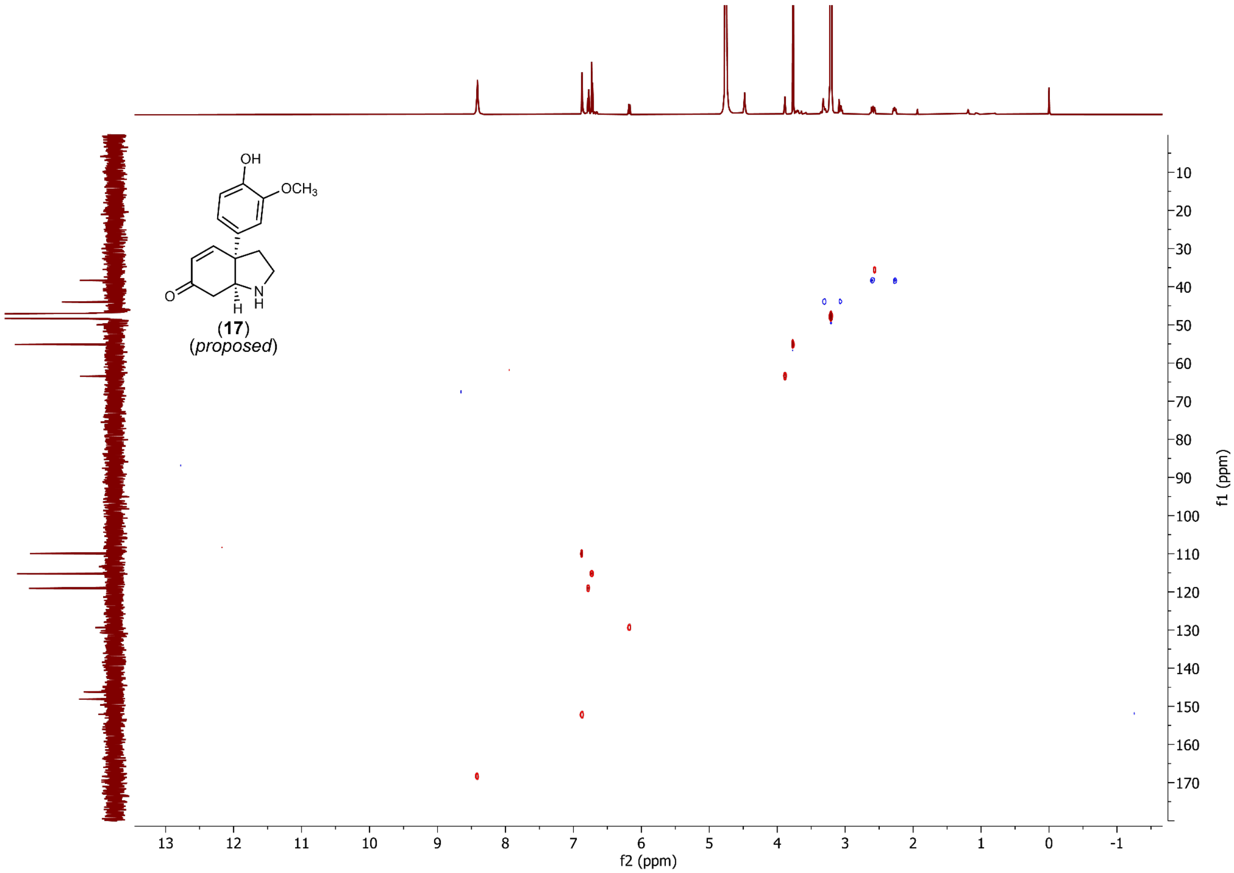


**fig. S45. HSQC spectrum of 17 ([M+H]$^+$ = *m/z* 260.1281).**

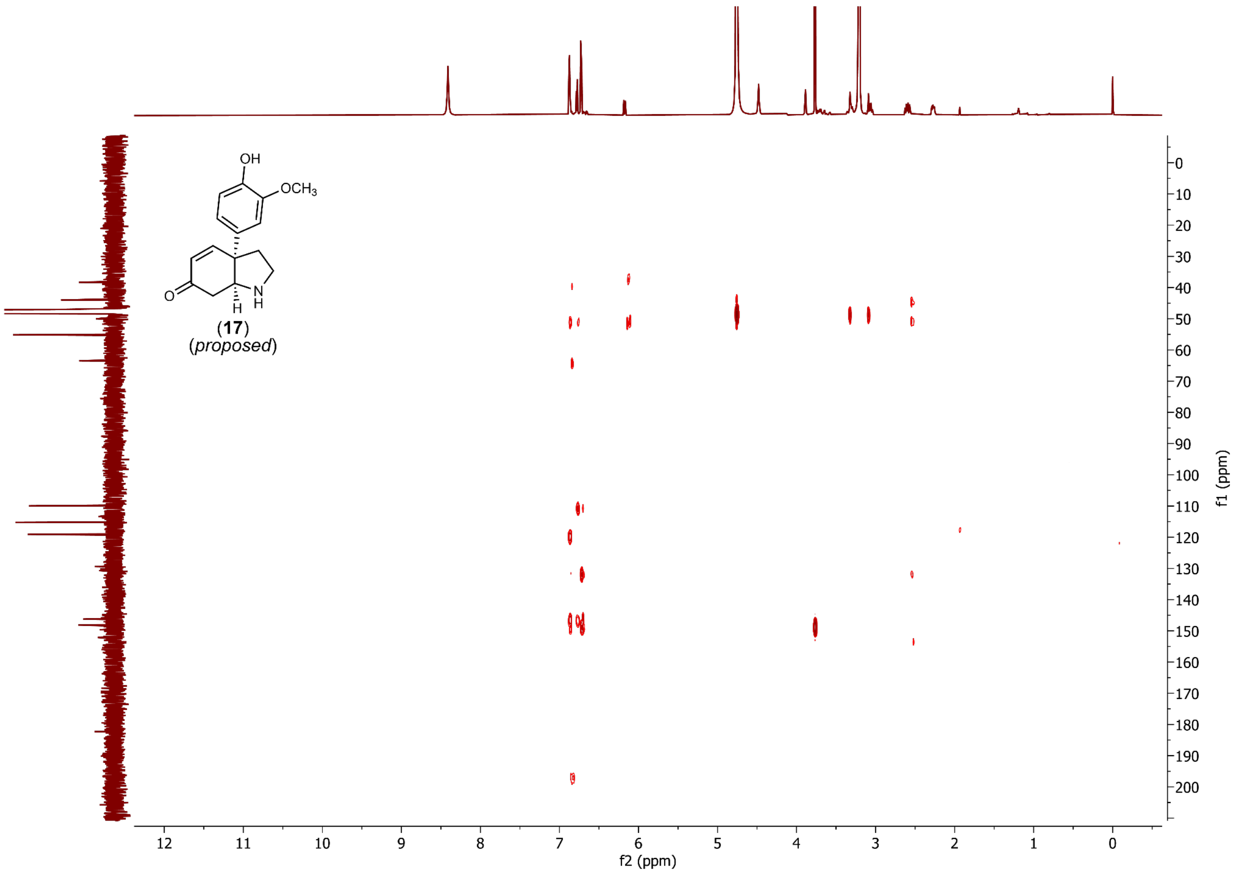


**fig. S46. HMBC spectrum of 17 ([M+H]$^+$ = *m/z* 260.1281).**

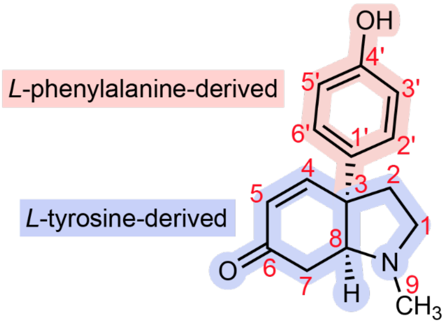


sceletenone (**14**)
(*proposed*)

| carbon | | 1H δ (ppm), 600 MHz CD3OD | multiplicity, J (Hz), integration | published, 100 MHz CDCl3 |
|---|---|---|---|---|
| **2', 6'** | CH | 7.24 | d, J = 8.7; 2.05 H (AA'BB') | 7.02 |
| **4** | CH | 6.88 | dd, J = 10.2, 1.9; 0.96 H | 6.74 |
| **3', 5'** | CH | 6.80 | d, J = 8.7; 2.00 H (AA'BB') | 7.02 |
| **5** | CH | 6.09 | d, J = 10.2; 0.93 H | 6.09 |
| **8** | CH | 3.20 | m; 1.00 H | 3.3 |
| **9** | CH3 | 2.43 | s; 3.00 H | 2.32 |
| **1 / 7** | CH2 | 2.93 | m; 0.99 H | 2.0-2.7 (6H) |
| **1 / 7** | CH2 | 2.71 | m; 1.06 H | 2.0-2.7 (6H) |
| **7 / 2** | CH2 | 2.57 | m; 2.04 H | 2.0-2.7 (6H) |
| **2** | CH2 | 2.22 | m; 1.01 H | 2.0-2.7 (6H) |
| **2** | CH2 | 2.04-2.09 | m; 0.64 H | 2.0-2.7 (6H) |

**fig. S47. NMR assignment of sceletenone (14) ($[M+H]^+$ = *m/z* 244.1332).** Assignments, as listed in the table below are based on the $^1$H-NMR shown in fig. S48 and comparison to previously published $^1$H-NMR spectrum(*22*).

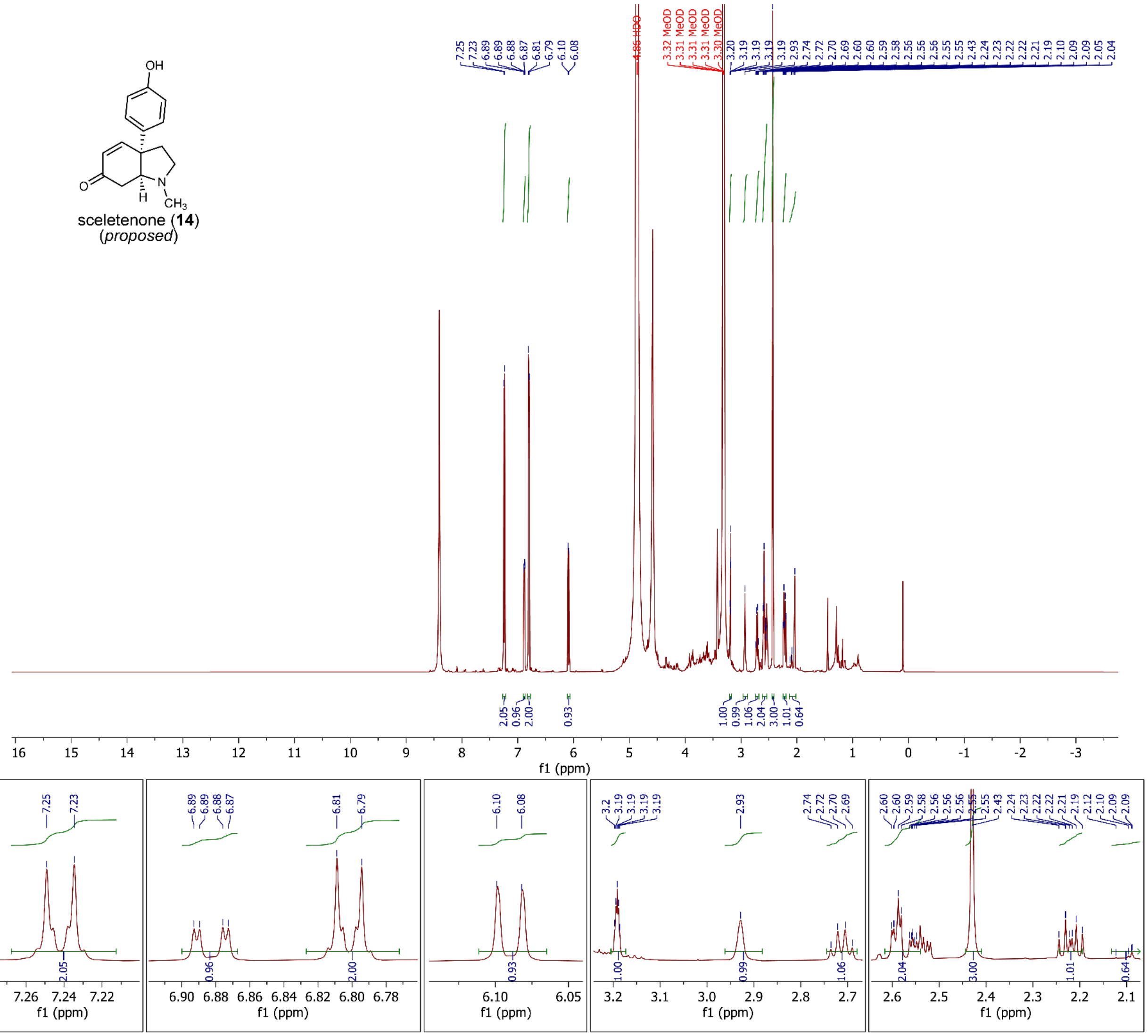


**fig. S48. $^1$H-NMR spectrum of sceletenone (14) ([M+H]$^+$ = *m/z* 244.1332).** Spectrum collected in methanol-$d_4$, 600 MHz, referenced to methanol-$d_3$ = 3.31 ppm. Spectrum showed an additional singlet (3.00 H) peak at 2.43 ppm, which is not present in the $^1$H-NMR spectrum of *N*-demethyl-sceletenone (**13**) shown in fig. S34. Ammonium formate salt contamination is detected at 8.4050 ppm.

### Supplementary Tables

**table S1. Source of plant genomes for comparative genomic analyses.** 13 taxa used for phylogenomic analysis in fig. S12A and for BLASTp sequence curation in the phylogenetic trees in Fig. 4G, figs. S7A, S12B.

| Plant Taxon | Common name | Reference |
|---|---|---|
| *Amaranthus tricolor* | amaranth | NCBI (GCA_026212465.1) |
| *Beta vulgaris* | beet | NCBI (GCF_026745355.1) |
| *Corchorus capsularis* | white jute | NCBI (GCA_001974805.1) |
| *Carpobrotus edulis* | highway ice plant | NCBI (GCA_965788445.1) |
| *Chenopodium quinoa* | quinoa | NCBI (GCF_001683475.1) |
| *Dianthus caryophyllus* | carnation | CNGBdb (CNA0069635) |
| *Hordeum vulgare* | barley | NCBI (GCF_904849725.1) |
| *Lycoris radiata* | red spider lily | CNGBdb (CNS1431320) |
| *Mesembryanthemum crystallinum* | ice plant | Perron *et al*. 2025(*69*) |
| *Mesembryanthemum tortuosum* | kanna | This study |
| *Simmondsia chinensis* | jojoba | CNCB (GWHAASQ00000000) |
| *Spinacia oleracea* | spinach | NCBI (GCF_020520425.1) |
| *Vitis vinifera* | grape | NCBI (GCF_030704535.1) |

### Supplementary Methods

Listed below are LC-MS methods and parameters used for data acquisition. During the course of this study, we predominantly used two HPLC methods for separating and analyzing metabolites discussed in this study. The first ("HILIC - 16 minute method") used an InfinityLab Poroshell 120 HILIC column (2.1 x 150 mm, 1.9 µm) with water with 10 mM ammonium formate and 0.1% formic acid (solvent A) and 90% acetonitrile, 10% water with 10 mM ammonium formate and 0.1% formic acid (solvent B) as the mobile phase solvents. This method was used for most analyses related to Kanna alkaloid biosynthesis in *M. tortuosum* and *N. benthamiana*. A second method ("C18 - 9 minute method") used a ZORBAX RRHD Eclipse Plus C18 column (2.1 × 50 mm, 1.8 µm) and consisted of water +0.1% (v/v) formic acid (solvent A) and acetonitrile +0.1% (v/v) formic acid (solvent B) as the mobile phase solvents. We used this method for activity assay for fractionation of KDR-1, *in vitro* enzyme assays (e.g. ATH, CYP72A1709), and additional analysis related to mesembrine biosynthesis.

<u>Mass spectrometry parameters:</u>

| **Instrument:** 6546 LC-MS (Agilent 1290 Infinity II UHPLC, Agilent 6546 Q-TOF) | |
|---|---|
| **Source** | Dual AJS ESI |
| **Mode** | ESI positive or negative MS |
| **Data type** | Centroid |
| **Drying gas temp** | 325 C |
| **Drying gas flow rate** | 10L/min |
| **Nebulizer** | 35 psig |
| **Sheath gas temp** | 350 C |
| **Sheath gas flow** | 12 L/min |
| **VCap** | 4000 V |
| **Fragmentor** | 135 V |
| **Skimmer** | 45 V |
| **OCT 1 RF Vpp** | 750 V |

HILIC - 16 minute method:

| Instrument | Agilent 1290 Infinity II UHPLC, Agilent 6546 Q-TOF |
|---|---|
| **Method** | HILIC - 16 minute method |
| **Column** | InfinityLab Poroshell 120 HILIC (2.1 x 150 mm, 1.9 µm) |
| **Solvent A** | water<br>with 10 mM ammonium formate and 0.1% formic acid |
| **Solvent B** | 90% acetonitrile / 10% water<br>with 10 mM ammonium formate and 0.1% formic acid |
| **Flow rate** | 0.4 mL/min |
| **Injection volume** | 1 µL |

| Time (min.) | A [%] | B [%] |
|---|---|---|
| 0.00 | 0.00 | 100.00 |
| 0.50 | 0.00 | 100.00 |
| 10.50 | 30.00 | 70.00 |
| 12.00 | 30.00 | 70.00 |
| 12.01 | 0.00 | 100.00 |
| 16.00 | 0.00 | 100.00 |

C18 - 9 minute method:

| Instrument | Agilent 1290 Infinity II UHPLC, Agilent 6546 Q-TOF |
|---|---|
| **Method** | C18 - 9 minute method |
| **Column** | ZORBAX RRHD Eclipse Plus C18 (2.1 × 50 mm, 1.8 µm) |
| **Solvent A** | water<br>with 0.1% formic acid |
| **Solvent B** | acetonitrile<br>with 0.1% formic acid |
| **Flow rate** | 0.6 mL/min |
| **Injection volume** | 1 µL |

| Time (min.) | A [%] | B [%] |
|---|---|---|
| 0.00 | 97.00 | 3.00 |
| 0.50 | 97.00 | 3.00 |
| 6.50 | 50.00 | 50.00 |
| 6.51 | 5.00 | 95.00 |
| 8.00 | 5.00 | 95.00 |
| 8.01 | 97.00 | 3.00 |
| 9.00 | 97.00 | 3.00 |

**Supplementary Code**

This file contains scripts used for data analysis and processing for this work. Certain code and related documents were generated using the assistance of AI tools (e.g. Codex, Claude Code).
